# FOXP1 regulates the development of excitatory synaptic inputs onto striatal neurons and induces phenotypic reversal with reinstatement

**DOI:** 10.1101/2023.10.23.563675

**Authors:** Nitin Khandelwal, Ashwinikumar Kulkarni, Newaz I. Ahmed, Mathew Harper, Genevieve Konopka, Jay Gibson

## Abstract

Long-range glutamatergic inputs from the cortex and thalamus are critical for motor and cognitive processing in the striatum. Transcription factors that orchestrate the development of these inputs are largely unknown. We investigated the role of a transcription factor and high-risk autism-associated gene, FOXP1, in the development of glutamatergic inputs onto spiny projection neurons (SPNs) in the striatum. We find that FOXP1 robustly drives the strengthening and maturation of glutamatergic input onto dopamine receptor 2-expressing SPNs (D2 SPNs) but has a comparatively milder effect on D1 SPNs. This process is cell-autonomous and is likely mediated through postnatal FOXP1 function at the postsynapse. We identified postsynaptic FOXP1-regulated transcripts as potential candidates for mediating these effects. Postnatal reinstatement of FOXP1 rescues electrophysiological deficits, reverses gene expression alterations resulting from embryonic deletion, and mitigates behavioral phenotypes. These results provide support for a possible therapeutic approach for individuals with FOXP1 syndrome.

## Introduction

Long-range excitatory glutamatergic afferents provide motor, sensory, and cognitive information into the striatum, where this information is integrated for learning and action planning1. These inputs originate primarily from the cortex and thalamus, and the mechanisms of their development and function have been studied ^2–4^ (and reviewed in ^5^). Transcription factors involved in striatal development have been widely studied in the context of early developmental processes, such as cell proliferation, differentiation, migration, and fate determination of striatal neurons ^6–9^. Surprisingly, the transcription factors and the genetic programming that orchestrate the construction of synaptic connectivity in striatal circuitry during development remain largely unexplored. Exceptions are autism-linked transcription factors, FOXP2 and MEF2C, which function together to regulate the formation of corticostriatal synapses ^10^. However, the functional relevance of transcription factors in evoked synaptic transmission has not been investigated so far. Furthermore, the striatum is composed of functionally distinct GABAergic spiny projection neurons, D1 and D2 SPNs, and our understanding of the regulation of cortical projections received by these SPN subtypes is limited.

Forkhead box P1 (FOXP1) is a key transcription factor that is highly enriched in the developing and mature striatum ^11^. *FOXP1* haploinsufficiency causes FOXP1 syndrome with symptoms that include Intellectual Disability (ID) and autism ^12,13^. Loss of function, heterozygous mutations are among the most significant recurrent de novo mutations associated with autism ^14–16^. FOXP1 exhibits a robust expression in SPNs and has been implicated in establishing gross striatal structure, behavior, fate determination, D1/D2 SPN physiology, and the regulation of essential genes in SPNs ^8, 17–19^. Unlike other transcription factors involved in early embryonic striatal development, FOXP1 expression begins later at E14.5 and is most strongly expressed in postmitotic neurons ^11, 20^ which is maintained through adulthood. The timing of FOXP1 expression suggests it may function in sculpting striatal circuitry during later developmental stages. While FOXP1 is expressed in both D1 and D2 SPNs, it plays a more consequential role in D2 SPNs ^8, 19^.

Previous investigations have reported *Foxp1*-dependent regulation of neuronal excitability and synaptic connections not only in the mouse cortex and hippocampus but also in the songbird brain ^18, 21, 22^. Our previous work demonstrated that FOXP1 is critical for maintaining normal intrinsic excitability of D2 SPNs by regulating the expression of leak and inwardly rectifying potassium channels ^19^. This was likely mediated by a later postnatal FOXP1 function since postnatal *Foxp1* deletion could recapitulate the hyperexcitability phenotype induced by embryonic deletion.

We wanted to determine if a similar D2 SPN locus and postnatal function of FOXP1 applies to synaptic regulation as well. It is unclear how the development of striatal circuits maintains a stable balance between the activity of D1 and D2 SPNs ^3,23^. Considering the preferential role of FOXP1 in the development of D2 SPNs, we hypothesize that FOXP1 plays a key role in regulating the synaptic balance of long-range glutamatergic inputs onto striatal neurons through cell-autonomous regulation of D2 SPNs. Finally, there are no reports describing pre vs. postnatal function of a transcription factor in sculpting striatal circuitry, which is crucial for designing successful therapeutics. Here, we hypothesize that FOXP1 functions postnatally to regulate glutamatergic synaptic inputs.

The extent and transcriptional mechanisms by which postnatal reinstatement of a transcription factor (after its absence during earlier developmental stages) can restore proper neuronal physiology is largely unexplored. A recent study has reported a rescue of EEG phenotype, behavior characteristics, and a few genes with postnatal reinstatement of *Tcf4* ^24^. However, to our knowledge, no study has used a large-scale, unbiased approach *in vivo* to more rigorously determine the extent of expression reversal and assess this at a cell-type-specific level. Moreover, it remains uncertain whether the rescue of neurophysiological phenotypes with reinstatement of a transcription factor is mediated by the reversal of transcript levels induced by deletion (“genetic” rescue) or is mediated by an entirely different set of genes.

To address these questions, we conducted cell-specific embryonic and postnatal *Foxp1* deletion, uncovering a vital role for *Foxp1* in the development of functional glutamatergic synaptic transmission onto SPNs. Our findings reveal a robust weakening of glutamatergic inputs specifically onto D2 SPNs highlighting the necessity of FOXP1 in strengthening these inputs. In contrast, minor changes were observed with *Foxp1* deletion in D1 SPNs. A decrease in evoked synaptic responses was also observed with postnatal *Foxp1* deletion in D2 SPNs and is consistent with postnatal *Foxp1*-mediated regulation of glutamatergic inputs. Most importantly, we achieved partial rescue of the synaptic transmission and complete rescue of intrinsic excitability defects in D2 SPNs through postnatal *Foxp1* reinstatement. Lastly, single-nuclei RNA sequencing (snRNA-seq) uncovered candidate genes for FOXP1-dependent regulation of glutamatergic inputs onto D2 SPNs. Remarkably, the majority of differentially expressed genes induced by postnatal *Foxp1* reinstatement exhibited rescued expression levels compared to their levels with embryonic *Foxp1* deletion. These findings provide a basis for a potential therapeutic strategy for individuals with FOXP1 syndrome.

## Results

### Foxp1 deletion in D2 SPNs weakens corticostriatal inputs onto D2 SPNs

We measured the functional strength of corticostriatal inputs by employing mCherry-tagged Channelrhodpsin-2 (ChR2) in callosal projecting corticostriatal axons and examined light-evoked EPSCs (excitatory postsynaptic currents) in SPNs (Fig. 1A). To do this, we expressed ChR2 through injecting AAV9 into one cortical hemisphere on postnatal day (P) 1 and conducted recordings in the striatum of the contralateral hemisphere at P14-P18. By studying this contralateral projection, we isolate the inputs provided by a single presynaptic, cortical neuron type – the layer 5 intratelencephalic pyramidal neuron^25^. We compared EPSCs in D2 SPNs between mice with and without D2 SPN-specific *Foxp1* deletion at E14-15 (referred to as D2 *Foxp1^cKO^* and D2 *Foxp1^CTL^* mice, respectively).

**Figure 1.**
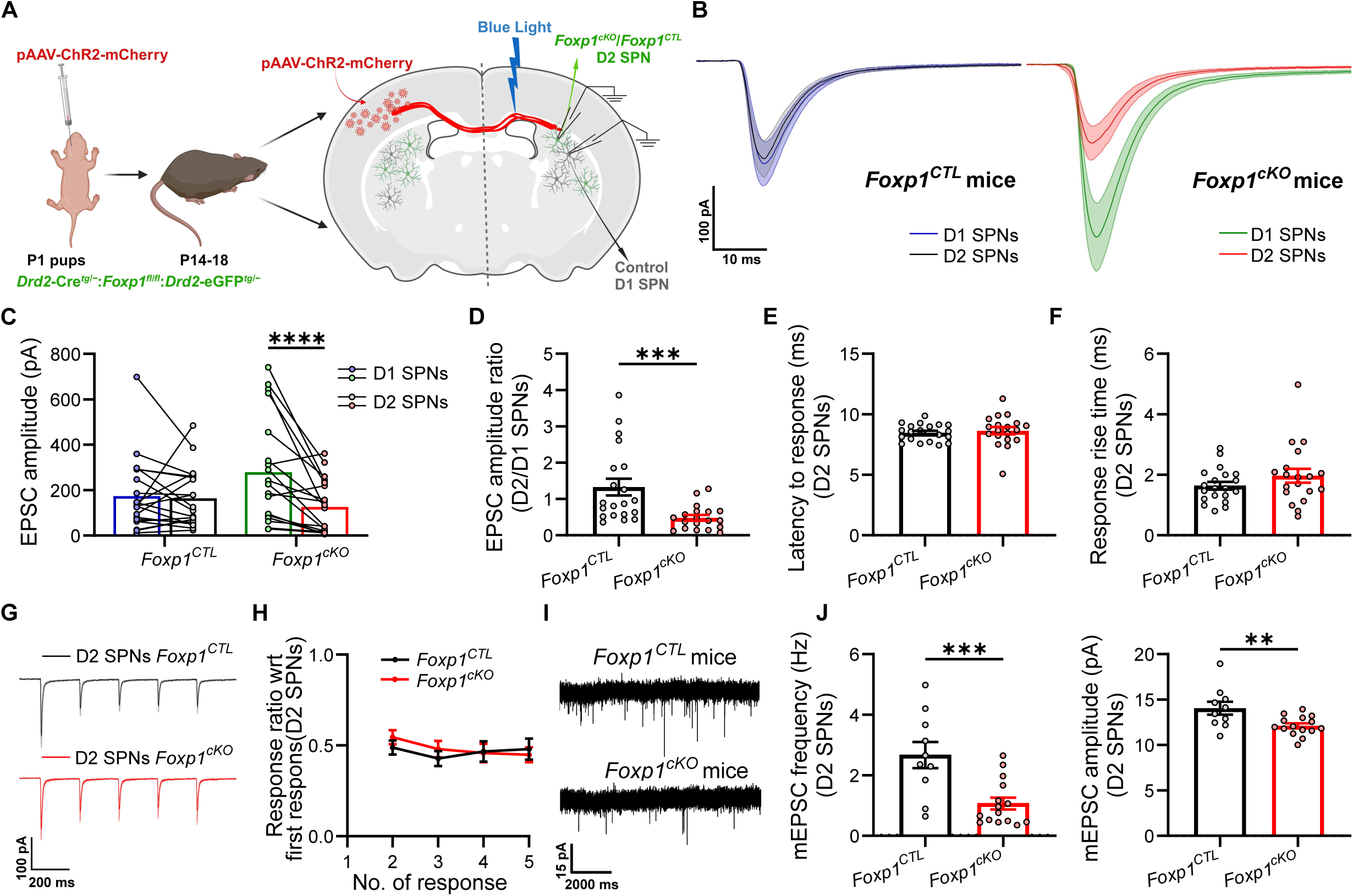
Excitatory cortical inputs onto D2 SPNs are decreased with D2-specific *Foxp1* deletion. **A.** pAAV-ChR2-mCherry virus was injected into the cortex of one hemisphere at P1, acute slices prepared at P14-18, and SPNs recorded in the contralateral striatum. **B.** Average (± SEM) EPSC responses triggered by blue light stimulation from D1 and D2 SPNs in D2 *Foxp1^CTL^*(blue and black, *left*) and in D2 *Foxp1^cKO^* mice (green and red, *right*). **C.** EPSC amplitudes in D1/D2 SPN pairs in D2 *Foxp1^CTL^* and D2 *Foxp1^cKO^* mice. **D.** D2/D1 ratio of EPSC amplitude for each D2/D1 pair in Fig. C. **E.** Latency to EPSC onset. **F.** Time taken by the EPSC response to reach the peak amplitude. **G.** Average (± SEM) EPSCs during a 5-pulse train of blue light in D2 SPNs. **H.** Ratio of 2^nd^-5^th^ response amplitude with respect to (wrt) first response amplitude shows no effect of *Foxp1* deletion on short-term dynamics of synaptic transmission in D2 SPNs. **I.** Example traces showing mEPSCs in D2 SPNs. **J.** Analysis of mEPSC measurements in both D2 *Foxp1^CTL^*and D2 *Foxp1^cKO^* mice show a reduced frequency (*left*) and amplitude (*right*) with embryonic *Foxp1* deletion in D2 SPNs. Ns in all figures are (D2 *Foxp1^CTL^*; D2 *Foxp1^cKO^*). For B-F. (20 D1/D2 pairs, 5 mice; 19 D1/D2 pairs, 4 mice). G, H. (18 D2 SPNs, 5 mice; 14 D2 SPNs, 4 mice). J. (10 D2 SPNs, 3 mice; 15 D2 SPNs, 3 mice). Statistics: C. 2-way ANOVA with Fisher’s LSD test. D. MW test. E, F, J. T-test. H. Mixed effects model with Hom Sidak corrections for multiple comparisons. ** p<0.01, *** p<0.001, **** p<0.001.

For every D2 SPN examined, we recorded from a neighboring, control D1 SPN to control for variability inherent in the stimulation method (*see methods for details*). EPSC amplitudes of paired D1 and D2 SPNs were nearly identical in D2 *Foxp1^CTL^* mice (Fig. 1B, C) as previously reported ^26^. However, in D2 *Foxp1^cKO^* mice, EPSC amplitude in D2 SPNs was roughly half of that observed in neighboring D1 SPNs (Fig. 1B, C), leading to a diminished EPSC amplitude ratio (D2/D1) (Fig. 1D). Although we observed a trend of increased amplitude in D1 SPNs in D2 *Foxp1^cKO^* mice as compared to D2 *Foxp1^CTL^* mice (p=0.0532), we believe this “absolute” measurement is unreliable due to stimulation variability (*see Methods*). We also measured EPSC rising slope in an earlier response time window to decrease contamination from disynaptic inhibition and observed a similar decline in D2/D1 ratio (Fig. 2A). No changes were observed in EPSC onset latency and rise time, suggesting unaltered action potential conduction in axons and AMPAR subunit composition (Fig. 1E, F). As reported previously, D2 SPNs displayed increased input resistance and intrinsic excitability in D2 *Foxp1^cKO^* mice (Fig. 2B, C). None of these changes were observed in the D1 SPNs (Fig. 2C-F). In summary, FOXP1 promotes the strength of corticostriatal glutamatergic inputs onto D2 SPNs relative to D1 SPNs.

**Figure 2.**
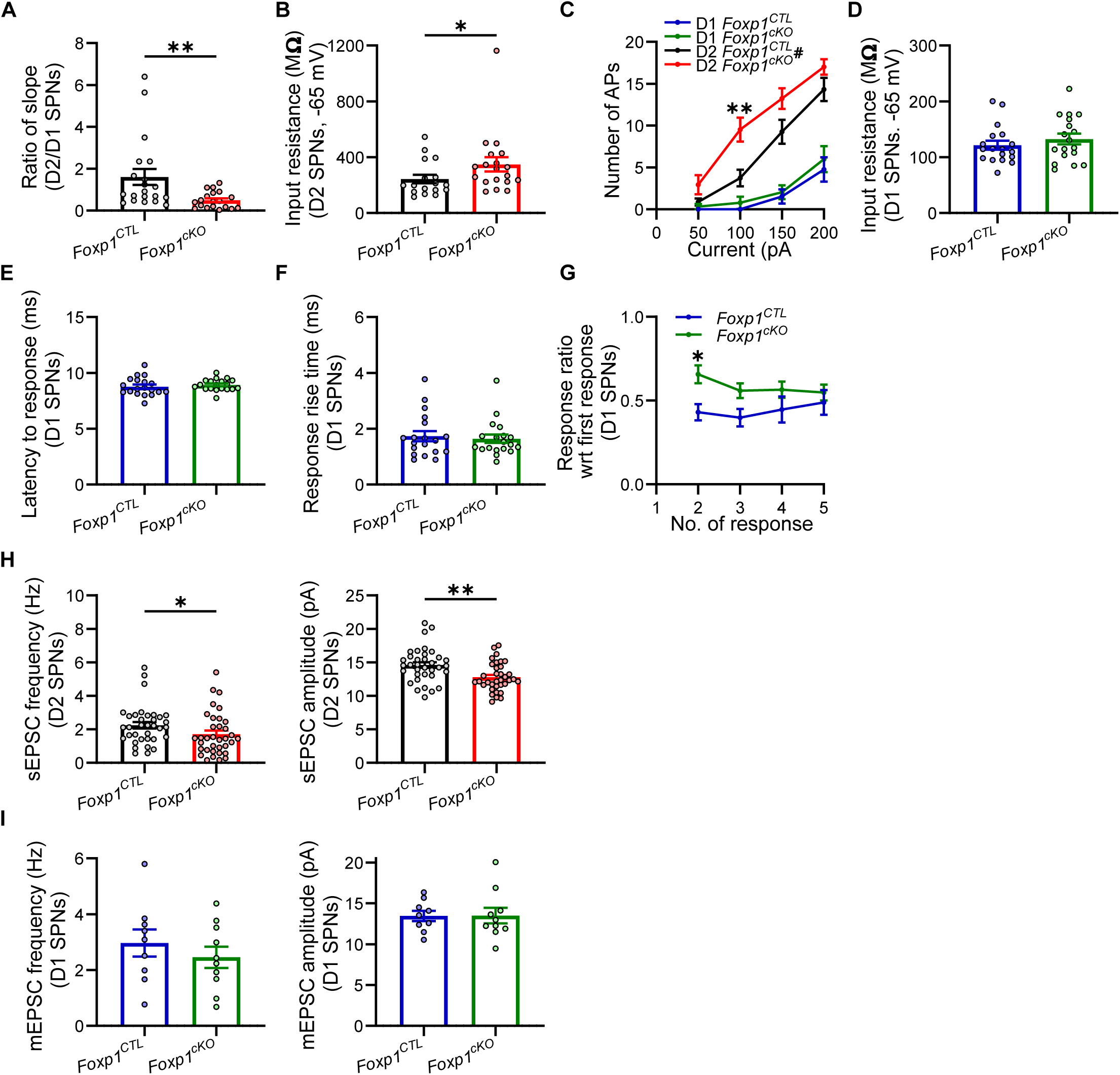
Excitatory cortical inputs onto D2 and D1 SPNs in D2 *Foxp1^cKO^* mice. **A.** D2/D1 ratio of the EPSC initial slope (line connecting 20% and 80% of EPSC peak). **B.** D2 SPNs show increased input resistance with embryonic *Foxp1* deletion in D2 *Foxp1^cKO^*mice. **C.** D2 SPNs with the loss of *Foxp1* fired more action potentials with the same amount of current injection while no changes were observed for D1 SPNs. **D.** D1 SPNs do not show changes in input resistance with *Foxp1* deletion in D2 SPNs. **E, F.** Latency to EPSC response (*left*) and response rise time (*right*) do not change in D1 SPNs in D2 *Foxp1^cKO^* mice. **G.** Ratio of 2^nd^-5^th^ response amplitude with respect to (wrt) the first response amplitude suggests decreased release probability in D1 SPNs with *Foxp1* deletion in D2 SPNs. **H.** Similar to mEPSCs, sEPSCs demonstrated reduced frequency (*left*) and amplitude (*right*) in D2 SPNs with *Foxp1* deletion. **I.** mEPSC frequency (*left*) and amplitude (*right*) of D1 SPNs did not change in D2 *Foxp1^cKO^* mice. Ns (D2 *Foxp1^CTL^*; D2 *Foxp1^cKO^*): For A. (20 D1/D2 pairs, 5 mice; 19 D1/D2 pairs, 4 mice). B, C. (17 D2 SPNs, 5 mice; 19 D2 SPNs, 4 mice). C, D. (18 D1 SPNs, 5 mice; 18 D1 SPNs, 4 mice). E, F. (19 D1 SPNs, 5 mice; 19 D1 SPNs, 4 mice). G. (16 D1 SPNs, 5 mice; 18 D1 SPNs, 4 mice). H. (35 D2 SPNs, 5 mice; 35 D2 SPNs, 9 mice). I. (9 D1 SPNs, 3 mice; 10 D1 SPNs, 3 mice). Statistics: A, B, F, H (*left*). MW test. C, G. Mixed effects model with Hom Sidak corrections for multiple comparisons. D, E, H (*right*), I, J. T-test. * p<0.05, ** p<0.01. # shows the genotypic difference in C.

To determine if weak synaptic function in D2 *Foxp1^cKO^*SPNs correlated with changes in presynaptic release probability, we recorded D2 SPNs while applying a train of light stimuli and measured the EPSC amplitude change during the train (Fig. 1G). The EPSC amplitudes (normalized to the first EPSC) evoked during these trains were not different in D2 SPNs between D2 *Foxp1^cKO^*and D2 *Foxp1^CTL^* mice, suggesting no changes in the presynaptic release probability (Fig. 1H). We also observed a non-cell-autonomous effect in D1 SPNs where normalized EPSC amplitude ratio of 2^nd^ – 5^th^ response was higher in D2 *Foxp1^cKO^*mice, suggesting reduced release probability (Fig. 2G).

To measure absolute changes in total glutamatergic input onto D2 SPNs with *Foxp1* deletion, we blocked action potential firing using TTX (1 µM) and measured miniature EPSCs (mEPSCs). Though both cortical and thalamic inputs likely contribute to mEPSC measurements ^1^, we considered the corticostriatal evoked response alterations robust enough to be reflected in mEPSC changes (*see Discussion*). We found a decrease in mEPSC frequency and amplitude in D2 SPNs in D2 *Foxp1^cKO^* mice (Fig. 1I, J). Similar results were observed for spontaneous EPSCs (sEPSCs) measurements in the absence of TTX (Fig. 2H). Since we did not observe changes in release probability, the decreased mEPSC frequency is consistent with fewer excitatory synapses while reduced amplitude suggests weaker individual synapses in D2 SPNs with the loss of *Foxp1*. Therefore, in addition to a shift in the balance of corticostriatal inputs onto D2 and D1 SPNs, *Foxp1* deletion reduces the absolute level of glutamatergic inputs onto D2 SPNs. D1 SPNs in D2 *Foxp1^cKO^*mice did not show any mEPSC changes (Fig. 2I). In summary, our findings indicate that FOXP1 is essential for the strengthening and maintenance of corticostriatal inputs onto D2 SPNs during development.

### Foxp1 deletion weakens both AMPAR and NMDAR mediated input but increases AMPAR/NMDAR synaptic composition

FOXP1 facilitates a social experience-dependent increase in the AMPAR/NMDAR composition ratio at synapses in songbirds ^22^, and such changes would likely impact synaptic plasticity, strengthening and information transfer ^27^. We investigated if a similar regulatory mechanism operates at corticostriatal synapses. We employed a protocol that effectively eliminates disynaptic inhibitory postsynaptic current (IPSC) contamination, allowing us to isolate NMDAR-mediated currents. By blocking GABAergic transmission with picrotoxin (0.1 mM), action potentials with TTX (1 µM) and potassium channels by 4-AP, light induced activation of ChR2 in corticostriatal presynaptic terminals resulted in isolated, action potential-independent, monosynaptic EPSCs ^28^. As observed above, the balance of corticostriatal excitation relative to D1 SPNs was decreased in D2 SPNs for AMPAR-mediated transmission in D2 *Foxp1^cKO^* mice (Fig. 3A, B). Interestingly, NMDAR-mediated EPSCs displayed a similar pattern with decreased amplitudes in D2 SPNs in D2 *Foxp1^cKO^* mice (Fig. 3C). Consequently, both AMPAR- and NMDAR-mediated EPSCs exhibited a reduced D2/D1 amplitude ratio (Fig. 3B, C). Moreover, D2 SPNs showed a decreased ratio of AMPAR/NMDAR-mediated EPSC amplitude in D2 *Foxp1^cKO^*mice indicating an increased relative contribution of NMDARs to corticostriatal inputs onto D2 SPNs (Fig. 3D). Higher NMDAR contribution is a signature of more immature excitatory synapses in SPNs ^29^. Importantly, these changes were specific to loss of *Foxp1* in D2 SPNs, as AMPAR/NMDAR ratio did not change for unmanipulated D1 SPNs in these mice (Fig. 3E).

**Figure 3.**
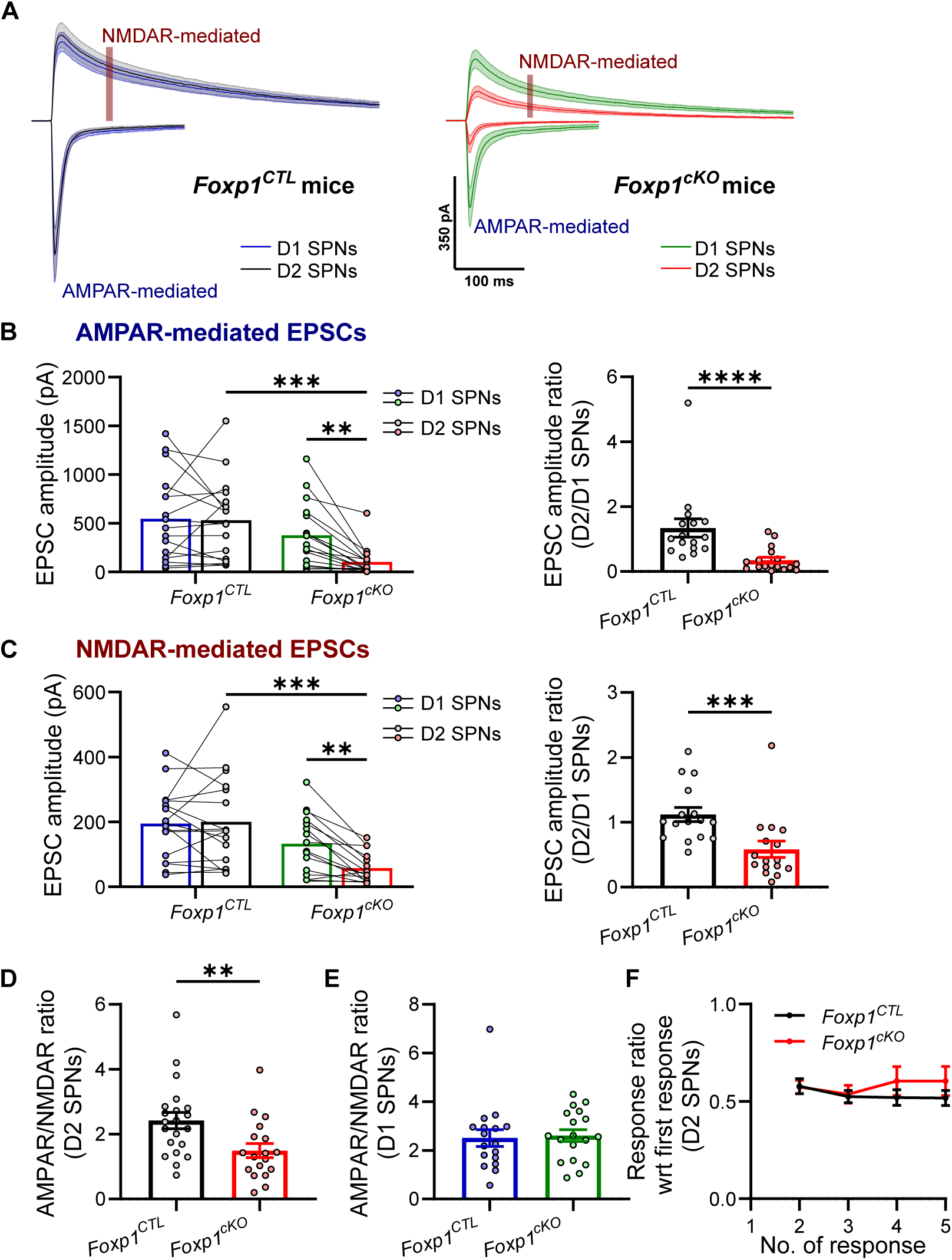
NMDAR-mediated component of evoked excitatory cortical inputs onto D2 SPNs is altered with D2-specific *Foxp1* deletion. **A.** Monosynaptic EPSCs were evoked by depolarization of presynaptic terminals during blockade of voltage-dependent Na^+^ channels. Average EPSC responses (± SEM) from D1 and D2 SPNs in slices from D2 *Foxp1^CTL^*mice (blue and black respectively, *left*) and from D2 *Foxp1^cKO^* mice (green and red respectively, *right*). AMPAR-mediated EPSC amplitude is measured as a response peak at -70 mV holding potential whereas, for NMDAR-mediated EPSC amplitude, it is the average response in a 90-100 ms window while holding at +50 mV (highlighted with a brown box). **B.** The amplitude of AMPAR-mediated EPSCs for each pair of D1/D2 SPNs in D2 *Foxp1^CTL^* and D2 *Foxp1^cKO^*mice (*left*). D2/D1 ratio of the EPSC amplitude for each pair (*right*). **C.** Amplitude of NMDAR-mediated EPSCs and D2/D1 EPSC amplitude ratio. **D, E.** The ratio of AMPAR- to NMDAR-mediated EPSC amplitudes for D2 and D1 SPNs in D2 *Foxp1^CTL^* and D2 *Foxp1^cKO^*mice. **F.** AMPAR-mediated EPSC amplitude measured during a stimulus train shows no change in short-term dynamics of synaptic transmission in D2 SPNs with *Foxp1* deletion. Ns in all figures are (D2 *Foxp1*^CTL^; D2 *Foxp1*^cKO^). For A-C. (16 D1/D2 pairs, 5 mice; 16 D1/D2 pairs, 3 mice). D. (21 D2 SPNs, 5 mice; 18 D2 SPNs, 3 mice). E. (17 D1 SPNs, 5 mice; 18 D1 SPNs, 3 mice). F. (18 D2 SPNs, 5 mice; 14 D2 SPNs, 3 mice). Statistics: B (*left*), C (*left*). 2-way ANOVA with Fisher’s LSD test. B (*right*), C (*right*). MW-test. D, E. T-test. F. Mixed effects model with Hom Sidak correction for multiple comparisons. ** p<0.01, *** p<0.001, **** p<0.001.

There was no change in the short-term dynamics of EPSC amplitude with *Foxp1* deletion in D2 SPNs under these conditions, again suggesting an absence of changes in presynaptic release probability (Fig. 3F). Furthermore, mEPSC frequency and amplitude were again decreased in D2 SPNs (Fig. 4A, B). In summary, *Foxp1* function is necessary for the strengthening and maturation of corticostriatal synapses, specifically in facilitating the inclusion of AMPARs in synapses onto D2 SPNs.

**Figure 4.**
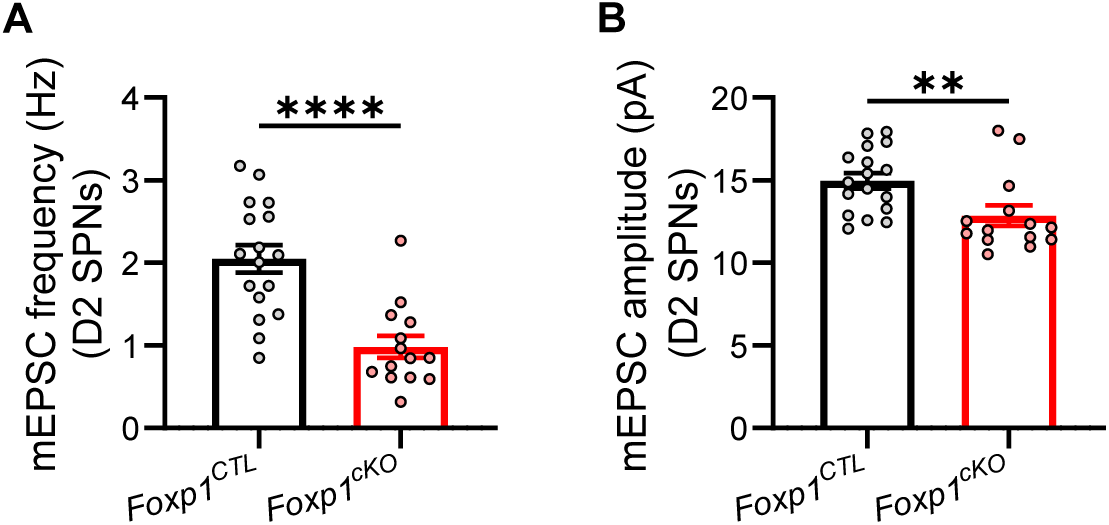
mEPSCs with *Foxp1* deletion in D2 SPNs. **A, B.** Decrease in the frequency and amplitude of mEPSCs in D2 SPNs with *Foxp1* deletion in modified ACSF used to record NMDAR-mediated EPSCs. Ns in all figures are (D2 *Foxp1^CTL^*; D2 *Foxp1^cKO^*). For A, B. (17 D2 SPNs, 3 mice; 14 D2 SPNs, 3 mice). Statistics: A, B. T-test. ** p<0.01, **** p<0.001.

### No detectable changes in germline heterozygous Foxp1 mouse

FOXP1 syndrome arises from heterozygous mutations leading to FOXP*1* haploinsufficiency ^12, 30^. The *Foxp1^Het^* mouse serves as a patient-relevant model, displaying increased D2 SPN excitability and some behavioral deficits ^17^. To determine whether the weakening of corticostriatal input to D2 SPNs also occurs in *Foxp1^Het^* mouse, we again employed blue light to stimulate ChR2-containing corticostriatal afferents and examined action potential-dependent synaptic responses in SPNs. The EPSC amplitude was comparable between D2 and D1 SPNs in both *Foxp1^WT^* and *Foxp1^Het^*mice, resulting in no significant difference in the EPSC amplitude ratio (D2/D1) (Fig. 5A, B). There were no changes in short-term dynamics in response to train stimuli in both D2 and D1 SPNs (Fig. 5C, D).

**Figure 5.**
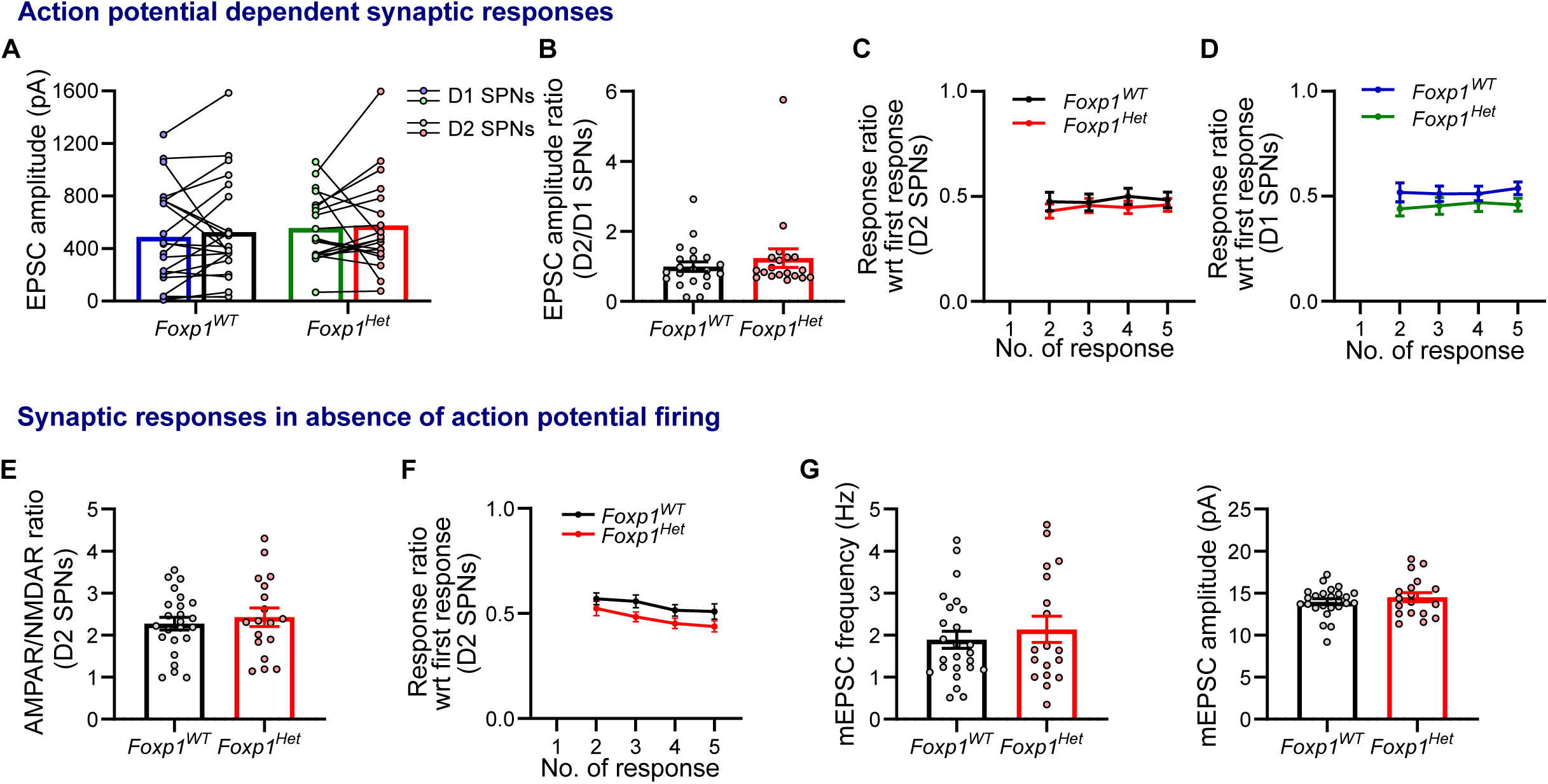
No changes were observed in excitatory cortical inputs in germline *Foxp1* heterozygous (*Foxp1^Het^*) mice. *Action potential dependent evoked EPSCs (A-D).* **A.** EPSC amplitude is comparable for D1/D2 SPN pairs in *Foxp1^WT^* (blue and black respectively, *left*) and *Foxp1^Het^* mice (green and red respectively, *right*). **B.** D2/D1 EPSC amplitude ratio for each D1/D2 pair. **C, D.** EPSC amplitude ratio for 2^nd^ – 5^th^ responses during a 5-pulse stimulus train did not change in both D2 and D1 SPNs in *Foxp1^Het^* mice (normalized to the first EPSC in the train). *Synaptic responses in the absence of action potential firing (E-G).* **E.** AMPAR/NMDAR ratio in D2 SPNs was similar in *Foxp1^WT^* and *Foxp1^Het^* mice. **F.** EPSC amplitude ratio of 2^nd^ – 5^th^ responses did not change in D2 SPNs in the absence of action potential firing in *Foxp1^Het^* mice. **G.** Both frequency (*left*) and amplitude (*right*) of mEPSCs in D2 SPNs did not change with heterozygous deletion of *Foxp1.* Ns in all figures are (*Foxp1^WT^*; *Foxp1^Het^*). For A, B. (21 D1/D2 pairs, 6 mice; 19 D1/D2 pairs, 5 mice). C. (20 D2 SPNs, 6 mice; 19 D2 SPNs, 5 mice). D. (22 D1 SPNs, 6 mice; 17 D1 SPNs, 5 mice). E. (24 D2 SPNs, 4 mice; 18 D2 SPNs, 3 mice). F. (23 D2 SPNs, 4 mice; 17 D2 SPNs, 3 mice). G. (25 D2 SPNs, 5 mice; 18 D2 SPNs, 5 mice). Statistics: A. 2-way ANOVA with Fisher’s LSD test. B, E. T-test. C, D, F. Mixed effects model with Hom Sidak correction for multiple comparisons. G. MW-test.

It is important to note that the D1 SPNs used for normalization in *Foxp1^Het^* mice also exhibit heterozygous *Foxp1* deletion, and therefore may not be ideal controls for the paired recordings. Consequently, we measured the AMPAR/NMDAR ratio in D2 SPNs, which is independent of D1 SPNs. However, we did not observe any differences (Fig. 5E). Responses to train stimuli and mEPSCs remained unaffected *in* these conditions (Fig. 5F, G). In summary, these results indicate that *Foxp1* expression levels in *Foxp1^Het^* mice are adequate for the normal development of corticostriatal inputs.

### Corticostriatal input onto D1 SPNs is unchanged with Foxp1 deletion in D1 SPNs

*Foxp1* deletion in D1 SPNs induces changes in the cellular and electrophysiological properties of D1 SPNs and causes behavioral defects ^8, 19^. We examined action potential-dependent corticostriatal inputs onto D1 SPNs in D1 *Foxp1^cKO^* mice. Recordings were conducted in D1/D2 pairs. In *Foxp1^CTL^*mice, there was no significant difference in the amplitude of AMPAR-mediated EPSCs between D1 and D2 SPNs (Fig. 6A, B). In contrast, D1 *Foxp1^cKO^*mice exhibited a significant decrease in the EPSC amplitude of D1 SPNs relative to their neighboring D2 SPNs (Fig. 6A, B). This effect, however, was not as pronounced as the changes observed in D2 SPNs in D2 *Foxp1^cKO^* mice, considering both the smaller magnitude difference and the lack of changes in the D1/D2 EPSC amplitude ratio in D1 *Foxp1^cKO^* mice (Fig. 6C). Moreover, *Foxp1* deletion did not affect the frequency and amplitude of mEPSCs in D1 SPNs (Fig. 6D-E). These data suggest that FOXP1 does not play a substantial role in regulating the development of corticostriatal inputs onto D1 SPNs. Interestingly, non-cell autonomous effects were observed on D2 SPNs in D1 *Foxp1^cKO^* mice with D2 SPNs displaying increased sEPSC frequency, reduced intrinsic excitability, and decreased input resistance (Fig. 7A-C).

**Figure 6.**
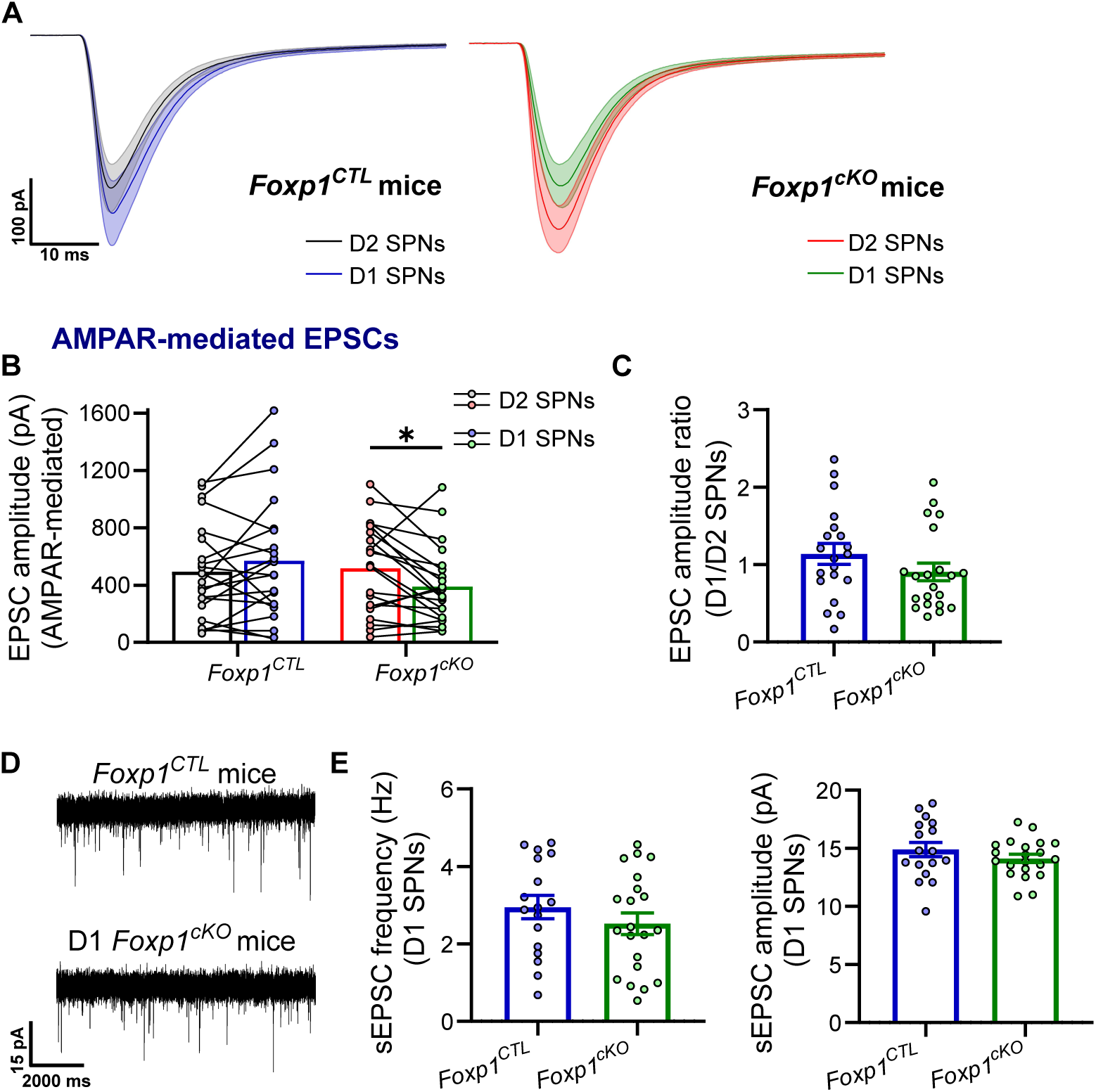
Excitatory cortical inputs are mostly unchanged in D1 SPNs with D1-specific *Foxp1* deletion. **A.** Average (± SEM) EPSC responses from D1 and D2 SPNs in D1 *Foxp1^CTL^*mice (blue and black respectively, *left*) and from D1 *Foxp1^cKO^*mice (green and red respectively, *right*). **B.** Amplitude of AMPAR-mediated EPSCs for D1/D2 SPN pairs in D1 *Foxp1^CTL^* and D1 *Foxp1^cKO^* mice (Interaction term, genotype x cell type; p<0.01). **C.** D1/D2 ratio of EPSC amplitude for each D1/D2 pair. **D.** Example traces of sEPSC in D1 SPNs. **E.** Both frequency (*left*) and amplitude (*right*) of sEPSCs in D1 SPNs did not change with *Foxp1* deletion in D1 *Foxp1^cKO^* mice. Ns in all figures are (D1 *Foxp1^CTL^*; D1 *Foxp1^cKO^*). For A-C. (21 D1/D2 pairs, 3 mice; 21 D1/D2 pairs, 4 mice). E. (17 D1 SPNs, 3 mice; 21 D1 SPNs, 4 mice). Statistics: B. 2-way ANOVA with Fisher’s LSD test. C. MW-test. E. T-test.

**Figure 7.**
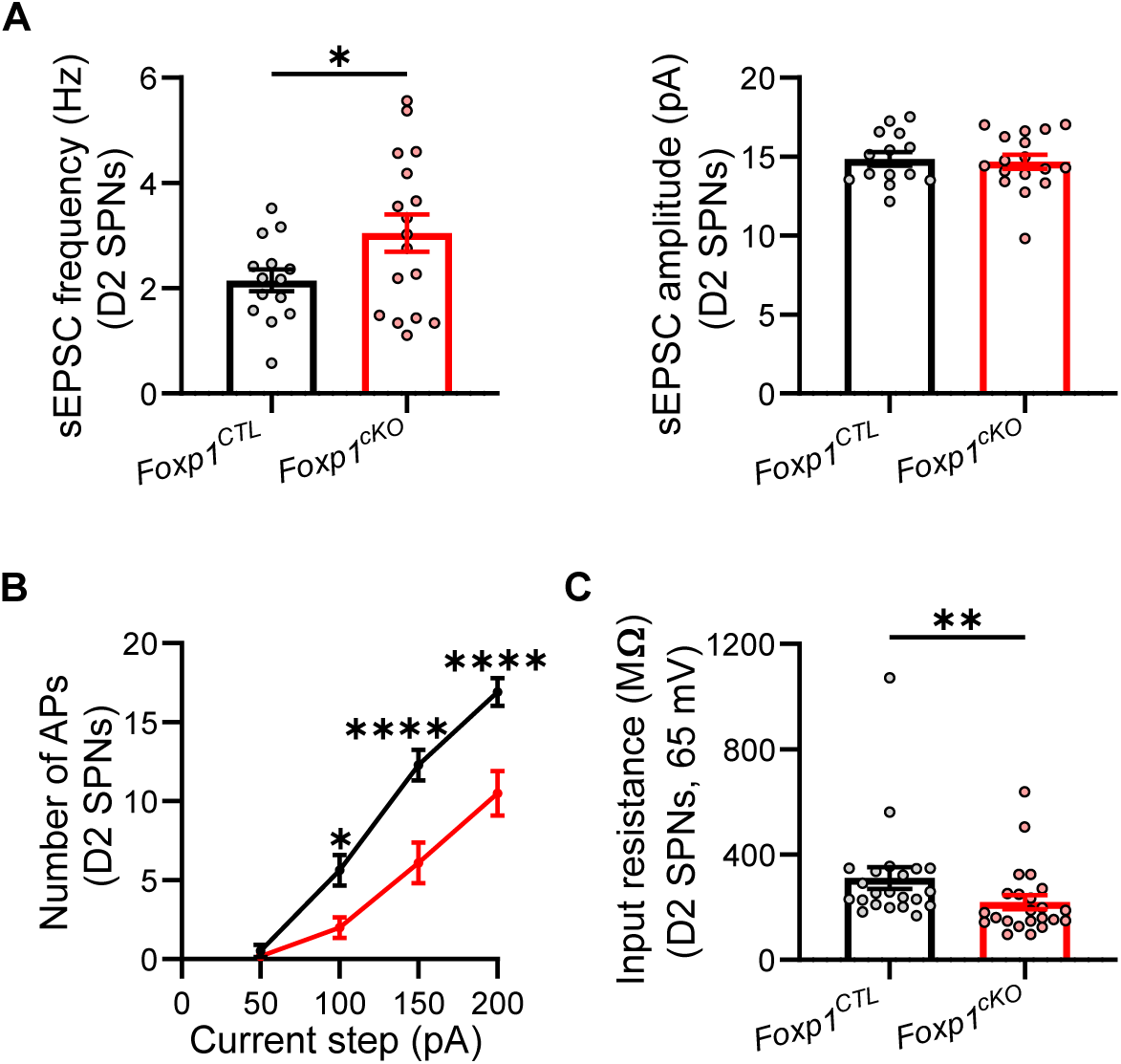
Non-cell autonomous effects are observed in D2 SPNs with *Foxp1* deletion in D1 SPNs in D1 *Foxp1^cKO^* mice. **A.** sEPSC frequency increases in D2 SPNs in D1 *Foxp1^cKO^* mice (*left*), however, the amplitude is not affected (*right*). **B, C.** D2 SPNs are intrinsically hypoexcitable in D1 *Foxp1^cKO^* mice with reduced input resistance at -65 mV as compared to *Foxp1^CTL^* mice. Ns in all figures are (D1 *Foxp1^CTL^*; D1 *Foxp1^cKO^*). For A. (14 D2 SPNs, 3 mice; 17 D2 SPNs, 4 mice). B, C. (22 D2 SPNs, 3 mice; 23 D2 SPNs, 4 mice). Statistics: A. T-test. B. 2-way ANOVA with Hom Sidak correction for multiple comparisons. C. MW-test. * p<0.05, ** p<0.01, **** p<0.001.

### Synaptically driven excitability is reduced in D2 SPNs with Foxp1 deletion

We have previously reported increased intrinsic excitability in D2 SPNs resulting from *Foxp1* deletion ^19^ and reproduce this finding here (Fig. 2 B-C). While this might suggest heightened activity of these neurons within the intact circuit ^19^, the observed decrease in corticostriatal input suggests reduced activity. To determine how these changes are integrated to regulate synaptically driven excitability of D2 SPNs, we employed blue light and ChR2 to stimulate corticostriatal afferents and measured action potential firing in SPNs. Light of variable intensity was applied that evoked subthreshold EPSPs at lower intensities and action potentials at higher intensities (Fig. 8A, *methods*). For each neuron, we obtained plots of firing probability as a function of light intensity where the firing probability refers to the fraction of trials where at least one action potential was evoked.

**Figure 8.**
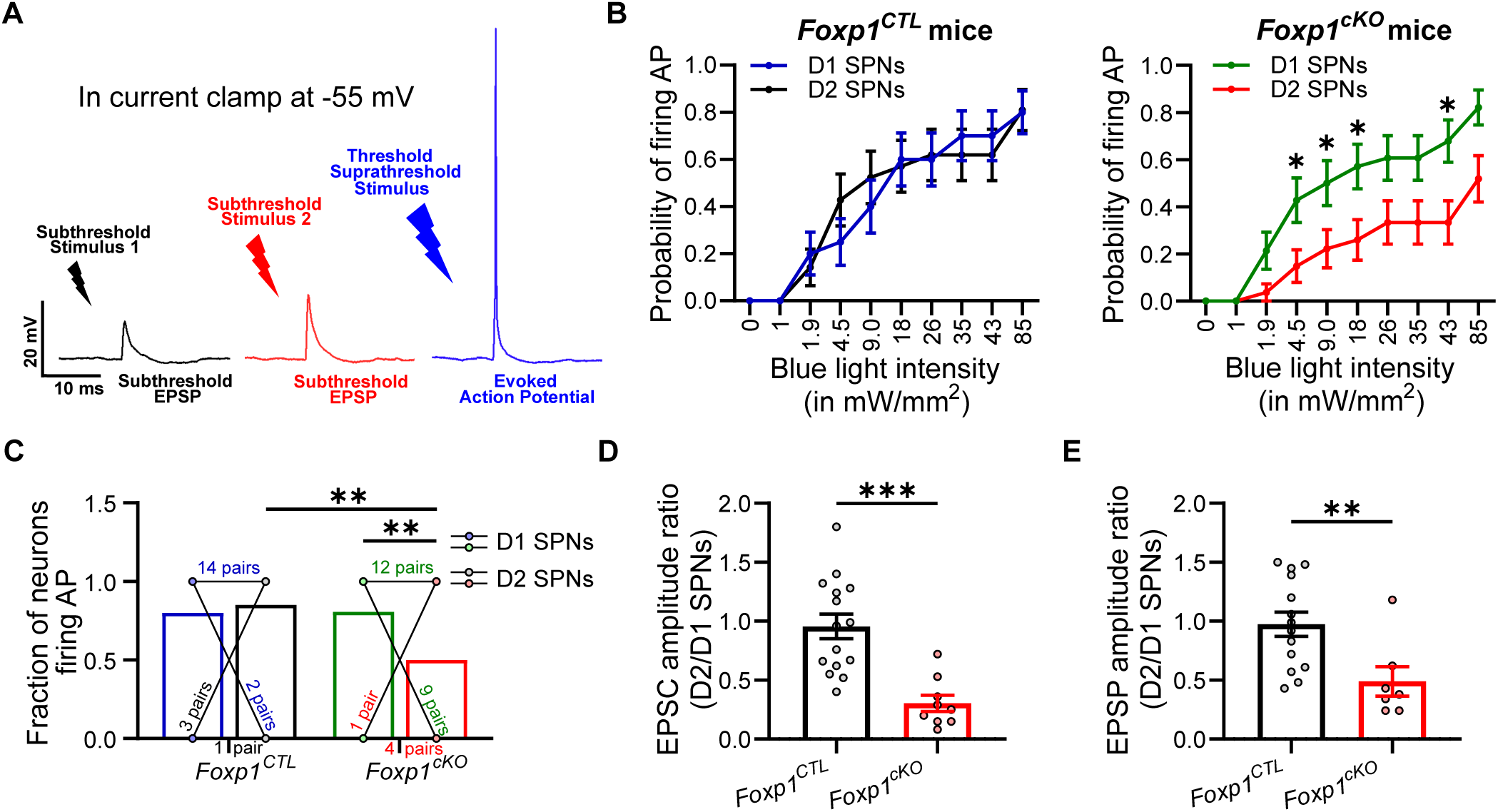
Synaptically induced excitability from the cortex is decreased in D2 SPNs with *Foxp1* deletion. **A.** Schematic of approach: blue light of different intensities where subthreshold stimuli generated EPSPs while threshold or suprathreshold stimuli generated action potentials. **B.** Excitability curves from D1/D2 SPN pairs in D2 *Foxp1^CTL^* control (*left*) and D2 *Foxp1^cKO^* mice (*right*). **C.** Fraction of SPNs with at least one action potential evoked at the maximum possible light intensity stimulus. 15% of D2 SPNs in *Foxp1^CTL^*mice and 50% in D2 *Foxp1^cKO^* mice did not ever fire action potentials. **D.** EPSC amplitude for each D1/D2 pair at -70 mV. **E.** D2/D1 ratio of EPSP amplitude for each D1/D2 pair. Ns in all figures are (D2 *Foxp1^CTL^*; D2 *Foxp1^cKO^*). For B, C. (20 D1/D2 pairs, 6 mice; 26 D1/D2 pairs, 7 mice). D. (15 D1/D2 pairs, 3 mice; 9 D1/D2 pairs, 2 mice). E. (14 D1/D2 pairs, 3 mice; 7 D1/D2 pairs, 2 mice). Statistics: B. Mixed effects model with Sidak correction for multiple comparisons. C. 2-way ANOVA with Fisher’s LSD test. D, E. T-test. ** p<0.01, *** p<0.001.

The average firing probability of D1 and D2 SPNs was equivalent in D2 *Foxp1^CTL^* mice whereas it was significantly reduced in D2 SPNs when compared with D1 SPNs in D2 *Foxp1^cKO^* mice (Fig. 8B). Moreover, the fraction of neurons capable of firing at least one action potential with the maximum light intensity was diminished in D2 SPNs in D2 *Foxp1^cKO^* mice (Fig. 8C). Additionally, the threshold intensity required to evoke an action potential was higher for D2 SPNs compared to D1 SPNs in D2 *Foxp1^cKO^* mice, whereas no difference was observed in D2 *Foxp1^CTL^* mice (Fig. 9A, *see methods for calculation*).

**Figure 9.**
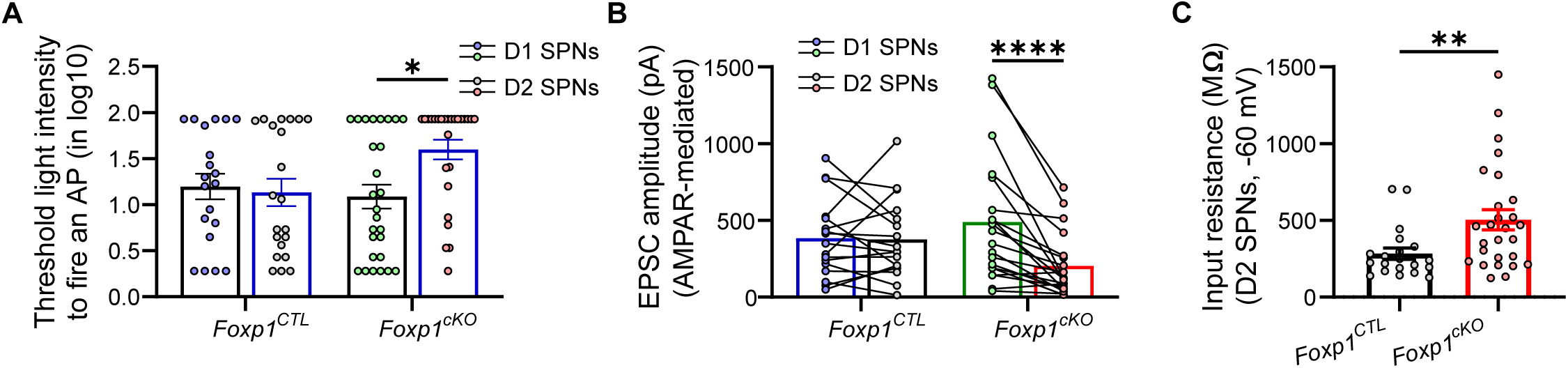
D2 SPNs are synaptically hypoexcitable in D2 *Foxp1^cKO^* mice. **A.** Log10 threshold stimulus to fire an action potential in D1/D2 SPN pairs in D2 *Foxp1^CTL^* and D2 *Foxp1^cKO^* mice. (*See methods for calculation*). **B.** EPSC amplitude of D1/D2 SPN pairs. **C.** D2 SPNs displayed an increase in input resistance in D2 *Foxp1^cKO^* mice. Ns in all figures are (D2 *Foxp1^CTL^*; D2 *Foxp1^cKO^*). For A, B. (20 D1/D2 pairs, 6 mice; 26 D1/D2 pairs, 7 mice). C. (20 D2 SPNs, 6 mice; 27 D2 SPNs, 7 mice). Statistics: A. 2-way ANOVA with Hom Sidak correction for multiple comparisons. B. 2-way ANOVA with Fisher’s LSD test. **C.** MW-test. * p<0.05, ** p<0.01, **** p<0.0001.

As expected, the relative EPSC amplitude was reduced (Fig. 9B, Fig. 8D) and input resistance increased ^19^ for D2 SPNs in D2 *Foxp1^cKO^* mice (Fig. 9C). The EPSP amplitude is arguably more relevant in determining synaptically driven excitability, and we would expect this to be weaker with *Foxp1* deletion. Indeed, this was roughly halved in D2 SPNs relative to neighboring D1 SPNs in D2 *Foxp1^cKO^*mice (Fig. 8E). Taken together, our results demonstrate that FOXP1 facilitates synaptically driven excitability mediated by corticostriatal inputs which is largely caused by a decrease in EPSP amplitude.

### Postnatal Foxp1 deletion weakens glutamatergic input onto D2 SPNs

The alterations in the synaptic physiology of D1/D2 SPNs were observed with embryonic *Foxp1* deletion where Cre-expression begins at E14-15 (*see methods*) ^8^. Hence, it remained unclear to what extent embryonic versus postnatal FOXP1 function underlies changes in glutamatergic inputs. Given that *Foxp1* plays a postnatal role in regulating intrinsic excitability of D2 SPNs ^19^, we hypothesized a similar postnatal function for *Foxp1* in regulating synapses. To test this hypothesis, we injected an AAV-Cre-eGFP virus into the striatum of P1 *Foxp1^flox/flox^* mice, resulting in *Foxp1* deletion by P6-P7 ^31, 32^. We employed extracellular electrical stimulation instead of ChR2-mediated stimulation to facilitate measurements of long-range glutamatergic input (*see methods*, Fig. 10A). We recorded AMPAR-mediated evoked responses in pairs of *Foxp1*/GFP-positive (*Foxp1^D2_vcKO^*) and negative (*Foxp1^D2_Flox^*) D2 SPNs.

**Figure 10.**
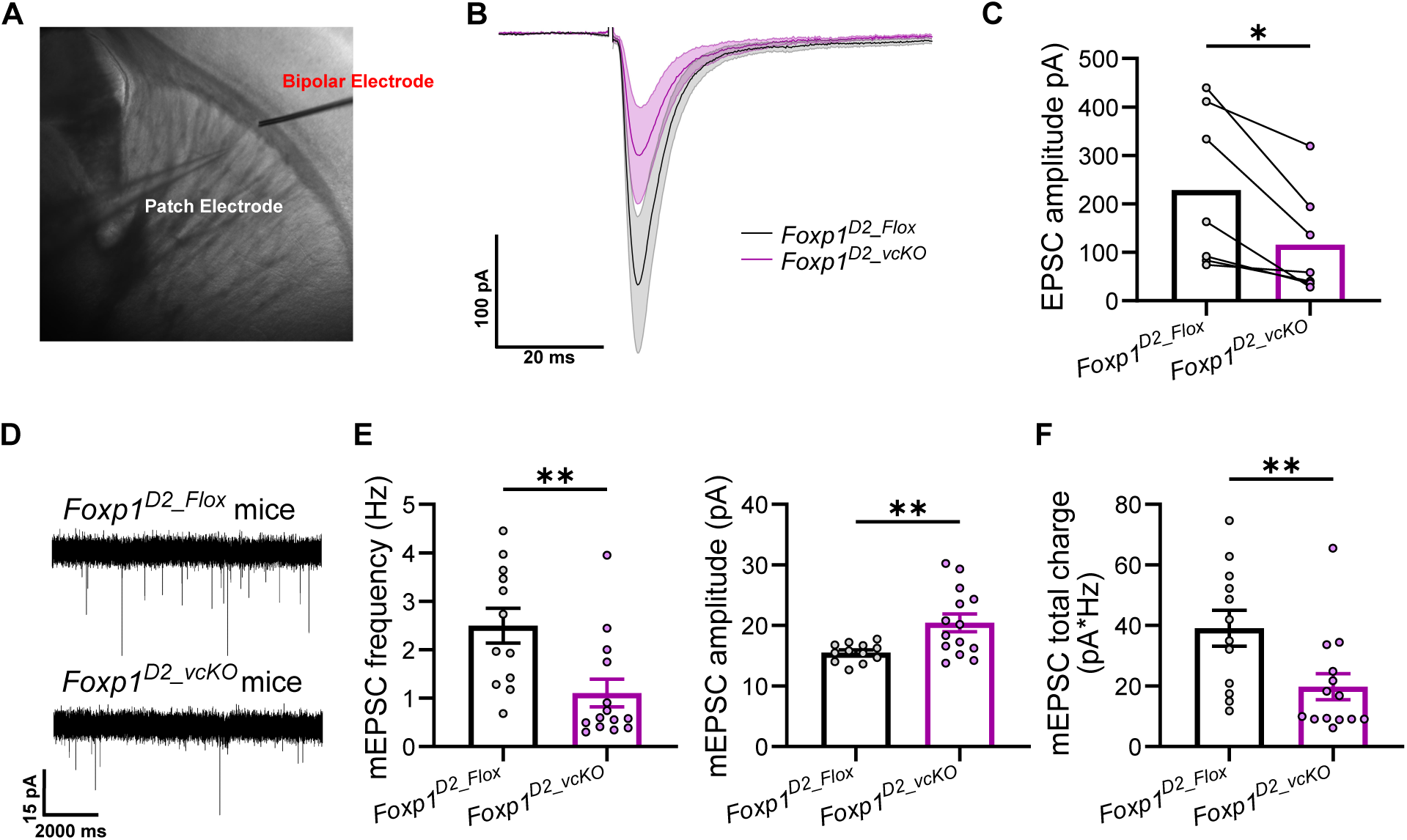
Excitatory cortical inputs onto D2 SPNs are decreased with postnatal deletion of *Foxp1*. **A.** Example image: A metal stimulating electrode was placed in underlying cortical white matter to stimulate the corticostriatal projections and responses were recorded in pairs of *Foxp1-*deleted (*Foxp1^D2_vcKO^*, eGFP positive) and control (*Foxp1^D2_Flox^*, eGFP negative) D2 SPNs. **B.** Average EPSC responses (±SEM) at -70 mV. **C.** D2 SPN with postnatal *Foxp1* deletion showed a reduction in EPSC amplitude. **D.** Example traces of mEPSCs in control and *Foxp1-*deleted D2 SPNs. **E.** mEPSC frequency (*left*) is reduced whereas mEPSC amplitude (*right*) is increased with postnatal *Foxp1* deletion in D2 SPNs. **F.** Total charge (frequency*amplitude) transfer showed a reduction in *Foxp1-*deleted D2 SPNs as compared to the control D2 SPNs. For B, C. 7 pairs of D2 SPNs in 4 mice. E, F. 12 *Foxp1^D2_Flox^* and 14 *Foxp1^D2_vcKO^* D2 SPNs from 3 mice. Statistics: C. Paired t-test. E (*left*), F. MW-test. E (*right*). T-test. * p<0.05, ** p<0.01.

Similar to embryonic deletion, EPSC amplitude was significantly smaller with postnatal *Foxp1* deletion compared to neighboring control D2 SPNs (Fig. 10B, C). We also observed a reduction in the frequency of mEPSCs with postnatal *Foxp1* deletion; however, unlike embryonic deletion, there was an increase in amplitude (Fig. 10D, E). This is consistent with fewer synapses but increased synaptic strength at each synapse. To assess the overall charge transfer through the mEPSCs, we multiplied the frequency and amplitude for each recording to obtain a single, “charge” parameter. Consistent with the reduced EPSC frequency, the total charge of mEPSCs was smaller with postnatal *Foxp1* deletion (Fig. 10F). Similar observations were made with sEPSC measurements (Fig. 11A, B). As expected, we observed an increase in the intrinsic excitability and input resistance of these D2 SPNs (Fig. 11C, D).

**Figure 11.**
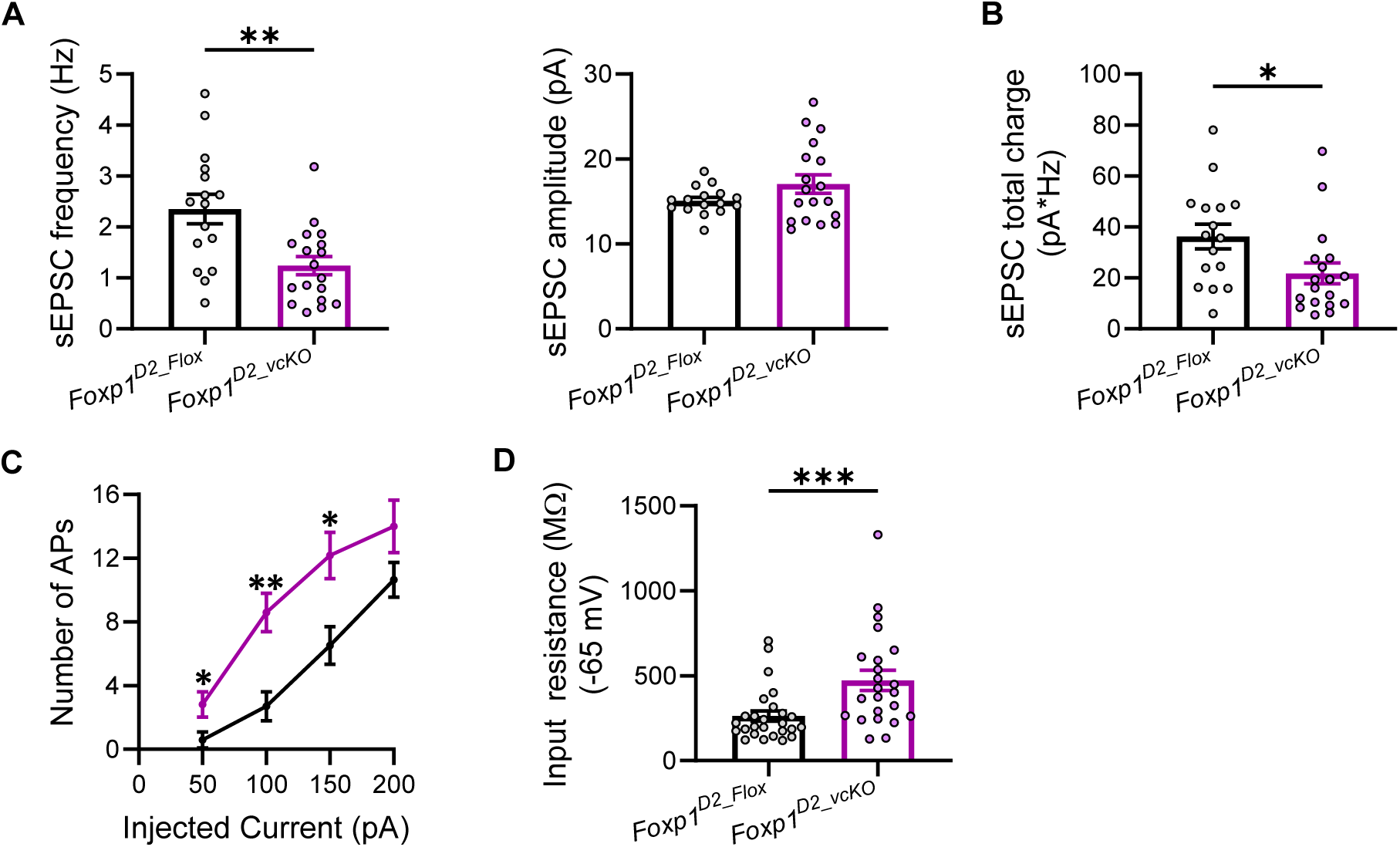
D2 SPNs with postnatal deletion of *Foxp1*. **A.** sEPSC frequency (*left*) is reduced while sEPSC amplitude (*right*) did not change with postnatal *Foxp1* deletion in D2 SPNs. **B.** Total charge transfer is also reduced in *Foxp1^D2_vcKO^*D2 SPNs as compared to the control D2 SPNs. **C, D.** D2 SPNs with postnatal *Foxp1* deletion were hyperexcitable and showed increased input resistance. For A. 16 *Foxp1^D2_Flox^*and 18 *Foxp1^D2_vcKO^*D2 SPNs, 6 mice. B, C. 24 *Foxp1^D2_Flox^* and 22 *Foxp1^D2_vcKO^* D2 SPNs from 7 mice. Statistics: A, B. T-test. C. Mixed effects model with Hom Sidak correction for multiple comparisons. D. MW-test. * p<0.05, ** p<0.01. **#** shows the genotypic difference in B.

Since postnatal deletion mimics the EPSC weakening observed with embryonic deletion, FOXP1-dependent positive regulation of excitatory inputs onto D2 SPNs is most likely due to postnatal FOXP1 function. Notably, while D2 SPN numbers decrease with embryonic deletion ^8^, this is not the case with postnatal deletion suggesting that the observed FOXP1 regulation of glutamatergic inputs is not an indirect effect of decreased number of D2 SPNs.

### Postnatal, postsynaptic reinstatement of Foxp1 is sufficient to rescue D2 SPN intrinsic hyperexcitability

We next tested whether postnatal *Foxp1* reinstatement is sufficient to rescue altered intrinsic excitability ^19^ and glutamatergic synaptic input in D2 SPNs. We cloned *Foxp1* cDNA and an mCherry reporter into an AAV construct driven by the human synapsin 1 (hSyn) promoter (pAAV-hSyn-*Foxp1*-T2A-mCherry or *Foxp1*^AAV^, Fig. 13A). Plasmids and viruses were tested in both HEK cells and the mouse striatum, respectively, to ensure the expression of FOXP1 protein (Fig. 13B, C, *see methods for details*). The control virus contained only mCherry reporter (pAAV-hSyn-mCherry or Control^AAV^). At P1, we injected D2 *Foxp1^cKO^* and D2 *Foxp1^CTL^* mice expressing *Drdr2-eGFP^tg/-^* with either *Foxp1*^AAV^ or Control^AAV^ into their striatum (Fig. 12A). The expression of reinstated *Foxp1* was first detected at P6 through mCherry fluorescence (Fig. 12B). Our recordings were limited to neurons positive for both eGFP (marker of D2 SPNs) and mCherry (marker for infected neuron) (Fig. 12B and Fig. 13D). We compared data among 4 groups: 1) D2 *Foxp1^CTL^* mice + Control^AAV^, 2) D2 *Foxp1^CTL^* mice + *Foxp1*^AAV^, 3) D2 *Foxp1^cKO^* mice + Control^AAV^, 4) D2 *Foxp1^cKO^* mice + *Foxp1*^AAV^.

**Figure 12.**
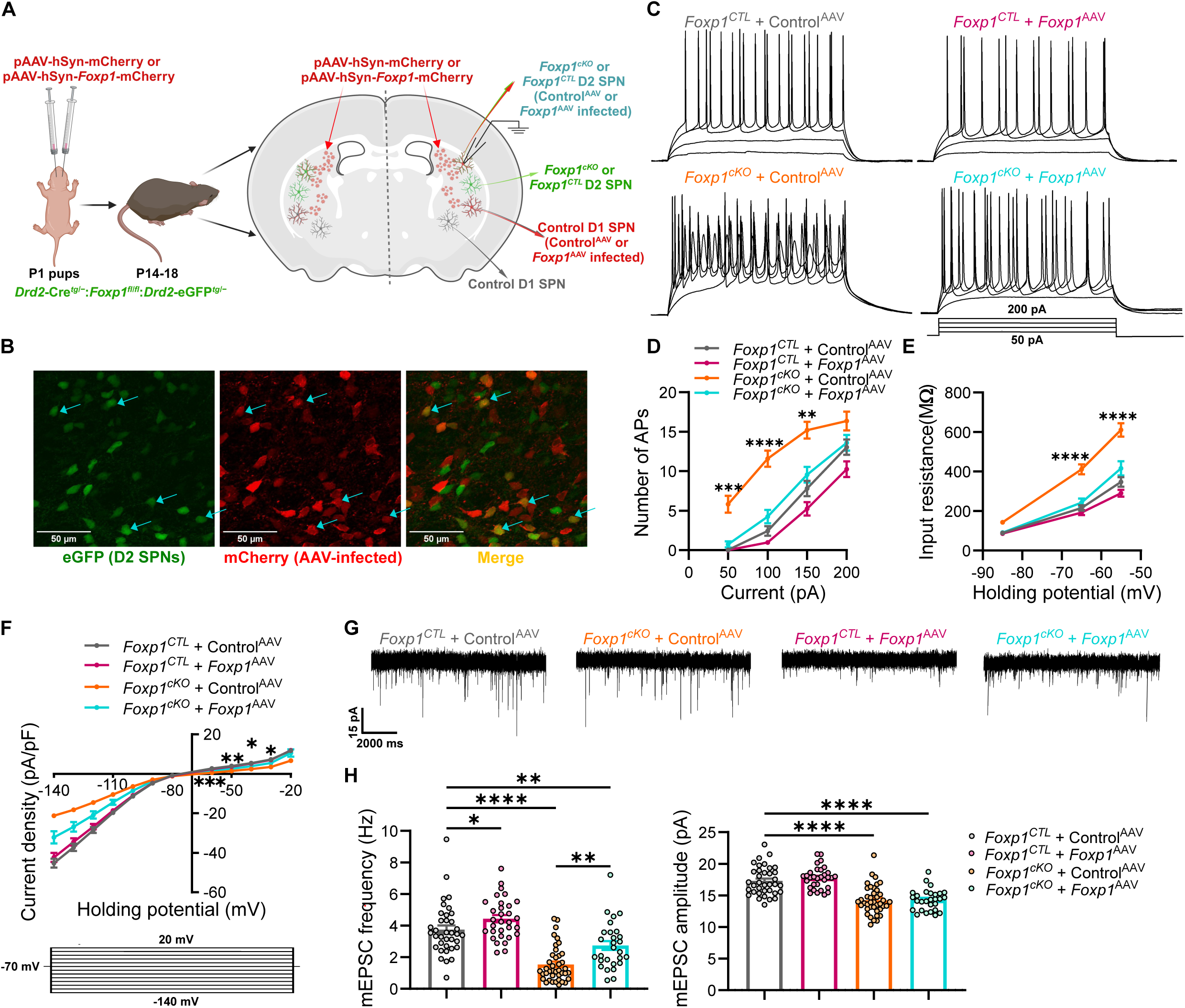
Postnatal *Foxp1* reinstatement in D2 SPNs rescues electrophysiological changes due to embryonic *Foxp1* deletion. **A.** Schematic describing injection of Control^AAV^ and *Foxp1*^AAV^ virus in the striatum at P1 and recording of D2 SPNs at P14-P18. **B.** eGFP positive D2 SPNs and mCherry positive AAV-infected neurons in D2 *Foxp1^cKO^*mice (P7). mCherry-positive neurons exhibit AAV expression as early as P7 in the mouse striatum. D2 SPNs with *Foxp1*^AAV^ infection (*merge, yellow*) are highlighted with arrows (*teal color*). **C.** Example traces of action potential firing in response to current injection (4 incrementing steps of 50 pA each) in D2 SPNs from all 4 mouse groups. **D.** Reinstatement of *Foxp1*^AAV^ in D2 SPNs in D2 *Foxp1^cKO^*mice rescues intrinsic hyperexcitability (*Teal color)* to the level of control D2 SPNs (*Gray color, Foxp1^CTL^* + Control^AAV^). **E.** Input resistance of *Foxp1*^AAV^ infected D2 SPNs in D2 *Foxp1^cKO^* mice decreased significantly to the level of control D2 SPNs. **F.** Current density versus voltage curve at different voltage steps (20 mV to -140 mV) shows partial rescue of currents in D2 SPNs with *Foxp1* reinstatement. **G.** Example traces of mEPSCs in D2 SPNs from all four mouse groups. **H.** mEPSC frequency is partially rescued with postnatal *Foxp1* reinstatement in D2 SPNs in D2 *Foxp1^cKO^* mice (*left*). mEPSC amplitude is not rescued in these D2 SPNs (*right*). Ns in all figures are (D2 *Foxp1^CTL^* + Control^AAV^; D2 *Foxp1^CTL^* + *Foxp1*^AAV^; D2 *Foxp1^cKO^* + Control^AAV^; D2 *Foxp1^cKO^* + *Foxp1*^AAV^). For D, E. (36 D2 SPNs, 3 mice; 34 D2 SPNs, 4 mice; 33 D2 SPNs, 5 mice; 32 D2 SPNs, 8 mice). F. (37 D2 SPNs, 3 mice; 34 D2 SPNs, 3 mice; 44 D2 SPNs, 4 mice; 31 D2 SPNs, 3 mice). H. (36 D2 SPNs, 3 mice; 31 D2 SPNs, 3 mice; 40 D2 SPNs, 4 mice; 28 D2 SPNs, 3 mice). Statistics: D-F. Mixed effect analysis with Hom Sidak correction for multiple comparisons. */**/*** Shows the statistics between rescued (*Foxp1*^AAV^ + D2 *Foxp1^cKO^*) and *Foxp1*-deleted (Control^AAV^ + D2 *Foxp1^cKO^*) D2 SPNs. H. One-way ANOVA with Hom Sidak correction for multiple comparisons. * p<0.05, ** p<0.01, *** p<0.001, **** p<0.0001.

**Figure 13.**
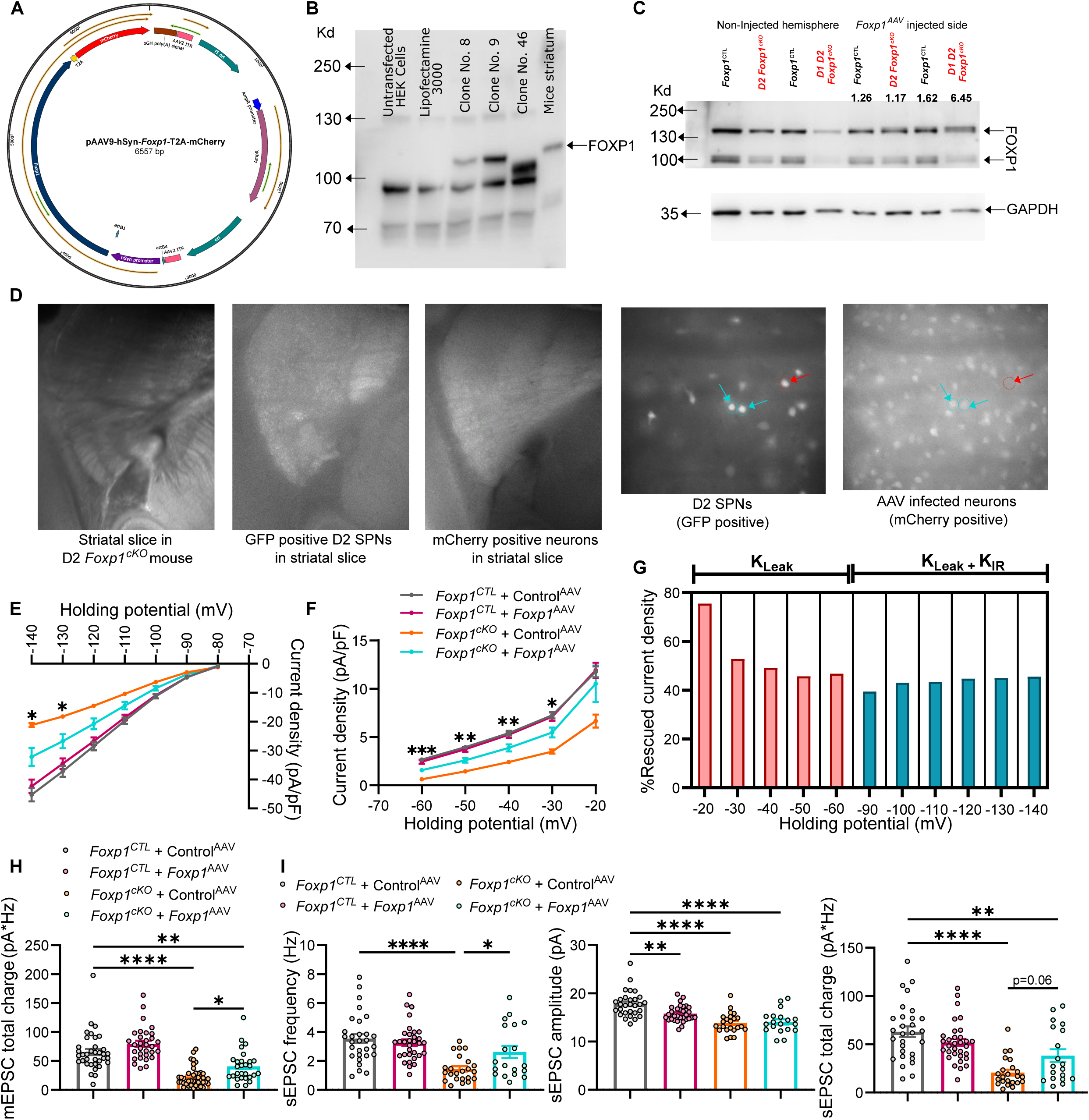
Postnatal reinstatement of *Foxp1* in mice striatum rescues the physiology of D2 SPNs. **A.** Map of *pAAV9-hSyn-Foxp1-T2A-mCherry* plasmid. **B.** FOXP1 immunoblot of HEK cells transfected with different *Foxp1* constructs. Clone 46 has truncated *Foxp1* whereas clones 8 and 9 have full-length *Foxp1*. Clone 9 was packaged into AAV and used for rescue experiments. The last lane depicts FOXP1 expression in mouse striatum and was used as a positive control. **C.** *Foxp1*^AAV^ was injected in the striatum of one hemisphere of *Foxp1^CTL^*, D2 *Foxp1^cKO^*, and D1D2 *Foxp1^cKO^* mice to test FOXP1 expression in mice brain with AAV injection. Protein was isolated from the striatum of both hemispheres separately and immunoblotted using antibodies against FOXP1 and GAPDH. The numerical values on the blot are the ratio of FOXP1 normalized (with GAPDH) intensities of injected hemisphere and non-injected hemisphere. A higher ratio suggests that *Foxp1^AAV^* was able to enhance FOXP1 expression. **D.** Brain slice showing the striatal region and GFP/mCherry fluorescence at low (4X) magnification (*3 images, left*) and eGFP/mCherry positive neurons at high (40X) magnification in D2 *Foxp1^cKO^*mice (*2 images, right*) at P16. eGFP indicates D2 SPNs while mCherry indicates *Foxp1*^AAV^ infection. Two arrows (*Teal color*) mark the D2 SPNs rescued with *Foxp1*^AAV^ and one red arrow marks non-rescued D2 SPN. **E, F.** Fig. 12F is replotted separately for hyperpolarized and depolarized potentials suggesting partially rescued currents at almost all potentials. **G.** Similar % rescued current density at different voltages suggests rescue of voltage-independent leak potassium current (K_Leak_) with *Foxp1* reinstatement. **H.** The total charge of mEPSCs is partially rescued with postnatal *Foxp1* reinstatement in D2 SPNs in D2 *Foxp1^cKO^* mice. **I.** The frequency (*left*) and total charge (*right*) of sEPSCs are partially rescued with postnatal reinstatement of *Foxp1* in D2 SPNs. sEPSC amplitude (*middle*) does not show rescue. Ns in all figures are (D2 *Foxp1^CTL^* + Control^AAV^; D2 *Foxp1^CTL^*+ *Foxp1*^AAV^; D2 *Foxp1^cKO^* + Control^AAV^; D2 *Foxp1^cKO^* + *Foxp1*^AAV^). For E-G. (37 D2 SPNs, 3 mice; 34 D2 SPNs, 3 mice; 44 D2 SPNs, 4 mice; 31 D2 SPNs, 3 mice). */**/*** Shows the statistics between rescued (*Foxp1*^AAV^ + D2 *Foxp1^cKO^*) and *Foxp1*-deleted (Control^AAV^ + D2 *Foxp1^cKO^*) D2 SPNs. H. (36 D2 SPNs, 3 mice; 31 D2 SPNs, 3 mice; 40 D2 SPNs, 4 mice; 28 D2 SPNs, 3 mice). I. (29 D2 SPNs, 3 mice; 32 D2 SPNs, 4 mice; 23 D2 SPNs, 5 mice; 19 D2 SPNs, 6 mice). Statistics: E. Two-way ANOVA with Hom Sidak correction for multiple comparisons. F. Mixed effect analysis with Hom Sidak correction for multiple comparisons. H, I. One-way ANOVA with Hom Sidak correction for multiple comparisons. * p<0.05, ** p<0.01, *** p<0.001, **** p<0.0001.

We measured the intrinsic excitability of the D2 SPNs with current injections of incremental size (Fig. 12C) and obtained curves of action potential number as a function of step size for each neuron. As expected ^19^, we observed intrinsic hyperexcitability with accompanying increased input resistance in D2 SPNs from the D2 *Foxp1^cKO^* + Control^AAV^ group (Fig. 12C-E). With *Foxp1* reinstatement, both excitability and input resistance were significantly reduced and not detectably different from D2 *Foxp1^CTL^* mice control groups (Fig. 12C-E).

The increased intrinsic excitability in D2 SPNs resulting from *Foxp1* deletion is primarily attributed to the downregulation of two potassium currents; K_IR_ - inward rectifying; K_Leak_ – leak.^19^. To investigate the effect of *Foxp1* reinstatement on K_IR_ and K_Leak_, we measured the induced currents in response to voltage steps (Fig. 12F). As anticipated, we replicated the decreased current density in D2 SPNs in the D2 *Foxp1^cKO^* + Control^AAV^ group at all voltage steps, and this reduction was partially rescued with *Foxp1* reinstatement (Fig. 12F). We previously demonstrated that differences in the upper right quadrant largely stem from reduced K_Leak_ whereas differences in the lower left quadrant result from both K_Leak_ and K_IR_ (Fig. 12F and Fig. 13E, F) ^19^. While K_IR_ is voltage-dependent, K_Leak_ is largely voltage-independent ^33–36^. Interestingly, the I-V curves in the *Foxp1^cKO^* + *Foxp1*^AAV^ group exhibited similar proportionate rescue in current density at nearly all voltage steps, indicating that the rescued current was voltage-independent. This suggests that postnatal *Foxp1* reinstatement primarily rescues K_Leak_ (Fig. 13G).

### Postnatal reinstatement of Foxp1 partially rescues the excitatory synaptic input onto D2 SPNs

We further investigated whether postnatal reinstatement of *Foxp1* could rescue the weakening of glutamatergic input onto D2 SPNs by measuring mEPSCs and sEPSCs. As anticipated for Control^AAV^ groups, frequency, amplitude, and total charge transfer by mEPSCs were decreased in D2 SPNs in D2 *Foxp1^cKO^* mice (Fig. 12G, H, Fig. 13H). Postnatal *Foxp1* reinstatement partially restored the frequency and total charge of mEPSCs in D2 SPNs of the D2 *Foxp1^cKO^* + *Foxp1*^AAV^ group (Fig. 12H *left*, Fig. 13H). Interestingly, the frequency of mEPSCs was further enhanced with reinstatement of *Foxp1* in D2 SPNs of *Foxp1^CTL^* mice, suggesting a direct *Foxp1*-mediated regulation. However, *Foxp1* reinstatement did not affect mEPSC amplitude when compared to controls (Fig. 12H *right*). Similar results were obtained for sEPSCs (Fig. 13I). This lack of change in mEPSC/sEPSC amplitude is consistent with the idea that FOXP1 determines EPSC amplitude embryonically and not postnatally (*see discussion*). To summarize, postnatal *Foxp1* reinstatement in D2 SPNs partially rescues the weakened excitatory inputs induced by embryonic *Foxp1* deletion.

### snRNA-seq of P18 SPNs with reinstatement of Foxp1

We investigated the cell-specific gene expression changes resulting from *Foxp1* reinstatement using the previously described AAV strategy and performed snRNA-seq on P18 striatal tissue – the same age as for electrophysiological experiments. We utilized three mice from each of the following groups: 1) D2 *Foxp1^CTL^* + Control^AAV^, 2) D2 *Foxp1^cKO^* + Control^AAV^, and 3) D2 *Foxp1^cKO^* + *Foxp1*^AAV^ (Fig. 14A). We obtained reads from a total of 89,745 nuclei across all three groups (Fig. 15A, B; *see methods for details*) and identified all major striatal cell types (Fig. 15C, D). Non-neuronal cells were distributed equally across all samples in the three genotypes (Fig. 15E). The number of D2 SPNs was reduced with *Foxp1* deletion, as previously reported ^8^, and this was accompanied by an increase in the number of eSPNs, progenitors, and neurogenic progenitors (Fig. 15E).

**Figure 14.**
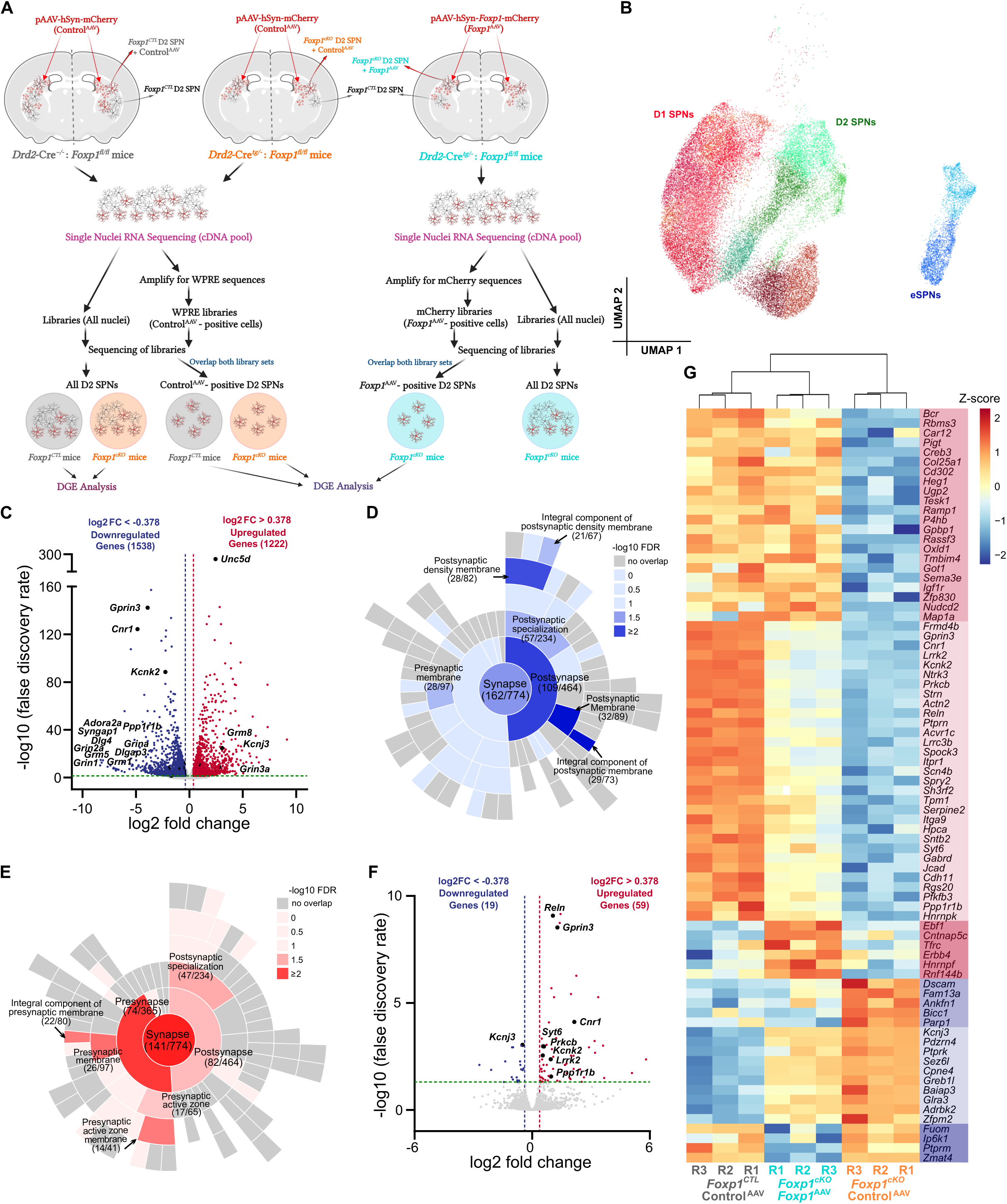
Single nuclei RNA sequencing of the striatum with postnatal reinstatement of *Foxp1.* **A.** Schematic showing the strategy of differential gene expression (DGE) analysis of (i) “all” D2 SPNs from D2 *Foxp1^CTL^* mice and D2 *Foxp1^cKO^* mice and (ii) “selected” D2 SPNs from D2 *Foxp1^CTL^*mice infected with Control^AAV^ and D2 SPNs from D2 *Foxp1^cKO^* mice infected with Control^AAV^ and *Foxp1*^AAV^. **B.** UMAP for SPNs from the subset of all cells from three groups of mice shows distinct clusters of the three SPN types. Different shades of red, green, and blue are indicative of subclusters in D1 SPNs, D2 SPNs, and eSPNs respectively. **C.** Volcano plot from DGE analysis of D2 SPNs from D2 *Foxp1^CTL^* and D2 *Foxp1^cKO^* demonstrate 1,538 genes downregulated and 1,222 genes upregulated with *Foxp1* deletion in D2 SPNs. (FDR ≤ 0.05, and log2 fold change ≥ 0.378). **D.** Gene ontology plot for synaptic genes using *SynGO* for downregulated DEGs demonstrates their involvement in postsynaptic function. The gray color represents the subcategories of synaptic genes that do not overlap with the DEGs. DEGs with significant overlap with synaptic genes (-log10 FDR>1.3) are highlighted with blue color where darker blue color indicates smaller FDR values. The number of DEGs overlapping with the total number of synaptic genes in the list is mentioned below the name of each category. **E.** Gene ontology plot for synaptic genes using *SynGO* for upregulated DEGs shows their association with post and presynaptic functions, however, there was more overlap with presynaptic categories. Gray color corresponds to no overlap, while red color demonstrates significant DEGs where darker red color depicts smaller FDR values. **F.** Volcano plot from DGE analysis for D2 SPNs from D2 *Foxp1^cKO^* mice infected with Control^AAV^ and *Foxp1*^AAV^ show 19 downregulated and 59 upregulated genes with *Foxp1* reinstatement in these SPNs. (FDR ≤ 0.05, and log2 fold change ≥ 0.378). **G.** Expression heat map of 78 genes in each sample from three groups (*Foxp1^CTL^* + Control^AAV^, *Foxp1^cKO^* + Control^AAV^, and *Foxp1^cKO^* + *Foxp1*^AAV^). Gene symbols in red and blue shades are upregulated and downregulated DEGs in D2 SPNs with *Foxp1* reinstatement in D2 *Foxp1^cKO^* mice respectively.

**Figure 15.**
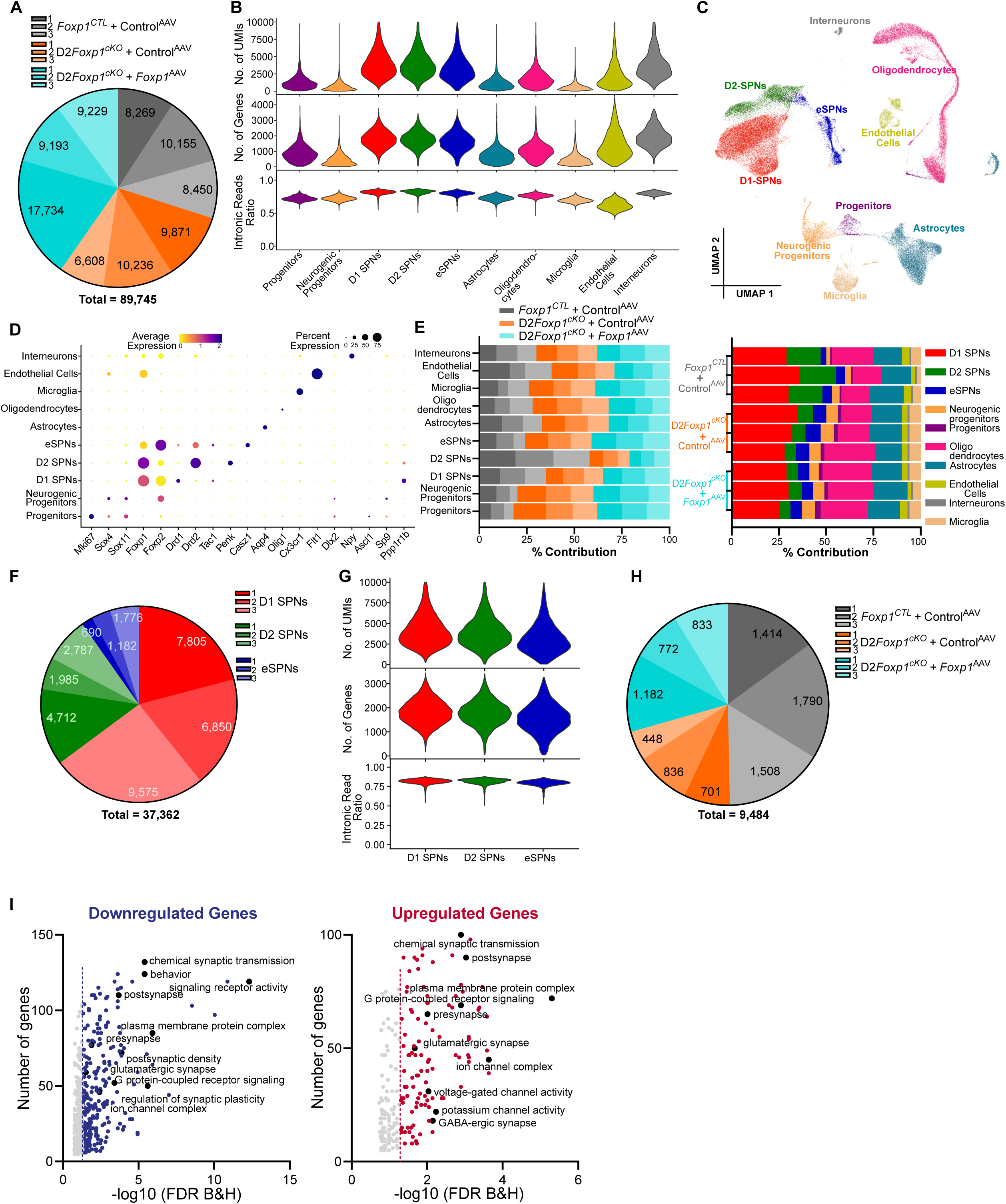
SnRNA sequencing of mouse striatum with postnatal *Foxp1* reinstatement. **A.** The pie chart depicts the total number of nuclei after filtering that were used for analysis from three samples in each group. **B.** Violin plots demonstrate quality control parameters (i.e., number of genes, number of UMIs, and intronic read ratio) separately for all detected cell types. **C.** UMAP shows major cell-type clusters from all nuclei from 9 striatal samples. **D.** Dot plot indicating the average and percentage expression of different marker genes in individual clusters for cell-type annotation. **E.** Bar plot showing the contribution of individual samples to different cell types (l*eft*) and contribution of individual cell types to different samples (*right*) normalized with total number of nuclei from each respective sample. There is a decrease in the number of D2 SPNs and an increase in the number of eSPNs and progenitors in D2 *Foxp1^cKO^*mice as compared to D2 *Foxp1^CTL^* mice. The proportional distribution of these cell types is not rescued with postnatal reinstatement of *Foxp1*^AAV^ in D2 *Foxp1^cKO^* mice. **F.** 37,362 SPNs (D1, D2, and eSPNs) were reclustered and the pie chart displays the contribution of each of these SPNs from three mice groups. **G.** Quality control parameters (i.e., number of genes, number of UMIs, and intronic read ratio) for all three SPN types that were used for further analysis. **H.** Pie chart depicts the total number of D2 SPNs from all 9 samples that were used for DGE analysis. **I.** Gene ontology pathways for both down and upregulated genes (FDR < 0.05 and log2 fold change ≥ 0.378) in D2 SPNs in D2 *Foxp1^cKO^* mice as compared to D2 *Foxp1^CTL^* mice. Pathways with FDR values < 0.05 are labeled with blue and red dots respectively, whereas FDR > 0.05 are in gray dots*. ClusterProfiler* followed by *Revigo* was used to generate the list of pathways.

### Embryonic Foxp1 deletion in D2 SPN results in the downregulation of postsynaptic genes

Next, we focused on examining the D2 SPNs in isolation by subclustering only the SPNs and retained 37,362 SPNs (Fig. 15F, G and Fig. 14B). To identify differentially expressed genes (DEGs) in D2 SPNs resulting from embryonic *Foxp1* deletion, we compared D2 *Foxp1*^CTL^ + Control^AAV^ mice with D2 *Foxp1*^cKO^ + Control^AAV^ mice (Fig. 15H). Our analysis revealed the downregulation of 1,538 genes and the upregulation of 1,222 genes in D2 SPNs of D2 *Foxp1*^cKO^ + Control^AAV^ mice (Fig. 14C, Table S1). Gene ontology analysis of both upregulated and downregulated DEGs show their enrichment in pathways related to (i) postsynaptic and presynaptic transmission, especially glutamatergic synapses, (ii) ion channels, particularly potassium channels, (iii) G-protein coupled receptor signaling, and (iv) the formation of plasma membrane complexes (Fig. 15I, Table S2). Using *SynGO* for a detailed synaptic ontology, we found that most of the downregulated genes were associated with postsynaptic functions (Fig. 14D, Table S3) while upregulated genes were enriched in both pre- and postsynaptic categories (Fig 7E, Table S3). This indicates a differential role for *Foxp1*-dependent regulation of synaptic genes where positive regulation of synaptic genes has a strong postsynaptic bias.

Consistent with the *SynGO* findings, an analysis of downregulated genes highlights ionotropic and metabotropic glutamate receptors as possible candidates for the reduced excitation onto D2 SPNs in D2 *Foxp1^cKO^* mice. We found a significant downregulation of *Grin1a* and *Grin2a* transcripts, which encode the NMDAR subunits GluN1 and GluN2, respectively, as well as group I metabotropic glutamate receptors, *Grm1 and Grm5*. Alternatively, reduced excitatory transmission may be due to downregulation of postsynaptic scaffold genes that are involved in glutamatergic transmission, such as *Dlg4* (encoding PSD95) and *Dlgap3* (also known as *Sapap3* or *Gkap3*) ^37–39^ ^40–42^. The expression of AMPAR subunit encoding genes did not change with the loss of *Foxp1*.

### Postnatal reinstatement of Foxp1 reverses the expression changes of several genes that are dysregulated with Foxp1 deletion in D2 SPNs

Next, we examined the gene expression changes in D2 SPNs after postnatal *Foxp1* reinstatement. Due to the inability to target all D2 SPNs with AAV injection, we used a strategy to isolate infected D2 SPNs (*see methods*) and performed differential gene expression analyses for two comparisons: 1) “embryonic deletion” using D2 *Foxp1^cKO^* + Control^AAV^ and D2 *Foxp1^CTL^* + Control^AAV^ mice (Fig. 16A-C), and 2) “postnatal reinstatement” using D2 *Foxp1^cKO^* + *Foxp1*^AAV^ and D2 *Foxp1^cKO^* + Control^AAV^ mice (Fig. 14F, G). For the embryonic deletion analysis, we identified 1196 downregulated and 942 upregulated genes in D2 *Foxp1^cKO^* + Control^AAV^ mice (Fig. 9C, Table S1). These DEGs greatly overlapped with the “all D2 SPNs” analysis described above (Fig. 14C). In the postnatal reinstatement analysis, we found 59 genes upregulated and 19 genes downregulated in *Foxp1*^cKO^ + *Foxp1*^AAV^ mice (Fig. 14F, G). Remarkably, 68 out of 78 of these postnatal reinstatement DEGS were also DEGs in the embryonic deletion analysis, but the transcript level changes for all 68 were reversed compared to deletion, and hence were considered rescued (Fig. 14G). Of these, 27 DEGs were not significantly different from *Foxp1*^CTL^ + Control^AAV^ mice indicating a full rescue (Fig. 14G). The remaining 10 out of 78 DEGs were not identified as DEGs in the embryonic deletion analysis and were exclusively regulated postnatally by *Foxp1*.

**Figure 16.**
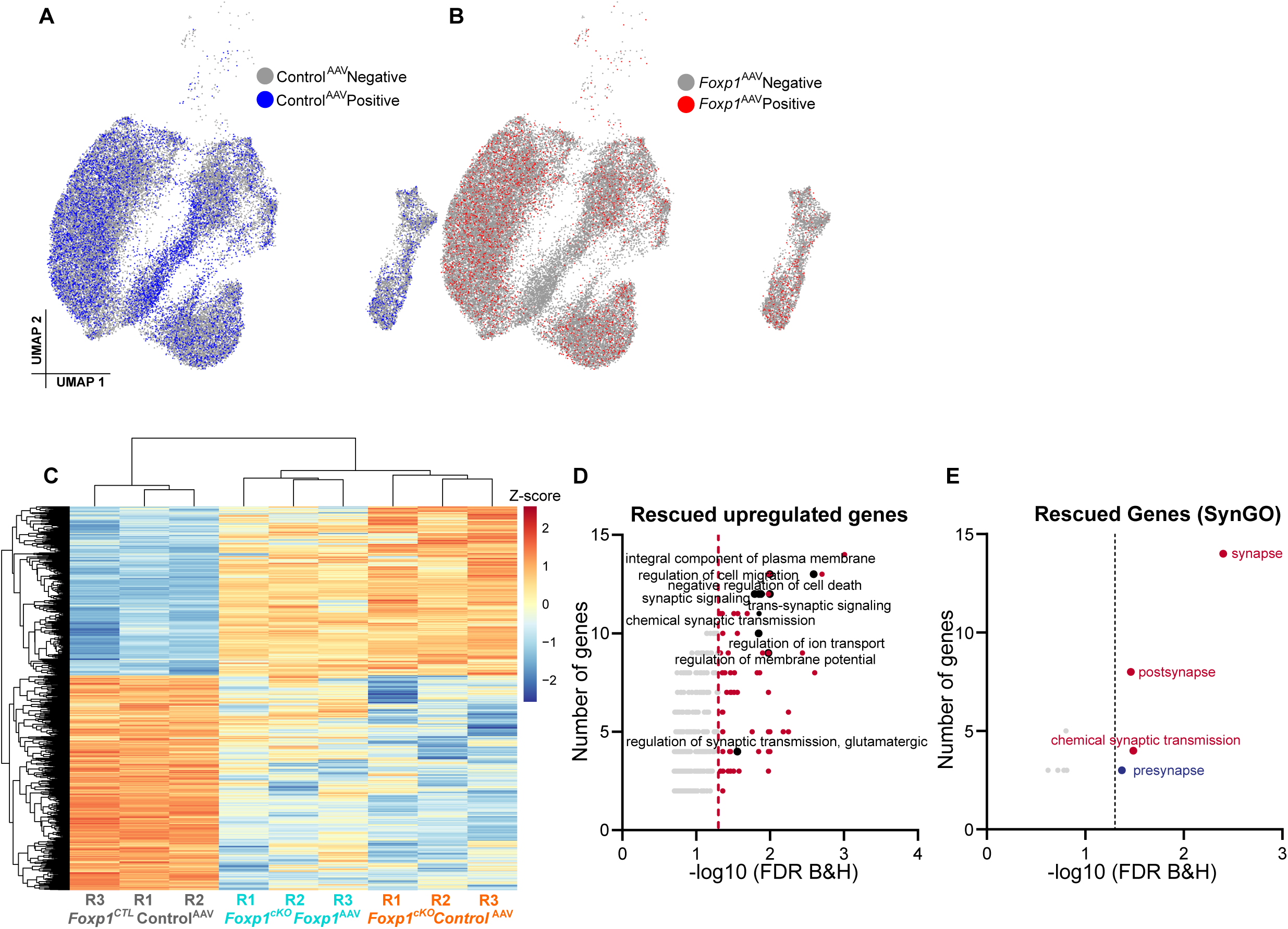
SnRNA sequencing analysis of Control^AAV^/ *Foxp1*^AAV^ positive D2 SPNs. **A.** The UMAP for SPNs where neurons positive for Control^AAV^ from D2 *Foxp1^CTL^* and D2 *Foxp1^cKO^* mice are highlighted with blue color. **B.** Similarly, UMAP for SPNs where neurons positive for *Foxp1*^AAV^ from D2 *Foxp1^cKO^* mice are highlighted with red color. **C.** Heatmap depicts the expression of DEGs from the comparison of Control^AAV^ positive D2 SPNs from D2 *Foxp1^CTL^* and D2 *Foxp1^cKO^* mice in all three groups (FDR < 0.05 and absolute log2 fold change ≥ 0.378). **D.** Gene ontology of the upregulated genes from the comparison of *Foxp1^cKO^* + *Foxp1*^AAV^ and *Foxp1^cKO^* + Control^AAV^, using *ClusterProfiler* followed by *Revigo*. **E.** *SynGO* pathway analysis for the rescued genes from Fig. 14I shows upregulated genes involved in postsynapse and downregulated genes involved in presynapse.

Gene ontology analysis of the upregulated rescued genes revealed enrichment in pathways related to synapse regulation, AMPK signaling, cell migration, and negative regulation of apoptosis (Fig. 16D, Table S2). *SynGO* analysis suggested a preferential rescue of downregulated postsynaptic genes and upregulated presynaptic genes (Fig. 16E, Table S3) – again consistent with FOXP1-dependent positive regulation of synaptic genes having a postsynaptic bias. Notably, there was an increased expression of genes involved in regulating plasticity and other aspects of striatal glutamatergic synaptic function, such as *Ppp1r1b* (DARPP32), *Reln*, *Sema3A*, and *Lrrk2* ^43–50^, suggesting potential candidates for the rescued synaptic function phenotype observed with *Foxp1* reinstatement. Relevant to our observation of FOXP1-dependent regulation of excitability, *Kcnk2* (Fig. 12F) ^19^, the gene that encodes leak potassium channel, is downregulated with embryonic *Foxp1* deletion and rescued with postnatal *Foxp1* reinstatement (Fig. 14G). Finally, the rescued expression of genes involved in the negative regulation of apoptosis such as *Igf1r*, *Spry2*, and *Sema3A* indicates a role for *Foxp1* in survival of D2 SPNs as its loss leads to a reduced number of D2 SPNs.

### Reinstatement of Foxp1 rescues behavioral phenotypes induced by Foxp1 deletion in both D1 and D2 SPNs

We next determined if postnatal *Foxp1* reinstatement can rescue the previously reported motor behavior alterations in D2 *Foxp1^cKO^* mice ^8^. We injected Control^AAV^ and *Foxp1*^AAV^ bilaterally into the striatum of both D2 *Foxp1^CTL^* and D2 *Foxp1^cKO^* mice at P1 and tested their behavior at 8-10 weeks of age. D2 *Foxp1^cKO^* mice recapitulated the motor learning deficit on the rotarod; however, this phenotype was not rescued with *Foxp1* reinstatement (Fig. 17A).

**Figure 17.**
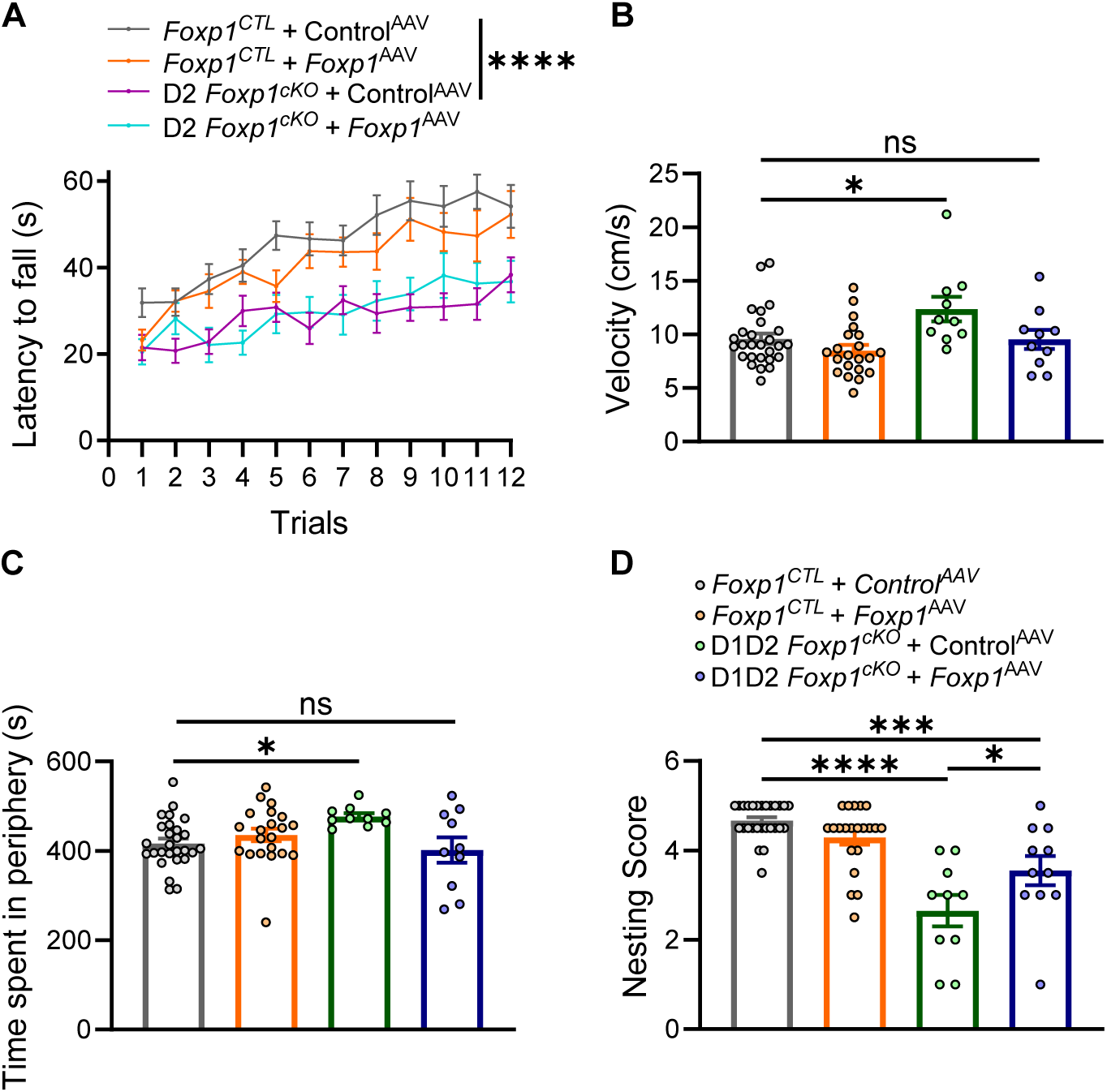
Postnatal *Foxp1* reinstatement rescues the behavior of mice with the loss of *Foxp1* in D1 and D2 SPNs. **A.** D2 *Foxp1^cKO^* mice showed a significant decrease in the latency to fall from the rotarod that was not rescued with postnatal *Foxp1* reinstatement. **B, C.** Mice with *Foxp1* deletion in both D1 and D2 SPNs showed an increase in the velocity and time spent in the periphery of the open field box. Both phenotypes were rescued with postnatal *Foxp1* reinstatement in these mice. **D.** The deficit in the nest building ability of D1D2 *Foxp1^cKO^* mice was also partially rescued. We used the same control mice (*Foxp1^CTL^*) for the comparison with D2 *Foxp1^cKO^* and D1D2 *Foxp1^cKO^* mice groups. Number of mice were: 27 *Foxp1^CTL^* + Control^AAV^; 21 *Foxp1^CTL^* + *Foxp1*^AAV^; 20 D2 *Foxp1^cKO^* + Control^AAV^; 15 D2 *Foxp1^cKO^* + *Foxp1*^AAV^; 10 D1D2 *Foxp1^cKO^*+ Control^AAV^; 10 D1D2 *Foxp1^cKO^* + *Foxp1*^AAV^ mice. Statistics: A. Two-way ANOVA with Hom Sidak correction for multiple comparisons. B-D. One-way ANOVA with Hom Sidak correction for multiple comparisons. * p<0.05, *** p<0.001, **** p<0.0001.

To further test the potential of postnatal *Foxp1* reinstatement to rescue motor deficits with loss of *Foxp1*, we used mice with simultaneous *Foxp1* deletion in both D1 and D2 SPNs (D1D2 *Foxp1^cKO^*). These mice exhibit more robust behavioral changes, making them suitable for detecting phenotypic rescue. In line with previous work ^8^, D1D2 *Foxp1^cKO^* + Control^AAV^ mice displayed hyperactivity and spent more time in the peripheral zone of the open field paradigm compared to *Foxp1^CTL^* + Control^AAV^ mice (Fig. 17B, C). With *Foxp1* reinstatement, the velocity and time spent in the periphery zone were no longer detectably different from control mice, suggesting a complete rescue of the phenotype (Fig. 17B, C). Additionally, the previously reported nesting impairment observed in D1D2 *Foxp1^cKO^*mice was partially rescued with *Foxp1* reinstatement (Fig. 17D). Nevertheless, rotarod deficits were not rescued in these mice (data not shown). In summary, postnatal reinstatement of *Foxp1* was able to rescue some of the behavioral phenotypes driven by simultaneous embryonic *Foxp1* deletion in both striatal SPN subtypes.

## Discussion

### Summary

Using cell-specific embryonic deletion mouse models, we find that the transcription factor, FOXP1, promotes the maturation and strengthening of long-range glutamatergic inputs selectively onto D2 SPNs in the striatum. This process is mediated by mechanisms that are cell-autonomous, postsynaptic, and postnatal. These effects are driven by a FOXP1-dependent transcription program that involves synaptic genes. This regulation differs between pre- and postsynaptic genes where positively regulated genes showed a strong postsynaptic bias (Fig. 14D, E). Postnatal *Foxp1* reinstatement successfully rescued both physiological and behavioral phenotypes. D2 SPNs with reinstated *Foxp1* exhibited a reversal in the expression of several genes that had been altered by embryonic deletion, indicating a rescued profile.

### Electrophysiology of excitatory input to SPNs

Synaptogenesis and strengthening of glutamatergic inputs onto SPNs are most prominent between P8 and P20 ^3, 4, 51^. This process is governed by intracellular signaling pathways as well as presynaptic and postsynaptic activity ^2–5^. However, the role of transcription factors in directing these inputs is relatively underexplored ^10^. Our results were obtained during this critical developmental time-window and connect FOXP1 function with strengthening of these inputs. Moreover, during this time, maturation of glutamatergic inputs onto SPNs involves a decreased contribution of NMDAR-mediated transmission^29^. Our findings, illustrated by an increased proportion of NMDAR-mediated responses following *Foxp1* deletion (Fig. 3D), suggest that FOXP1 plays a facilitative role in the maturation of excitatory synapses by reducing the relative proportion of synaptic NMDARs.

In addition to the direct action of FOXP1, it is possible that the glutamatergic synaptic input changes resulting from either *Foxp1* deletion or reinstatement are mediated by the change in intrinsic excitability through a FOXP1-independent process. Conversely, the inverse could also hold true, where alterations in synaptic inputs induce changes in intrinsic excitability. Previous studies have already established a close relationship between intrinsic excitability and synaptic transmission ^52–55^. Therefore, the observed reduction in synaptic excitation induced by *Foxp1* deletion might be a FOXP1-independent homeostatic response to compensate the increased intrinsic excitability ^56^. Similarly, the combination of reduced intrinsic excitability and increased synaptic excitation in hippocampal neurons following *Foxp1* deletion may also represent a similar homeostatic response ^18^.

In our experimental paradigm, contralateral inputs originate from a single population of intratelencephalic layer 5 pyramidal neurons^26^ devoid of any thalamic contribution. Consequently, the light-evoked responses observed in contralateral SPNs were induced from well-isolated corticostriatal inputs (Fig. 1-10). In contrast, our measurements of mEPSC/sEPSC and responses evoked by extracellular electrical stimulation included long-range glutamatergic inputs originating from both cortical and thalamic neurons (Fig. 1-12). However, the fact that we observed a similar effect on both evoked (light and electrical) and spontaneous recordings suggest a common regulation of both input pathways or a strong regulation of corticostriatal inputs that can be observed even in the presence of thalamic inputs. Either way, changes observed with these measurements likely reflect changes in corticostriatal inputs.

### Foxp1 is required for both embryonic and postnatal development of corticostriatal circuitry

FOXP1 is crucial for normal brain development during embryonic and postnatal stages. During early embryonic stages, it is involved in the self-renewal and differentiation of neural stem cells ^57, 58^, formation of midbrain dopaminergic neurons ^59^, and migration of several neuronal types within the developing cortex ^58, 60–62^. Additionally, in the striatum, embryonic FOXP1 is essential for the development of D2 SPNs ^8^. During postnatal stages, FOXP1 continues to promote the survival of SPNs while inhibiting apoptosis and oxidative stress ^20, 63, 64^. Our findings are consistent with a postnatal function for FOXP1 in regulating the intrinsic excitability of D2 SPNs ^19^ and glutamatergic synaptic inputs onto these neurons (Fig. 1, 3, 1). We show that postnatal function is both necessary and sufficient for regulation of these physiological properties by implementing postnatal deletion and reinstatement, respectively (Fig. 10, 12). These data also indicate that the decrease in D2 SPN numbers induced by embryonic deletion is a distinct phenomenon and is independent of the accompanying electrophysiological changes.

An exception to the role of postnatal FOXP1 regulation of excitatory input is related to mEPSC/sEPSC amplitude. The frequency and amplitude of mEPSCs/sEPSCs attain maturity at different time points in SPNs during postnatal development^29^. Interestingly, we also observe a differential FOXP1-mediated development of these properties. In contrast to embryonic deletion, postnatal *Foxp1* deletion did not reduce mEPSC/sEPSC amplitude (Fig. 10, 11 vs. Fig. 1, S1, S2). Moreover, the amplitude remained unchanged with postnatal reinstatement of FOXP1 (Fig. 12, 13). On the contrary, mEPSC/sEPSC frequency decreased with both embryonic and postnatal deletion and was subsequently rescued with postnatal reinstatement (Fig. 1-4, 10, 12, 13). Assuming the frequency of mEPSCs/sEPSCs represents the number of synapses and amplitude corresponds to the strength of individual synapses, our data suggests a dual FOXP1 function to promote both the number of synapses and synaptic strength.

### Postsynaptic candidate genes for FOXP1 synaptic regulation supported by embryonic Foxp1 deletion

Our data strongly suggests that FOXP1 positively regulates glutamatergic inputs onto D2 SPNs, likely by modulating the expression of key synaptic genes. Interestingly, a significant number of downregulated synaptic genes in D2 SPNs following embryonic *Foxp1* deletion are primarily postsynaptic and are relevant with the observed synaptic phenotypes. We found decreased expression of *Grin1a, Grin2a, Grm1* and *Grm5,* which is consistent with the reduction in NMDAR-mediated currents. *Grin1a/2a* encode NMDAR subunits, while *Grm1/5* encode for mGluR1/5. Both mGluR1/5 are abundantly expressed postsynaptically in striatal SPNs and are involved in excitatory synaptic transmission by activating the phospholipase C pathway^65, 66^. Moreover, mGluR1/5 potentiate the activation of NMDARs in SPNs by enhancing their expression on the membrane surface ^65–68^. Although we observed higher expression of *Grin3a* (GluN3) and *Grm8* (mGluR5), it is not expected to enhance glutamatergic transmission. GluN3 subunits are primarily activated by glycine, not glutamate ^69^ and mGluR8 is expressed presynaptically, where it decreases NMDA receptor activity. Despite a substantial decrease in AMPAR-mediated transmission with embryonic deletion, we did not observe changes in the transcripts encoding AMPAR subunits. However, we did observe downregulation of *Dlg4* and *Dlgap3*, which regulate postsynaptic receptor trafficking and AMPAR-mediated transmission, and therefore could account for the reduced AMPAR currents ^37–42^. In summary, we observe transcription changes in postsynaptic genes that may directly and indirectly participate in FOXP1-dependent regulation of synaptic function in D2 SPNs.

### Postnatal Foxp1 reinstatement rescues gene expression and cellular electrophysiology

*Foxp1* deletion in D2 *Foxp1^cKO^* mice occurs during an early phase of striatal development (E14-15), leading to a substantial reduction in the population of D2 SPNs. Postnatal reinstatement of *Foxp1* expression occurred at P6-7, a stage when most neurons have largely acquired their identity. This postnatal reinstatement of FOXP1 effectively rescues most of the physiological properties of D2 SPNs while also altering the expression of certain genes. Most importantly, 87% of these altered genes (68 out of 78 “postnatal reinstatement” DEGs) were also identified as “embryonic deletion” DEGs. However, their expression levels were reversed with postnatal reinstatement, indicating a “genetic” rescue. Therefore, most of the reinstatement DEGs were in fact a subset of and were not different from the “embryonic deletion DEGs”. Importantly, while reinstatement DEGs represent a minority among the large number of embryonic deletion DEGs, their impact is substantial enough to rescue the affected neuronal physiology of D2 SPNs. For instance, specific candidate genes observed with embryonic deletion - like *Grin1a/2a* or *Grm1/5 -* did not exhibit differential gene expression during postnatal reinstatement. This suggests that the transcriptional mechanisms underlying embryonic deletion and postnatal reinstatement may overlap but at the same time, differ, perhaps due to differences in electrophysiological and synaptic processes that occur during embryonic versus postnatal periods.

Alternatively, upregulation of other postsynaptic genes that were rescued with reinstatement may be more directly involved in FOXP1-dependent regulation of glutamatergic synaptic function. In fact, some of these genes may have been more relevant to embryonic deletion as compared to the obvious receptor and scaffolding candidates discussed above. For example, Leucine-rich repeat kinase 2 (*Lrkk2*) is highly expressed in the SPNs ^48, 70–72^ and plays a critical role in regulating synaptogenesis and corticostriatal glutamatergic synaptic function ^47, 73, 74^. Another gene of interest, Protein phosphatase 1 regulatory inhibitor subunit 1B (*Ppp1r1b, also known as* DARPP32) is critical for the corticostriatal synaptic plasticity and controls both long term potentiation and depression in the striatum ^44–46^. Lastly, *Reln* is another gene that has been implicated in regulating the synaptic plasticity in the striatum and hippocampus ^43, 75^. Altogether, our data suggest that these genes likely contribute to the intricate network of molecular factors that regulate synaptic function and plasticity through regulating AMPAR/NMDAR mediated currents in D2 SPNs.

The rescued expression of *Kcnk2* is particularly noteworthy, as it appears to be tightly regulated by FOXP1 and is consistently associated with its function (K_Leak_) in our studies ^8, 17, 19^. In our previous study, we showed that *Foxp1* deletion in D2 SPNs leads to intrinsic hyperexcitability, primarily due to reduced K_Leak_ and decreased expression of *Kcnk2* ^19^. In contrast, *Foxp1* reinstatement enhances *Kcnk2* levels, which likely rescues the intrinsic excitability phenotype by increasing K_Leak_. This suggests a transcriptional mechanism through which FOXP1 directly regulates K_Leak_ and thereby intrinsic excitability. To our knowledge, this is one of the very few clear and reproducible demonstrations of a transcriptional mechanism regulating electrophysiological function in neurons ^24^.

## Supporting information

DEG Analysis

Pathway Analysis_Clusterprofiler

Pathway Analysis_Syngo

## Acknowledgements

Our sincerest thanks to Dr. Kimberly Huber and Dr. Todd Roberts for providing critical feedback on the manuscript. We thank Dr. Shin Yamazaki and the Neuroscience Microscopy Facility, supported by the UT Southwestern Neuroscience Department and the UTSW Peter O’Donnell, Jr. Brain Institute. G.K. is a Jon Heighten Scholar in Autism Research and Townsend Distinguished Chair in Research on Autism Spectrum Disorders at UT Southwestern. This work was supported by grants from NIMH (MH126481, MH102603), NINDS (NS126143, NS115821), and the James S. McDonnell Foundation 21st Century Science Initiative in Understanding Human Cognition – Scholar Award (220020467) to G.K and the Simons Foundation (573689) to G.K. and J.G.

## Author Contributions

N.K., J.G., and G.K. designed the study and wrote the paper. N.K. performed all the experiments. A.K. analyzed the data for single-nuclei RNA sequencing experiments. N.A. contributed to mice behavior experiments. M.H. contributed to the maintenance of mice colonies and performed genotyping of the experimental mice.

## Declaration of interests

The authors declare no competing interests.

## Inclusion and diversity

We support inclusive, diverse, and equitable conduct of research.

## STAR Methods

### Resource availability

#### Lead contact

Further information and requests should be directed to and will be fulfilled by the Lead Contact, Jay Gibson and Genevieve Konopka.

### Materials availability

Animals and materials generated from this study are available from the lead contact with a completed Materials Transfer Agreement.

### Experimental mice

We used C57BL/6J background mice for our experiments consistent with our previous study ^19^. We used *Foxp1*^flox/flox^ mice ^21, 76^ and BAC-transgenic lines for fluorescence reporter: *Drd2*-eGFP^tg/-^ and *Drd1α*-tdTomato^tg/-^ ^77^ and for Cre: *Drd2*-Cre^tg/-^ and *Drd1α*-Cre^tg/-^ ^78^. *Drd2-*specific *Foxp1* conditional knockout mice were *Drd2*-Cre^tg/-^:*Foxp1*^flox/flox^:*Drd2*-eGFP^tg/-^ (D2 *Foxp1^cKO^*) whereas control mice were *Foxp1*^flox/flox^:*Drd2*-eGFP^tg/-^ (D2 *Foxp1^CTL^*). Similarly, *Drd1α-*specific *Foxp1^cKO^* were also triple transgenic: *Drd1α*-Cre^tg/-^:*Foxp1*^flox/flox^:*Drd1α*-tdTomato^tg/-^ (D1 *Foxp1^cKO^*) mice and control mice are *Foxp1*^flox/flox^:*Drd1α*-tdTomato^tg/-^ (D1 *Foxp1^CTL^*) ^19^. Both Cre lines turn on between embryonic day (E) 14 and E15, which coincides with the start of *Foxp1* expression in striatal cells ^8^. We used both male and female mice which were maintained on 12 h light on/off schedule with access to food and water *ab limitum*. All experiments were performed according to the procedures approved by the UT Southwestern Institutional Animal Care and Use Committee.

### Electrophysiology methods

#### Brain slices and Recordings

We used mice aged postnatal day (P) 15-18 for the electrophysiological measurements. Mice were anesthetized with a combination of ketamine (125 mg/kg) and xylazine (25 mg/kg), and brains were quickly dissected and transferred to partially frozen dissection buffer ^19^. 300 μm thick brain slices containing the dorsal striatum were prepared using a Vibratome (*Leica VTS 1200S*) and immediately placed in nominal artificial cerebrospinal fluid (nACSF) (bubbled with a mixture of 5% CO_2_ and 95% O_2_), first at 35°C for 30 min, and then at room temperature for at least 30 min before commencement of recordings ^19, 79, 80^. We used glass pipettes of 5-8 MΩ resistance to perform whole cell patching on D1 and D2 SPNs in D1 and D2 (*Foxp1^cKO^*or *Foxp1^CTL^*) mice, which were identified based on the expression of tdTomato (tdTom) and eGFP reporter, respectively. In certain instances, we adopted a “negative” identification approach where cells negative for eGFP and tdTom fluorescence were considered as D1 and D2 SPNs respectively. This approach proved reliable as 95% of striatal neurons are SPNs and can be distinguished based on their smaller size ^77^. We included neurons with a maximum acceptable resting membrane potential of -50 mV, capacitance greater than 15 pF, stable baseline current within 15 pA, and a series resistance of less than 20 MΩ in our analysis. The average series resistance did not exhibit significant differences between *Foxp1^cKO^* and *Foxp1^CTL^* neurons. The junction potential was ∼10 mV and was not corrected.

All recordings were performed at a temperature of 30°C. To maintain consistency with our previous study, we adopted the same nomenclature for most of the experiments. Wild-type (WT) control neurons and *Foxp1*-deleted neurons were referred to as *Foxp1^CTL^* and *Foxp1^cKO^* D1/D2 SPNs, respectively. For the D2 SPNs, where *Foxp1* was deleted postnatally using AAV-mediated Cre expression, uninfected neurons were named *Foxp1^D2_Flox^* D2 SPNs while AAV-infected neurons were named *Foxp1^D2_vcKO^* SPNs.

Unless stated otherwise, we recorded and analyzed all electrophysiological parameters using custom software (*Labview; National Instruments, Austin, TX*; https://github.com/ColdP1228/Custom-Data-Acquisition-Program- ^31^; copy archived at https://github.com/elifesciences-publications/Custom-Data-Acquisition-Program-).

#### Current steps, input resistance and multi-voltage step protocols

For these measurements, we followed our previously published protocol ^19^. Briefly, we applied incremental current steps (50 pA, 500 ms duration) at the resting potential in current clamp mode to assess intrinsic excitability. The resulting number of action potentials elicited at each step was plotted to generate “firing versus current injection” curve.

To determine input resistance, we applied a single -10 mV voltage step (500 ms duration) in voltage clamp mode and measured input resistance based on the average current values within a 100 ms window before and 200 ms after the step onset. Input resistance was measured either at -85, -65, and -55 mV holding potentials or solely at -65 mV holding potential depending on the specific experimental requirements.

For the IV plots at various voltages, we applied a multi-step voltage protocol (10 mV steps, 500 ms duration) while maintaining a holding potential of -70 mV (ranging from -20 to -140 mV voltages). We measured the average current within a 200 ms window at the end of the step and divided it by the capacitance to obtain the current density. Capacitance was measured as described earlier ^19^. For these measurements, nACSF contained tetrodotoxin (TTX, 1 µM).

#### Calculating rescued current density

For fig. 13G, we first calculated the loss of current due to *Foxp1* deletion by subtracting the current densities of D2 SPNs in D2 *Foxp1^cKO^* + Control^AAV^ from those in D2 *Foxp1^CTL^* + Control^AAV^ mice at each voltage. Rescued current density was calculated as the difference between D2 *Foxp1^cKO^* + *Foxp1*^AAV^ and D2 *Foxp1^cKO^* + Control^AAV^ measurements. Finally, the ratio of rescued and lost current was the % rescued current density at respective voltages. The current density at -20 to -60 mV is exclusively K_Leak_ whereas it is a combination of K_Leak_ and K_IR_ at -90 to -140 mV voltages.

#### ChR2-mediated optogenetic stimulation of corticostriatal evoked postsynaptic currents EPSCs

We employed commercially prepared pAAV9.CAG.hChR2(H134R)-mCherry.WPRE.SV40 (*Addgene, Cat*#*100054*) viral particles expressing mCherry-tagged human channel rhodopsin (hChr2) to stimulate corticostriatal inputs. Briefly, we diluted the viral particles to a titer of ∼10^12^ viral particles (vp) per ml in sterile 1X PBS and added 1-2 µl of diluted Fast Green FCF dye (*Sigma)* for visualization and performed injections as described previously ^19, 31, 32^. P1 pups were anesthetized on ice, and we injected 350 nl of diluted viral particles into the superficial cortical layers (0.5 mm underneath the skull) on the left hemisphere using a beveled glass injection pipette and a Nanoject^TM^ II injector (*Drummond Scientific, Inc*.). The pups were allowed to recover on a heating pad until they regained consciousness and were then transferred to their home cage. Slices were prepared from P14-18 juvenile mice and contralateral projections from ChR2-AAV infected cortical neurons were stimulated with blue light to trigger synaptic responses in the striatal neurons in the right hemisphere. To avoid possible contamination by ChR2 expression in ipsilateral striatal neurons, we recorded from slices prepared from the contralateral right hemisphere. The strength of the evoked response in SPNs using this protocol could vary due to multiple factors, including variability in slice preparation, efficiency of viral infection, density of corticostriatal inputs, and ChR2 expression in the contralateral striatum (Fig. 1A). Such variability could obscure the changes in the amplitude of synaptic responses induced by *Foxp1* deletion. Therefore, to mitigate this concern, we measured EPSCs in neighboring neuron pairs (<50 µm distance) consisting of a GFP-positive D2 SPN and a GFP-negative D1 SPN in D2 *Foxp1^cKO^* mice. In this design, D1 SPNs consistently expressed *Foxp1* and served as a control comparison. Similarly, in D1 *Foxp1^cKO^*mice, we recorded from pairs consisting of a tdTom-positive-D1 SPN and tdTom-negative D2 SPN. For each recording pair, both the D1 and D2 SPN were stimulated with blue light of identical intensity and were patched in alternating sequence (Fig. 1A).

#### AMPAR and NMDAR mediated EPSCs

We recorded action potential-dependent AMPAR-mediated EPSCs from D2/D1 SPNs at a holding potential of -70 mV in striatal slices incubated in nACSF containing 2mM CaCl_2_ and 2mM MgCl_2_. Corticostriatal axons were stimulated by a 12 ms blue light (470 nm) pulse from a fluorescent lamp (*X-Cite, Lumen Dynamics*) through a 40x water immersed objective (*Olympus*). Similar intensity light, with power ranging from 1 to 85 mW/mm^2^ (resulting in a final beam diameter of 340 µm and power of 0.1 to 7.6 mW), was used to record responses from D1 and D2 SPNs in each pair.

We measured the amplitude, onset latency, response rise time, and slope of the EPSC responses. Amplitude is the maximum current response in pA to the individual light pulse. Onset latency is used as a measure to filter out non-synaptic responses. SPNs infected due to the leak of AAV-ChR2 from the ipsilateral cortex have onset latencies of 1-2 ms, however, all responses obtained from the SPNs in contralateral striatum had onset latency of more than 5 ms, indicative of pure synaptic currents. Response rise time is the time (in ms) to reach the peak response. Finally, we used a 5 - 15 ms time window, relative to the stimulus onset and measured the slope of the AMPAR-mediated EPSC responses between 20 - 80% of the rise time.

To study the effect of *Foxp1* deletion on release probability, we stimulated afferent axons with a train of 5 blue light pulses of 200 ms interval and 12 ms duration and recorded the responses from both D1 and D2 SPNs (Fig. 1G). The ratio of the peak amplitude of 2^nd^ to the 5^th^ response relative to the first amplitude response indicated release probability. If the relative decrease in input to the neurons is due to a decrease in release probability, we would expect an increase in EPSC amplitudes during the train.

We also measured AMPAR- and NMDAR-mediated responses in the absence of action potential firing. This protocol isolates monosynaptic EPSCs while avoiding contamination from disynaptic inputs and was based on a previous study^28^. The protocol involved recording in the presence of TTX (1 µM), picrotoxin (100 µM) and 4-AP (100 µM) to block voltage-dependent Na^+^ channels, GABA_A_ receptors, and K^+^ currents, respectively. We used an NMDAR agonist, Glycine (20 µM), and reduced MgCl_2_ (0.5 mM), to activate and record NMDAR-mediated currents while maintaining 3 mM CaCl_2_ concentration to balance cation levels. We also used Cesium based internal pipette solution to enhance the resolution of NMDAR-mediated currents.

We first recorded amplitude, slope, latency, and release probability of AMPAR-mediated evoked EPSCs as explained above at -70 mV holding potential. Subsequently, we depolarized the neurons to +50 mV to record NMDAR-mediated evoked EPSCs. Corticostriatal axons were stimulated with the same intensity of blue light while recording both D1 and D2 SPNs in a pair. NMDAR-mediated responses are slower in rise and decay than the AMPAR-EPSCs and initial 20-40 ms fast EPSC response could have contamination from AMPAR-mediated responses. Therefore, we measured the peak NMDAR-mediated amplitude in a 90 -100 ms window after the onset of EPSCs.

#### Spontaneous and miniature EPSCs

The synaptic inputs onto SPNs in our measurements likely involve a combination of corticostriatal and thalamostriatal inputs. However, assuming that corticostriatal inputs contribute to the AMPAR/NMDAR differences, we can relate changes in sEPSC/mEPSC to our corticostriatal results. For sEPSC recordings, neurons were voltage clamped at −70 mV and spontaneously occurring synaptic events were recorded in 10 traces, each lasting 10 s. Similarly, mEPSCs were recorded for the same duration with nACSF supplemented with TTX (1 µM), picrotoxin (100 µM) and 4-AP (100 µM). Data analysis was performed using an automatic detection program (*MiniAnalysis; Synaptosoft Inc, Decatur, Ga.*) with a detection threshold at a value exceedingly at least 5 standard deviations (10 pA) of the baseline noise. Additionally, we visually confirmed the detected events. The detection threshold remained constant throughout each recording.

#### Synaptically driven excitability

Synaptically driven excitability curves were generated by applying blue light (with a diameter of 300 µm, centered at the recorded cell) to depolarize corticostriatal axons expressing ChR2. D1/D2 SPNs were patched and maintained in current clamp at -55 mV and 12 ms blue light flashes with power ranging 1 to 85 mW/mm^2^ were applied efficiently to determine the threshold stimulus intensity. We started with half of the maximum power (40 mW/mm^2^) and adjusted the power either up or down depending on whether an action potential was evoked or not, respectively. When we approached the threshold intensity, we made smaller changes in the light intensity to precisely determine the threshold stimulus. To facilitate the comparison of the threshold intensity between two groups (Fig. 9A), we calculated the log10 value of the threshold stimulus for each neuron. For neurons that did not elicit an action potential even at the maximum possible light stimulus (85 mW/mm^2^), we considered maximum light intensity as the threshold intensity. In addition to estimating the threshold light intensity, we also calculated the probability of firing an action potential at various stimulus intensities for these neurons.

#### Electrophysiology Solutions

We prepared brain slices in a semi-frozen dissection buffer (bubbled with 95% O_2_ and 5% CO_2_) containing the following components (in mM): 110 Choline-Cl, 25 Dextrose, 25 NaHCO_3_, 3 Ascorbic Acid, 3.1 Sodium Pyruvate, 2.5 KCl, 1.25 NaH_2_PO_4_, 7 MgCl_2_, 0.5 CaCl_2_.2H_2_O, and 0.2 Kyneuric Acid, with a pH of 7.3-7.4 and an osmolality of 280-285 mOsm. The slices were transferred to nACSF that contained (in mM): 125 NaCl, 25 NaHCO_3_, 10 Dextrose, 3 KCl, 1.25 NaH_2_PO_4_, 2 MgCl_2_, and 2 CaCl_2_.2H_2_O with a pH of 7.3-7.4 and an osmolality of 300-305 mOsm, saturated with 95% O_2_/5% CO_2_. The pipette (internal) solution consisted of (in mM): 130 K-Gluconate, 10 HEPES, 6 KCl, 3 NaCl, 0.2 EGTA, 14 phosphocreatine-Tris, 4 ATP-Mg and, 0.4 GTP-Tris with a pH of 7.25 and an osmolality of 300 mOsm.

For NMDAR-mediated currents, nACSF was modified and consisted of (in mM): 125 NaCl, 25 NaHCO_3_, 10 Dextrose, 3 KCl, 1.25 NaH_2_PO_4_, 0.5 MgCl_2_, 3 CaCl_2_.2H_2_O, 0.1 picrotoxin, 0.1 4-AP and 0.02 Glycine. Also, the internal pipette solution was revised and contained (in mM): 125 Cs-Gluconate, 10 HEPES, 5 TEA-Cl, 3 NaCl, 2.5 BAPTA, 0.6 EGTA, 14 phosphocreatine-Tris, 4 ATP-Mg and, 0.4 GTP-Tris with a pH of 7.4 and an osmolality of 290 mOsm.

### Viral-mediated postnatal deletion of *Foxp1*

We used AAV-mediated Cre expression to perform postnatal *Foxp1* deletion. AAV9 particles expressing eGFP-Cre fusion protein (pAAV.CMV.HI.eGFP-Cre.WPRE.SV40) were obtained commercially (*Addgene, Cat#105545*). Briefly, viral particles were diluted to a titer of ∼10^12^ viral particles (vp) per ml in sterile 1X PBS with the addition of Fast Green FCF dye (*Sigma*) for visualization. P1 *Foxp1*^flox/flox^:*Drd1α*-tdTomato^tg/-^ C57BL/6J pups were anesthetized on ice, and 350 nl of diluted viral particles were injected into the striatum at a depth of 1.5 mm as explained above. For AAV-Cre mediated postnatal *Foxp1* deletion, we employed *Foxp1*^flox/flox^:*Drd1α*-tdTomato^tg/-^ mice and specifically targeted smaller, tdTomato-negative neurons, assuming these to be D2 SPNs ^77^. The eGFP fluorescence in the nuclei served as a marker for AAV-mediated Cre expression. In each case, an infected (GFP-positive) D2 or D1 SPN was paired with an adjacent (<50 µm distance) uninfected (GFP-negative) D2 or D1 SPN, which served as a control.

We did not employ ChR2-mediated stimulation of corticostriatal projections for the EPSCs measurement in D2 SPNs with postnatal *Foxp1* deletion. This was because strong mCherry fluorescence in the ChR2-expressing afferents made the reliable detection of tdTomato-positive D1 SPNs challenging. Instead, we induced EPSCs using an extracellular stimulating bipolar 2-conductor cluster electrode positioned in the cortical projections within approximately 50-100 µm distance from the recorded D2 SPN. We used monophasic current pulses (10–300 μA, 0.2–1 ms) for stimulation. The same current intensity was applied sequentially from the stimulating electrode to both infected and uninfected D2 SPN pair.

### Viral-mediated postnatal reinstatement of *Foxp1*

#### Cloning of Foxp1 into the AAV-hSyn vector

We cloned *Foxp1* cDNA, tagged with a fluorescence reporter mCherry (connected with T2A linker) into an AAV construct driven by the human synapsin (hSyn) promoter. Briefly, we initially amplified the vector backbone including the promoter sequence using the following primers: Vector_Fwd (5’GTACAAGTAGACCCAGCTTTCTTGTACAAAGTG3’) and Vector_Rev (5’CTTGCATCATGGTGGCAGCCTGCTTTTTTG3’). The *Foxp1* and mCherry sequences were amplified, with the T2A linker sequence added in between *Foxp1* and mCherry. *Foxp1* was amplified from a mouse cDNA library using *Foxp1*_Fwd_1 (5’ATGATGCAAGAATCTGGGTCTG3’) and *Foxp1*_Rev_1 (5’CTCCATGTCCTCATTTACTGGTTC3’). Thereafter, part of the T2A sequence was added to the 3’end of the *Foxp1* cDNA in two consecutive rounds of reactions, employing two reverse primers: *Foxp1*_Rev_2 (5’GATGTTAGAAGACTTCCCCTGCCCTCTCCGCTTCCCTCCATGTCCTCATTTACTG3’) and *Foxp1*_Rev_3 (5’CACGTCCCCGCATGTTAGAAGACTTCCC3’), along with the forward primer *Foxp1*_Fwd_2 (5’GGCTGCCACCATGATGCAAGAATCTGGG3’). Similarly, mCherry was amplified using another mCherry-containing plasmid as a template. The remaining part of the T2A sequence was added to the 5’end of the mCherry sequence by two consecutive PCR reactions utilizing mCherry_Fwd_1 (5’CGGGGACGTGGAGGAAAATCCCGGCCCCATGGTGAGCAAGGGCGAGGAGGATAA3’) and mCherry_Fwd_2 (5’TTCTAACATGCGGGGACGTGGAGGAAAATC3’), along with the reverse primer mCherry_Rev (5’AAAGCTGGGTCTACTTGTACAGCTCGTCCATG3’), respectively. Subsequently, the three fragments were ligated using the NEB Gibson assembly cloning kit (*NEB*, *Cat#E5510*) and the ligated product was transformed into NEB5-alpha competent *E. coli* cells (*NEB*, *Cat#C2987H*). Positive clones were verified through restriction digestion and PCR. The sequences of the positive clones were also confirmed with Sanger sequencing (Fig. 13A).

To confirm the expression of FOXP1 protein expression in these clones, we performed transfection in HEK293 cells followed by western blotting (Fig. 13B). Briefly, HEK cells were cultured at 37°C with 5% CO_2_ in 1X DMEM (*Gibco, Cat#12491015*) supplemented with 10% Fetal Bovine Serum and 1X Penicillin Streptomycin mix (*Gibco*, *Cat#15070063*). Cells in a 6-well plate were transfected at 70% confluency with 2 µg of the plasmid mixed with Lipofectamine 3000 transfection reagent (*Invitrogen, Cat#L3000001*) as per the manufacturer’s protocol. After 48 h of transfection, cells were washed with 1X PBS and lysed in 300 µl of 2x Laemmli Sample Buffer (*Biorad*, *Cat#1610737*), followed by incubation at 95°C for 5 min. Equal volumes of each lysate were loaded onto a 10% SDS PAGE and transferred onto a PVDF membrane. After blocking with 5% milk in TBST (1× TBS and 0.05% Tween20) for 1 h at room temperature, the membranes were incubated with primary antibodies in 2% BSA in TBST at 4°C overnight. Following washing with TBST, the blots were incubated with HRP-conjugated secondary antibodies, mixed in 5% milk in TBST, for 1 h at room temperature. The membranes were washed again and then developed using the SuperSignal™ West Pico PLUS Chemiluminescent Substrate (*Thermo Scientific, Cat#34580*). Signal acquisition was performed using a ChemiDoc MP (*BioRad*) and analyzed using ImageLab (*BioRad*) or Fiji.

Once the clones were confirmed, bacterial colonies for the selected plasmid were grown in 200 ml LB broth for 14-15 h at 37°C with agitation at 220 rpm in a shaking incubator to generate enough plasmid for AAV packaging. Bacterial cells were then pelleted, and plasmid isolation was carried out using NucleoBond Xtra Midi EF (*Machery Nagel*, *Cat#740420.50*) following the manufacturer’s protocol. A total of 140 µg of purified plasmid was packaged into AAV particles by the *Neuroconnectivity Core at Baylor college of Medicine* resulting in a titer of 9.42 x 10^12^ GC/ml. The control virus for this experiment, pAAV9-hSyn-mCherry was obtained (*Addgene, Cat#114472*) with a titer of 2.7x 10^13^ GC/ml. Viral particles were diluted to a final titer of ∼3×10^12^ GC/ml in sterile 1X PBS, along with Fast Green FCF dye and were subsequently injected into the striatum of P1 pups as described above. In our experiments, we utilized D2 *Foxp1^cKO^* and D2 *Foxp1^CTL^* mice where eGFP positive cells represented D2 SPNs while mCherry marked the cells expressing either control pAAV9-hSyn-mCherry (Control^AAV^) or pAAV9-hSyn-*Foxp1*-T2A-mCherry (*Foxp1*^AAV^) virus.

#### Immunoblotting from brain tissue

The striatum was dissected from mice and snap frozen in liquid nitrogen followed by storage at -80°C. For immunoblotting, tissue samples were homogenized using a Fisherbrand™ Pellet Pestle™ Cordless Motor (*Fisher Scientific, Cat#12-141-361*) in a lysis buffer composed of 20 mM Tris (pH 7.4), 150 mM NaCl, 2 mM EDTA, 1% NP-40, 0.1% SDS, 0.1% Sodium deoxycholate, 5% Glycerol, EDTA free protease inhibitor cocktail (*Sigma, Cat# 11873580001*), and phosphatase inhibitor mixture 2 (*Sigma, Cat#P5726*) and 3 (*Sigma, Cat#P0044*). Tissue lysates were further homogenized using brief 20 s pulses of sonication until clear lysates were obtained. Lysates were then centrifuged at 10,000 g for 10 min at 4°C, to remove the insoluble fraction. The resulting supernatant was used for protein quantification using a BCA protein assay kit (*Pierce, Cat#23227*). Equal amounts of protein were mixed with the 4X loading dye (40% Glycerol, 240 mM Tris pH 6.8, 8% SDS, 0.045 Bromophenol blue, 5% Beta-mercaptoethanol) and then incubated for 5 min at 95°C before being loaded on to a 10% SDS-PAGE. The subsequent steps of the immunoblotting procedure followed the same protocol as detailed above for HEK cells. To enable immunoblotting for multiple proteins on the same membrane, membranes were carefully cut at appropriate molecular weights. The primary antibodies were: FOXP1 (*in house*), mCherry (*Proteintech, Cat#26765-1*-AP), GAPDH (*Millipore*, *Cat# MAB374*), Actin (*Millipore, Cat#MAB1501*). Proteins of interest were quantified after normalization with GAPDH or Actin, which served as housekeeping proteins.

#### Immunohistochemistry

Mice aged one week or older were anesthetized with ketamine/xylazine and subsequently transcardially perfused with phosphate-buffered saline (PBS), followed by 4% paraformaldehyde (PFA) in PBS. Brains were then post-fixed overnight in 4% PFA. For mice younger than 1 week, perfusion was not conducted, and their brains were solely subjected to post-fixation. After post-fixation, brains were washed with PBS and then immersed in a 30% sucrose in PBS for 48 h followed by embedding in tissue plus OCT compound (*Fisher Health Care, Cat#4585*) and storing at -80°C. The brains were sectioned into 30 µm thick coronal sections using a cryostat (*Leica, Cat#CM1950*) at -20°C and collected in PBS. The sections containing the striatum were transferred onto Superfrost plus microscope slides (*Fisher Scientific, Cat#12550-15*) and allowed to dry. Subsequently, the sections were washed and permeabilized in 1X tris-buffered saline (TBS) followed by 1h incubation in a blocking solution (0.3M Glycine, 3% BSA and 10% normal donkey serum (NDS) in 1X TBS) at room temperature. Sections were again washed with TBS and incubated with primary antibodies (Rabbit anti-mCherry, *Proteintech, Cat#26765-1-AP*) in blocking solution (without Glycine) overnight at 4°C. Sections were then washed in TBS and incubated with secondary antibodies (Donkey anti-Rabbit IgG (H+L), Alexa Fluor^TM^ 555, *Invitrogen*, *Cat#A-31572*) prepared in 3% BSA in 1X TBS for 2 h at room temperature. The eGFP signal from *Drd2-eGFP* transgene did not require additional staining for eGFP and it was naturally detectable. Finally, following three more washes with 1X TBS, the sections were dried and mounted using Prolong Diamond Antifade with DAPI mountant (*Invitrogen, Cat#P36971*). Fluorescence images were acquired using a confocal laser scanning microscope (*Zeiss, LSM 880*) with a 20X objective.

### Statistics

For all datasets, the sample size is represented by the number of neurons, and the data are presented in the following order: WT control followed by *Foxp1* deletion groups. All data are presented as mean ± standard error. Unless otherwise specified, we conducted a repeated measures (RM) 2-way ANOVA (or Mixed Model) followed by the Holm-Sidak test for multiple comparison correction. In cases where multiple comparison correction was not applicable, we utilized Fisher’s LSD test. For the remaining datasets, we employed a t-test (if the data followed a normal distribution) or a Mann-Whitney (MW) test (for non-Gaussian distribution).

### Single-nuclei RNA-Sequencing (snRNA-Seq)

#### Tissue Collection and nuclei isolation

P18 mice were sacrificed by rapid decapitation and the brain was quickly dissected and transferred to ice-cold nACSF that contained (mM): 125 NaCl, 25 NaHCO_3_, 10 Dextrose, 3 KCl, 1.25 NaH_2_PO_4_, 2 CaCl_2_.2H_2_O, 2 MgCl_2_ with a pH of 7.3-7.4 and an osmolality of 300-305 mOsm. We placed the brain onto an ice-cold 1.0 mm stainless steel brain matrix (*Stoelting, Cat#51386*) and obtained three 1.0 mm thick striatal sections. These sections were then transferred into a petri-dish containing cold nACSF. Under a dissection microscope, we dissected the striatum from these slices using forceps. The isolated striatal tissue was flash frozen in liquid nitrogen and stored at -80^°^C. The striatal tissue was collected from mice of required genotypes at different time points.

Protocol for isolating nuclei was adapted and modified from our previously published work^8^. Briefly, tissue was homogenized 20 times with pestle A, followed by 20 times with pestle B, in a glass Dounce tissue grinder (*Sigma, Cat#D8938*) using 2 ml ice-cold EZ Nuclei Lysis Buffer (*Sigma, Cat #NUC-101*). The homogenized samples were then transferred into a 15 ml tube with an additional 2 ml of ice-cold EZ Nuclei Lysis Buffer, incubated on ice for 3 min and centrifuged at 500 X g for 5 min at 4^°^C. The supernatant was discarded, and the nuclei pellet was resuspended in 3 ml ice-cold Nuclei Suspension Buffer (NSB; 1X PBS, 0.01% Ultrapure BSA, and 0.1% RNase inhibitor) and centrifuged again at 500 x g for 5 min at 4^°^C. The supernatant was then discarded, and nuclei pellet was resuspended in 300-400 µl NSB, and filtered through 40 µm size Flowmi cell strainer (*Bel-Art Products, Cat# H13680-0040*). An aliquot of the filtered nuclei suspension was mixed with Trypan Blue and transferred into a disposable cell counting chamber nuclei to visualize and count the healthy nuclei under a light microscope ensuring a desired nuclei concentration of 700-1,200 nuclei/µl.

#### Preparation of cDNA libraries and sequencing

We prepared libraries from a total of 9 mice (3 mice/genotype) using the Chromium Next GEM Single Cell 3ʹ Reagent Kit v3.1 (*10X Genomics, Cat#1000121*) with a targeting goal of 10,000 nuclei per sample as per the manufacturer’s instructions. All mice received injections of either Control^AAV^ or *Foxp1*^AAV^ in their striatum at P1 where only a fraction of neurons was infected that were identified based on the presence of AAV transcripts. However, the relative proportion of AAV transcripts within the pool of total transcripts captured by snRNA-seq experiment was small. To address this, we used a modified approach ^81^ in which we specifically amplified the viral transcript sequences and prepared separate libraries. Briefly, we used 5% of the total cleaned up cDNA (amplified from the GEMs that were generated from each sample) and amplified viral transcripts using reverse primer: 10X_Enrich_R (5’CTACACGACGCTCTTCCG3’) and forward primers 10X_WPRE_F1 (5’ACGAGTCGGATCTCCCTTT3’, for Control^AAV^ infected mice) and 10X_mCherry_F1 (5’GCATGGACGAGCTGTACA3’. for *Foxp1*^AAV^ infected mice). We used different forward primers for Control^AAV^ and *Foxp1*^AAV^ infected mice since Control^AAV^ vector has a WPRE sequence while *Foxp1*^AAV^ has mCherry sequence at the 3’end (we removed WPRE sequence from the *Foxp1*^AAV^ vector to accommodate *Foxp1* and mCherry into the vector). The first round of PCR amplification was carried out for 25 cycles using Phusion High-Fidelity PCR Master Mix with HF Buffer (*NEB, Cat# M0531S*). The amplified PCR product was then run on an agarose gel, where a ∼250 bp product was excised from the gel and eluted using QIAquick Gel Extraction Kit (*Qiagen, Cat#28706*) in 20 µl elution buffer (*NEB*). 5% of this first round amplified product was used to perform a second round of amplification using the reverse primer 10X_WPRE_F1 and forward primers: 10X_WPRE_F2 (5’GTGACTGGAGTTCAGACGTGTGCTCTTCCGATCTACGAGTCGGATCTCCCTTT3’Control^AAV^) and 10X_mCherry_F2 (5’GTGACTGGAGTTCAGACGTGTGCTCTTCCGATCTGTACAAGTAGACCCAGCTT3’ *Foxp1*^AAV^) for 20 cycles using the same Phusion polymerase. Once again, this amplified PCR product was loaded on an agarose gel where similar size product (∼250 bp) was excised and eluted in 30 µl elution buffer. The eluted PCR products were cleaned up using SPRIselect reagent and subjected to sample-index PCR amplification according to the manufacturer’s instructions (10X Genomics) to generate cDNA libraries. Finally, both the total and AAV_transcript libraries were subjected to quality control assessments using the 4200 TapeStation System (*Agilent, Cat# G2991BA*) and quantified using a Qubit 2.0 Fluorometer (*Invitrogen*). We pooled 5 nM of the total libraries with 0.5 nM of the AAV_transcript libraries (10% of the total library) for each sample and sequenced on a NovaSeq 6000 (*Illumina*) platform using an S4 flow cell.

#### Pre-processing of sequencing data

We obtained raw sequencing data for both total and AAV_transcript libraries from the McDermott Sequencing Core at UT Southwestern in the form of binary base call (BCL) files. These BCL files were subsequently demultiplexed using 10X Genomics i7 index (used during library preparation) and Illumina’s bcl2fastq v2.19.1 and *mkfastq* command from 10X Genomics CellRanger v3.0.2 suite. Resulting FASTQ files were assessed for quality using FASTQC (v0.11.5) ^82^. To facilitate alignment and quantification, a reference mouse genome-annotation index was created using mouse genome (GRCm38p6) and Gencode annotation (vM17) with *mkref* command from 10X Genomics CellRanger v3.0.2 suite. Extracted and quality passed FASTQ reads for primary samples were then aligned to reference mouse genome-annotation index and raw count tables were generated using *count* command from 10X Genomics CellRanger v3.0.2 suite. CellBender’s ^83^ *remove-background* command was then run on the raw unfiltered count tables to discard potential ambient RNA. Also, potential doublets were identified with DoubletFinder^84^ utilizing filtered count tables generated by CellBender. This comprehensive pipeline produced an expression matrix with nuclei as columns and genes as rows, encompassing all primary samples and serving as the basis for subsequent analysis. A custom reference mouse genome-annotation index was separately constructed to include the amplified viral sequences along with mouse genome (GRCm38p6) and Gencode annotation (vM17) with *mkref* command from 10X Genomics CellRanger v3.0.2 suite. Extracted and quality passed FASTQ reads for these samples with amplified viral transcript specific sequences were then aligned to the custom reference mouse genome-annotation index and raw count tables for amplified viral transcript sequences were generated using *count* command from 10X Genomics CellRanger v3.0.2 suite.

#### Clustering analysis

Cleaned count tables for primary samples were used to run Seurat pipeline following the vignette (https://satijalab.org/seurat/articles/pbmc3k_tutorial.html). Samples across genotypes were processed through Harmony ^85^ using command *RunHarmony* to remove the effects of covariates such as batch and sex. Harmonized Seurat objects were then integrated following vignette (https://portals.broadinstitute.org/harmony/SeuratV3.html). Integrated data was clustered to identify groups of nuclei by cell-types using the top 59 principal components identified using *JackStraw* and *ScoreJackStraw* approach within Seurat. Resolution of 1.2 was selected using *Clustree* (https://github.com/lazappi/clustree) approach. *FindAllMarkers* was then run to identify gene markers corresponding to each identified cluster. Cell type annotation was executed utilizing known marker genes and by performing a Fisher exact test against cell type marker genes previously identified in a reference dataset ^86^. Furthermore, nuclei corresponding to spiny projection neuron (SPN) classes were sub-clustered to resolve specific SPN sub-types (D1, D2, and eccentric SPNs) with higher granularity. Cell-barcodes that belong to SPN classes were also marked for presence of viral amplified sequences with at least six UMIs per cell-barcode from samples with viral amplified sequences. Specifically, we designated any D2 SPN that exhibited at least six or more WPRE/mCherry UMIs as Control^AAV^/*Foxp1*^AAV^ -positive D2 SPN respectively (Fig. 16A, B).

#### Differential gene expression (DGE) analysis

For the analysis of differentially expressed genes (DEGs), we focused on nuclei corresponding to the D2-SPN class and grouped by genotype. To identify genes with significant differential expression within D2-SPNs in D2 *Foxp1^cKO^*mice compared to D2 *Foxp1^CTL^* mice, we utilized edgeR ^87^-based pseudobulk approach ^88^. The cut off for significance was established as a false discovery rate (FDR) < 0.05 and log2 fold change ≥ 0.378. Similarly, we also identified DEGs for AAV-infected D2 SPNs that contain a minimum of six UMIs per cell-barcode for viral amplified sequences, within the same genotype groups. The number of AAV-infected D2 SPNs exhibited variability across different samples. Thus, to impart a stringent cutoff to exclude false positives, we established a threshold of six UMIs to exclude at least 50% of D2 SPNs in each sample and included the remaining D2 SPNs, with the greatest number of viral UMI’s, in our analysis. We compared AAV-infected D2 SPNs in D2 Foxp1cKO + Foxp1AAV with those in D2 Foxp1cKO + ControlAAV and identified DEGs. Gene transcripts originating from mitochondrial (chr M), and sex chromosomes (chr X and chr Y) were excluded from the DGE output, given that the samples encompassed mixed genders.

#### Gene ontology analysis

For the analysis of gene ontology associated with DEGs, we employed *ClusterProfiler* ^89^. Redundant ontology terms were subsequently removed using *Revigo* ^90^. Additionally, we conducted synaptic gene ontology analysis using *SynGO* (https://www.syngoportal.org) ^91^ to annotate significant DEGs across different comparisons. All expressed genes within their respective groups were used as background for the ontological analyses.

### Behavior

#### Nest Building

Mice were separated and single housed for 24 h in new cages with intact nestlet material. Nests were scored in adult mice for their quality based on previously established criteria ^8^ on a scale from 1-5 as previously described ^92^.

#### Open Field Test

This test was performed by the behavior core at UT Southwestern. Mice were individually placed in a 16’’ X 16’’ Plexiglass box and allowed to explore the arena for 10 min. Total distance moved and the amount of time spent in the periphery was measured using *EthoVision XT* software package (*Noldus*). Average velocity was determined by dividing the total distance by the amount of time spent in the box (600 s).

#### Rotarod Test

Rotarod test was performed as described earlier ^8, 17^. Mice were marked on their tails and habituated for 30 min before being placed individually on a textured rod with lanes of a Series 8 IITC Life Science Rotarod. The rod was programmed to accelerate from 4 to 40 rpm over a 5 min period. Mice were positioned facing forward on the rod and sensors beneath the rod were activated when a mouse fell off. If a mouse managed to make one full rotation while holding onto the rod, the sensor was manually triggered, and the mouse was removed from the rod. Mice were tested for three consecutive days, with four trials each day and at least a 10 min interval between trials and latency to fall was recorded for each trial.

**<u>Table S1.</u> Differentially Expressed Genes in D2 SPNs, related to Figure 7**

Differentially expressed genes in D2 SPNs where columns show gene name, log fold change, log CPM, P value and FDR. Tab 1 shows DEGs in “all” D2 SPNs from D2 *Foxp1^CTL^* mice and D2 *Foxp1^cKO^* mice while Tabs 2-4 show DEGs in “selected” D2 SPNs from D2 *Foxp1^CTL^* mice infected with Control^AAV^ and from D2 *Foxp1^cKO^* mice infected with Control^AAV^ and *Foxp1*^AAV^.

**<u>Table S2.</u> GO terms associated with the DEGs.**

GO terms where columns show GO IDs, gene ratio, description, p-value, adjusted p-value, q-value, etc., generated using *Clusterprofiler*. Tabs 1, 2 show gene ontology terms for upregulated and downregulated DEGs in D2 *Foxp1^cKO^* mice in “all” D2 SPNs as compared to D2 *Foxp1^CTL^* mice. Tabs 3, 4 show gene ontology terms for upregulated and downregulated DEGs in D2 *Foxp1^cKO^* mice in D2 SPNs infected with *Foxp1*^AAV^ as compared to the D2 SPNs infected with Control^AAV^.

**<u>Table S3.</u> SynGO terms associated with the DEGs.**

GO terms where columns show GO term name, hierarchical structure, p-value, etc., generated using *SynGO*. Tabs 1, 2 show synaptic gene ontology terms for upregulated and downregulated DEGs in D2 *Foxp1^cKO^* mice in “all” D2 SPNs as compared to D2 *Foxp1^CTL^* mice. Tabs 3, 4 show gene ontology terms for upregulated and downregulated DEGs in D2 *Foxp1^cKO^* mice in D2 SPNs infected with *Foxp1*^AAV^ as compared to the D2 SPNs infected with Control^AAV^.

## Key Resources Table

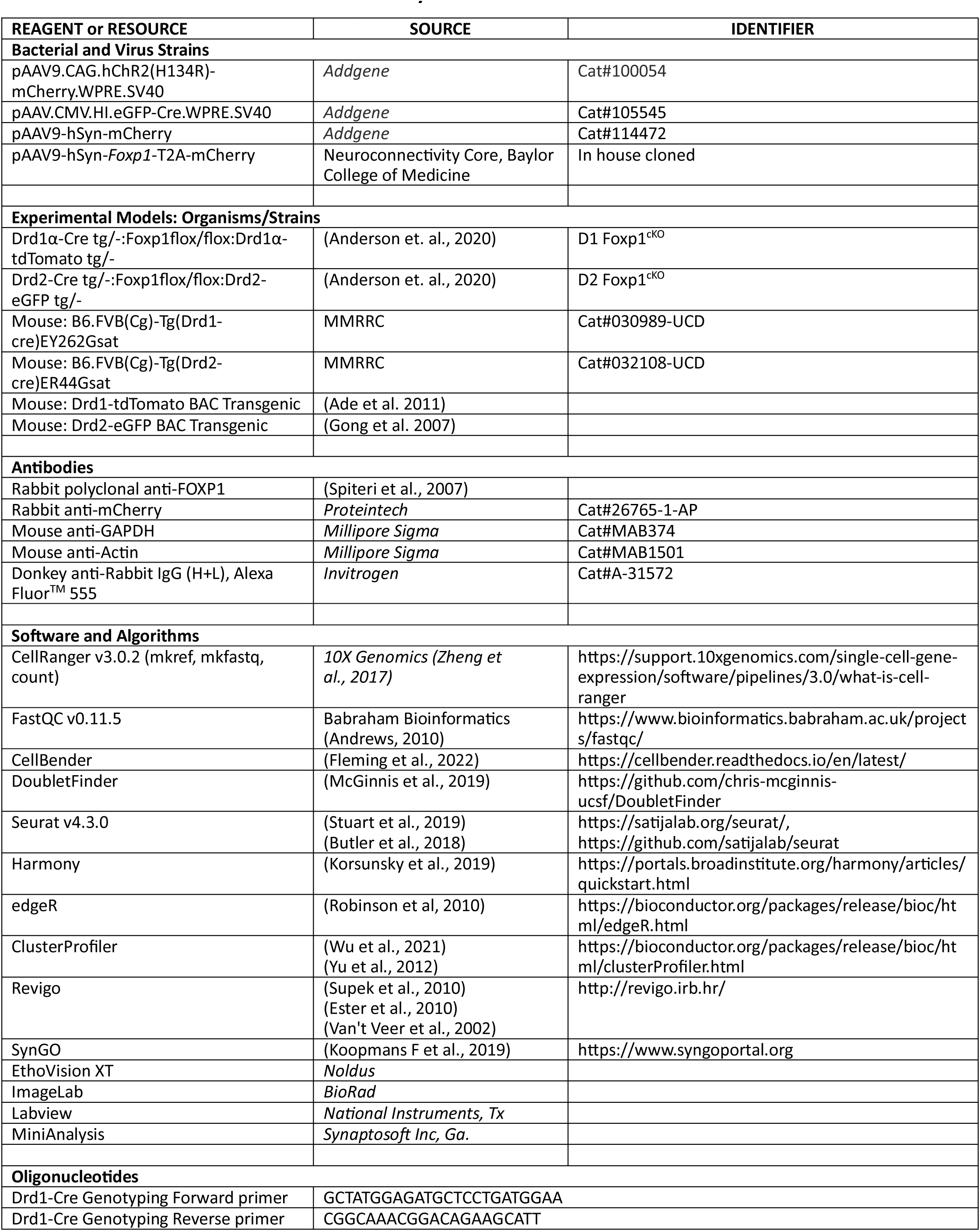
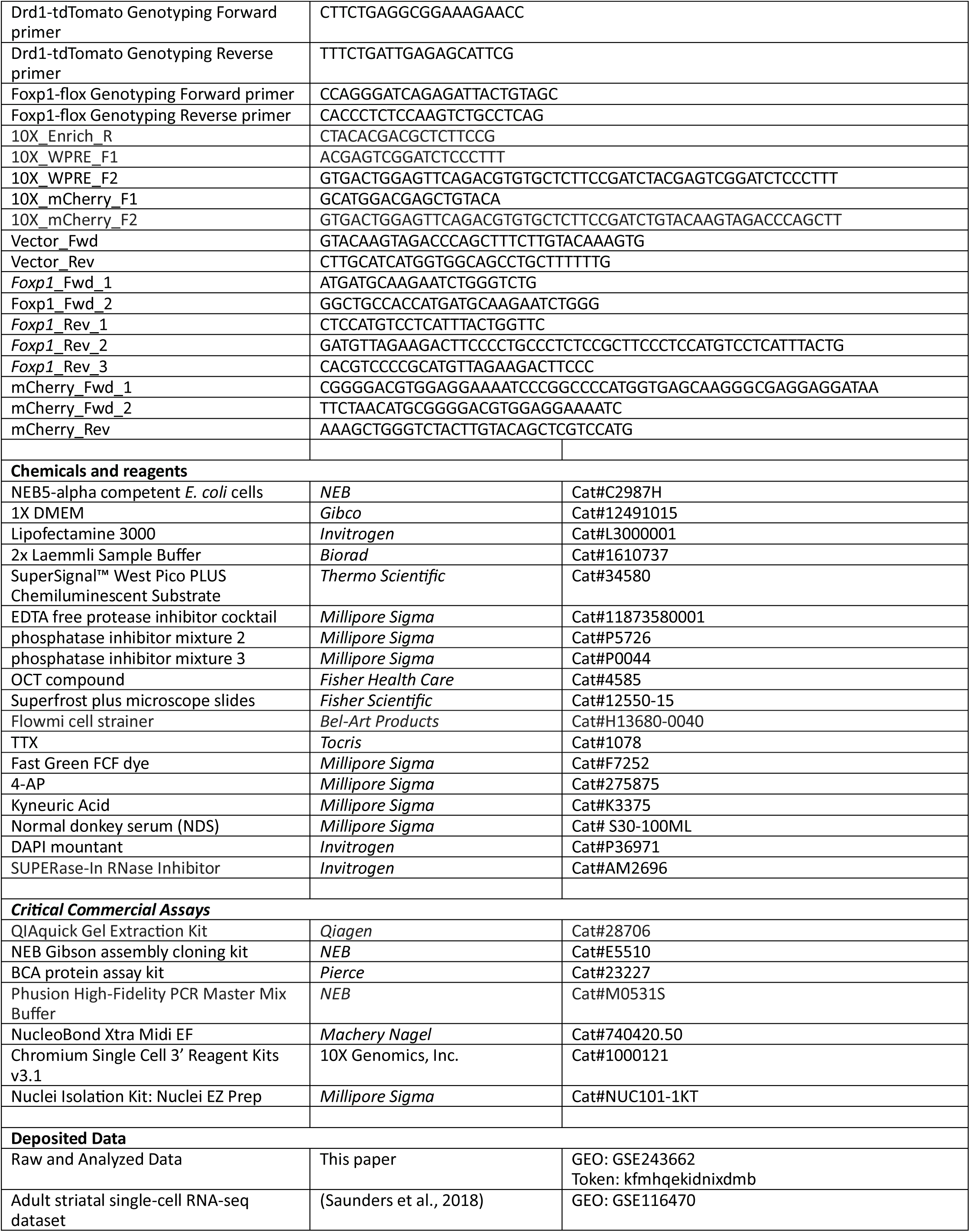

## Notes

### Competing Interest Statement

The authors have declared no competing interest.

### Summary of Updates

This version has been submitted to revise the page numbers of the figure file.

## References

1. Haber SN. Corticostriatal circuitry. Dialogues Clin Neurosci. 2016;18(1):7–21. doi: 10.31887/DCNS.2016.18.1/shaber. PubMed PMID: 27069376; PMCID: PMC4826773.

2. Lieberman OJ, McGuirt AF, Mosharov EV, Pigulevskiy I, Hobson BD, Choi S, Frier MD, Santini E, Borgkvist A, Sulzer D. Dopamine Triggers the Maturation of Striatal Spiny Projection Neuron Excitability during a Critical Period. Neuron. 2018;99(3):540–54 e4. doi: 10.1016/j.neuron.2018.06.044. PubMed PMID: 30057204.

3. Kozorovitskiy Y, Saunders A, Johnson CA, Lowell BB, Sabatini BL. Recurrent network activity drives striatal synaptogenesis. Nature. 2012;485(7400):646–50. Epub 20120513. doi: 10.1038/nature11052. PubMed PMID: 22660328; PMCID: PMC3367801.

4. Kozorovitskiy Y, Peixoto R, Wang W, Saunders A, Sabatini BL. Neuromodulation of excitatory synaptogenesis in striatal development. Elife. 2015;4. Epub 20151109. doi: 10.7554/eLife.10111. PubMed PMID: 26551563; PMCID: PMC4716836.

5. Li W, Pozzo-Miller L. Dysfunction of the corticostriatal pathway in autism spectrum disorders. J Neurosci Res. 2020;98(11):2130–47. Epub 2019/11/24. doi: 10.1002/jnr.24560. PubMed PMID: 31758607; PMCID: PMC7242149.

6. Yang L, Su Z, Wang Z, Li Z, Shang Z, Du H, Liu G, Qi D, Yang Z, Xu Z, Zhang Z. Transcriptional profiling reveals the transcription factor networks regulating the survival of striatal neurons. Cell Death Dis. 2021;12(3):262. Epub 20210312. doi: 10.1038/s41419-021-03552-8. PubMed PMID: 33712552; PMCID: PMC7955055.

7. Arlotta P, Molyneaux BJ, Jabaudon D, Yoshida Y, Macklis JD. Ctip2 controls the differentiation of medium spiny neurons and the establishment of the cellular architecture of the striatum. J Neurosci. 2008;28(3):622–32. doi: 10.1523/JNEUROSCI.2986-07.2008. PubMed PMID: 18199763; PMCID: PMC6670353.

8. Anderson AG, Kulkarni A, Harper M, Konopka G. Single-Cell Analysis of Foxp1-Driven Mechanisms Essential for Striatal Development. Cell Rep. 2020;30(9):3051–66 e7. doi: 10.1016/j.celrep.2020.02.030. PubMed PMID: 32130906; PMCID: PMC7137930.

9. Zhang Q, Zhang Y, Wang C, Xu Z, Liang Q, An L, Li J, Liu Z, You Y, He M, Mao Y, Chen B, Xiong ZQ, Rubenstein JL, Yang Z. The Zinc Finger Transcription Factor Sp9 Is Required for the Development of Striatopallidal Projection Neurons. Cell Rep. 2016;16(5):1431–44. Epub 20160721. doi: 10.1016/j.celrep.2016.06.090. PubMed PMID: 27452460; PMCID: PMC4972643.

10. Chen YC, Kuo HY, Bornschein U, Takahashi H, Chen SY, Lu KM, Yang HY, Chen GM, Lin JR, Lee YH, Chou YC, Cheng SJ, Chien CT, Enard W, Hevers W, Paabo S, Graybiel AM, Liu FC. Foxp2 controls synaptic wiring of corticostriatal circuits and vocal communication by opposing Mef2c. Nat Neurosci. 2016;19(11):1513–22. doi: 10.1038/nn.4380. PubMed PMID: 27595386; PMCID: PMC5083203.

11. Ferland RJ, Cherry TJ, Preware PO, Morrisey EE, Walsh CA. Characterization of Foxp2 and Foxp1 mRNA and protein in the developing and mature brain. J Comp Neurol. 2003;460(2):266–79. doi: 10.1002/cne.10654. PubMed PMID: 12687690.

12. Siper PM, De Rubeis S, Trelles MDP, Durkin A, Di Marino D, Muratet F, Frank Y, Lozano R, Eichler EE, Kelly M, Beighley J, Gerdts J, Wallace AS, Mefford HC, Bernier RA, Kolevzon A, Buxbaum JD. Prospective investigation of FOXP1 syndrome. Mol Autism. 2017;8:57. doi: 10.1186/s13229-017-0172-6. PubMed PMID: 29090079; PMCID: PMC5655854.

13. Horn D, Kapeller J, Rivera-Brugues N, Moog U, Lorenz-Depiereux B, Eck S, Hempel M, Wagenstaller J, Gawthrope A, Monaco AP, Bonin M, Riess O, Wohlleber E, Illig T, Bezzina CR, Franke A, Spranger S, Villavicencio-Lorini P, Seifert W, Rosenfeld J, Klopocki E, Rappold GA, Strom TM. Identification of FOXP1 deletions in three unrelated patients with mental retardation and significant speech and language deficits. Hum Mutat. 2010;31(11):E1851-60. doi: 10.1002/humu.21362. PubMed PMID: 20848658; PMCID: PMC3049153.

14. Iossifov I, O’Roak BJ, Sanders SJ, Ronemus M, Krumm N, Levy D, Stessman HA, Witherspoon KT, Vives L, Patterson KE, Smith JD, Paeper B, Nickerson DA, Dea J, Dong S, Gonzalez LE, Mandell JD, Mane SM, Murtha MT, Sullivan CA, Walker MF, Waqar Z, Wei L, Willsey AJ, Yamrom B, Lee YH, Grabowska E, Dalkic E, Wang Z, Marks S, Andrews P, Leotta A, Kendall J, Hakker I, Rosenbaum J, Ma B, Rodgers L, Troge J, Narzisi G, Yoon S, Schatz MC, Ye K, McCombie WR, Shendure J, Eichler EE, State MW, Wigler M. The contribution of de novo coding mutations to autism spectrum disorder. Nature. 2014;515(7526):216–21. doi: 10.1038/nature13908. PubMed PMID: 25363768.

15. Sanders SJ, He X, Willsey AJ, Ercan-Sencicek AG, Samocha KE, Cicek AE, Murtha MT, Bal VH, Bishop SL, Dong S, Goldberg AP, Jinlu C, Keaney JF, 3rd, Klei L, Mandell JD, Moreno-De-Luca D, Poultney CS, Robinson EB, Smith L, Solli-Nowlan T, Su MY, Teran NA, Walker MF, Werling DM, Beaudet AL, Cantor RM, Fombonne E, Geschwind DH, Grice DE, Lord C, Lowe JK, Mane SM, Martin DM, Morrow EM, Talkowski ME, Sutcliffe JS, Walsh CA, Yu TW, Autism Sequencing C, Ledbetter DH, Martin CL, Cook EH, Buxbaum JD, Daly MJ, Devlin B, Roeder K, State MW. Insights into Autism Spectrum Disorder Genomic Architecture and Biology from 71 Risk Loci. Neuron. 2015;87(6):1215–33. doi: 10.1016/j.neuron.2015.09.016. PubMed PMID: 26402605; PMCID: PMC4624267.

16. Stessman HA, Xiong B, Coe BP, Wang T, Hoekzema K, Fenckova M, Kvarnung M, Gerdts J, Trinh S, Cosemans N, Vives L, Lin J, Turner TN, Santen G, Ruivenkamp C, Kriek M, van Haeringen A, Aten E, Friend K, Liebelt J, Barnett C, Haan E, Shaw M, Gecz J, Anderlid BM, Nordgren A, Lindstrand A, Schwartz C, Kooy RF, Vandeweyer G, Helsmoortel C, Romano C, Alberti A, Vinci M, Avola E, Giusto S, Courchesne E, Pramparo T, Pierce K, Nalabolu S, Amaral DG, Scheffer IE, Delatycki MB, Lockhart PJ, Hormozdiari F, Harich B, Castells-Nobau A, Xia K, Peeters H, Nordenskjold M, Schenck A, Bernier RA, Eichler EE. Targeted sequencing identifies 91 neurodevelopmental-disorder risk genes with autism and developmental-disability biases. Nat Genet. 2017;49(4):515–26. doi: 10.1038/ng.3792. PubMed PMID: 28191889; PMCID: PMC5374041.

17. Araujo DJ, Anderson AG, Berto S, Runnels W, Harper M, Ammanuel S, Rieger MA, Huang HC, Rajkovich K, Loerwald KW, Dekker JD, Tucker HO, Dougherty JD, Gibson JR, Konopka G. FoxP1 orchestration of ASD-relevant signaling pathways in the striatum. Genes Dev. 2015;29(20):2081–96. doi: 10.1101/gad.267989.115. PubMed PMID: 26494785; PMCID: PMC4617974.

18. Bacon C, Schneider M, Le Magueresse C, Froehlich H, Sticht C, Gluch C, Monyer H, Rappold GA. Brain-specific Foxp1 deletion impairs neuronal development and causes autistic-like behaviour. Mol Psychiatry. 2015;20(5):632–9. Epub 20140930. doi: 10.1038/mp.2014.116. PubMed PMID: 25266127; PMCID: PMC4419151.

19. Khandelwal N, Cavalier S, Rybalchenko V, Kulkarni A, Anderson AG, Konopka G, Gibson JR. FOXP1 negatively regulates intrinsic excitability in D2 striatal projection neurons by promoting inwardly rectifying and leak potassium currents. Mol Psychiatry. 2021;26(6):1761–74. Epub 20210105. doi: 10.1038/s41380-020-00995-x. PubMed PMID: 33402705; PMCID: PMC8255328.

20. Knowles R, Dehorter N, Ellender T. From Progenitors to Progeny: Shaping Striatal Circuit Development and Function. J Neurosci. 2021;41(46):9483–502. doi: 10.1523/JNEUROSCI.0620-21.2021. PubMed PMID: 34789560; PMCID: PMC8612473.

21. Araujo DJ, Toriumi K, Escamilla CO, Kulkarni A, Anderson AG, Harper M, Usui N, Ellegood J, Lerch JP, Birnbaum SG, Tucker HO, Powell CM, Konopka G. Foxp1 in Forebrain Pyramidal Neurons Controls Gene Expression Required for Spatial Learning and Synaptic Plasticity. J Neurosci. 2017;37(45):10917–31. Epub 20171004. doi: 10.1523/JNEUROSCI.1005-17.2017. PubMed PMID: 28978667; PMCID: PMC5678023.

22. Garcia-Oscos F, Koch TMI, Pancholi H, Trusel M, Daliparthi V, Co M, Park SE, Ayhan F, Alam DH, Holdway JE, Konopka G, Roberts TF. Autism-linked gene FoxP1 selectively regulates the cultural transmission of learned vocalizations. Sci Adv. 2021;7(6). Epub 20210203. doi: 10.1126/sciadv.abd2827. PubMed PMID: 33536209; PMCID: PMC7857683.

23. Zhai S, Tanimura A, Graves SM, Shen W, Surmeier DJ. Striatal synapses, circuits, and Parkinson’s disease. Curr Opin Neurobiol. 2018;48:9–16. Epub 20170824. doi: 10.1016/j.conb.2017.08.004. PubMed PMID: 28843800; PMCID: PMC6022405.

24. Kim H, Gao EB, Draper A, Berens NC, Vihma H, Zhang X, Higashi-Howard A, Ritola KD, Simon JM, Kennedy AJ, Philpot BD. Rescue of behavioral and electrophysiological phenotypes in a Pitt-Hopkins syndrome mouse model by genetic restoration of Tcf4 expression. Elife. 2022;11. Epub 20220510. doi: 10.7554/eLife.72290. PubMed PMID: 35535852; PMCID: PMC9090324.

25. Shepherd GM. Corticostriatal connectivity and its role in disease. Nat Rev Neurosci. 2013;14(4):278–91. doi: 10.1038/nrn3469. PubMed PMID: 23511908; PMCID: PMC4096337.

26. Kress GJ, Yamawaki N, Wokosin DL, Wickersham IR, Shepherd GM, Surmeier DJ. Convergent cortical innervation of striatal projection neurons. Nat Neurosci. 2013;16(6):665–7. Epub 20130512. doi: 10.1038/nn.3397. PubMed PMID: 23666180; PMCID: PMC4085670.

27. Huang YH, Schluter OM, Dong Y. Cocaine-induced homeostatic regulation and dysregulation of nucleus accumbens neurons. Behav Brain Res. 2011;216(1):9–18. Epub 20100811. doi: 10.1016/j.bbr.2010.07.039. PubMed PMID: 20708038; PMCID: PMC2975799.

28. Petreanu L, Mao T, Sternson SM, Svoboda K. The subcellular organization of neocortical excitatory connections. Nature. 2009;457(7233):1142–5. Epub 2009/01/20. doi: nature07709 [pii]10.1038/nature07709 [doi]. PubMed PMID: 19151697; PMCID: 2745650.

29. Krajeski RN, Macey-Dare A, van Heusden F, Ebrahimjee F, Ellender TJ. Dynamic postnatal development of the cellular and circuit properties of striatal D1 and D2 spiny projection neurons. J Physiol. 2019;597(21):5265–93. Epub 20191010. doi: 10.1113/JP278416. PubMed PMID: 31531863; PMCID: PMC6900874.

30. Bacon C, Rappold GA. The distinct and overlapping phenotypic spectra of FOXP1 and FOXP2 in cognitive disorders. Hum Genet. 2012;131(11):1687–98. Epub 20120627. doi: 10.1007/s00439-012-1193-z. PubMed PMID: 22736078; PMCID: PMC3470686.

31. Rajkovich KE, Loerwald KW, Hale CF, Hess CT, Gibson JR, Huber KM. Experience-Dependent and Differential Regulation of Local and Long-Range Excitatory Neocortical Circuits by Postsynaptic Mef2c. Neuron. 2017;93(1):48–56. doi: 10.1016/j.neuron.2016.11.022. PubMed PMID: 27989458; PMCID: PMC5215966.

32. Adesnik H, Li G, During MJ, Pleasure SJ, Nicoll RA. NMDA receptors inhibit synapse unsilencing during brain development. Proc Natl Acad Sci U S A. 2008;105(14):5597–602. Epub 2008/04/01. doi: 10.1073/pnas.0800946105. PubMed PMID: 18375768; PMCID: PMC2291097.

33. Feliciangeli S, Chatelain FC, Bichet D, Lesage F. The family of K2P channels: salient structural and functional properties. J Physiol. 2015;593(12):2587–603. Epub 20150122. doi: 10.1113/jphysiol.2014.287268. PubMed PMID: 25530075; PMCID: PMC4500345.

34. Mermelstein PG, Song WJ, Tkatch T, Yan Z, Surmeier DJ. Inwardly rectifying potassium (IRK) currents are correlated with IRK subunit expression in rat nucleus accumbens medium spiny neurons. J Neurosci. 1998;18(17):6650–61. doi: 10.1523/JNEUROSCI.18-17-06650.1998. PubMed PMID: 9712637; PMCID: PMC6792959.

35. Nisenbaum ES, Xu ZC, Wilson CJ. Contribution of a slowly inactivating potassium current to the transition to firing of neostriatal spiny projection neurons. J Neurophysiol. 1994;71(3):1174–89. doi: 10.1152/jn.1994.71.3.1174. PubMed PMID: 8201411.

36. Wilson CJ, Kawaguchi Y. The origins of two-state spontaneous membrane potential fluctuations of neostriatal spiny neurons. J Neurosci. 1996;16(7):2397–410. doi: 10.1523/JNEUROSCI.16-07-02397.1996. PubMed PMID: 8601819; PMCID: PMC6578540.

37. Prybylowski K, Wenthold RJ. N-Methyl-D-aspartate receptors: subunit assembly and trafficking to the synapse. J Biol Chem. 2004;279(11):9673–6. Epub 20040123. doi: 10.1074/jbc.R300029200. PubMed PMID: 14742424.

38. Malinow R, Malenka RC. AMPA receptor trafficking and synaptic plasticity. Annu Rev Neurosci. 2002;25:103–26. Epub 20020304. doi: 10.1146/annurev.neuro.25.112701.142758. PubMed PMID: 12052905.

39. Nicoll RA, Tomita S, Bredt DS. Auxiliary subunits assist AMPA-type glutamate receptors. Science. 2006;311(5765):1253–6. doi: 10.1126/science.1123339. PubMed PMID: 16513974.

40. Wan Y, Ade KK, Caffall Z, Ilcim Ozlu M, Eroglu C, Feng G, Calakos N. Circuit-selective striatal synaptic dysfunction in the Sapap3 knockout mouse model of obsessive-compulsive disorder. Biol Psychiatry. 2014;75(8):623–30. Epub 20130213. doi: 10.1016/j.biopsych.2013.01.008. PubMed PMID: 23414593; PMCID: PMC3687030.

41. Wan Y, Feng G, Calakos N. Sapap3 deletion causes mGluR5-dependent silencing of AMPAR synapses. J Neurosci. 2011;31(46):16685–91. doi: 10.1523/JNEUROSCI.2533-11.2011. PubMed PMID: 22090495; PMCID: PMC3475185.

42. Welch JM, Lu J, Rodriguiz RM, Trotta NC, Peca J, Ding JD, Feliciano C, Chen M, Adams JP, Luo J, Dudek SM, Weinberg RJ, Calakos N, Wetsel WC, Feng G. Cortico-striatal synaptic defects and OCD-like behaviours in Sapap3-mutant mice. Nature. 2007;448(7156):894–900. doi: 10.1038/nature06104. PubMed PMID: 17713528; PMCID: PMC2442572.

43. Weeber EJ, Beffert U, Jones C, Christian JM, Forster E, Sweatt JD, Herz J. Reelin and ApoE receptors cooperate to enhance hippocampal synaptic plasticity and learning. J Biol Chem. 2002;277(42):39944–52. Epub 20020807. doi: 10.1074/jbc.M205147200. PubMed PMID: 12167620.

44. Bateup HS, Santini E, Shen W, Birnbaum S, Valjent E, Surmeier DJ, Fisone G, Nestler EJ, Greengard P. Distinct subclasses of medium spiny neurons differentially regulate striatal motor behaviors. Proc Natl Acad Sci U S A. 2010;107(33):14845–50. Epub 20100803. doi: 10.1073/pnas.1009874107. PubMed PMID: 20682746; PMCID: PMC2930415.

45. Calabresi P, Gubellini P, Centonze D, Picconi B, Bernardi G, Chergui K, Svenningsson P, Fienberg AA, Greengard P. Dopamine and cAMP-regulated phosphoprotein 32 kDa controls both striatal long-term depression and long-term potentiation, opposing forms of synaptic plasticity. J Neurosci. 2000;20(22):8443–51. doi: 10.1523/JNEUROSCI.20-22-08443.2000. PubMed PMID: 11069952; PMCID: PMC6773171.

46. Blank T, Nijholt I, Teichert U, Kugler H, Behrsing H, Fienberg A, Greengard P, Spiess J. The phosphoprotein DARPP-32 mediates cAMP-dependent potentiation of striatal N-methyl-D-aspartate responses. Proc Natl Acad Sci U S A. 1997;94(26):14859–64. doi: 10.1073/pnas.94.26.14859. PubMed PMID: 9405704; PMCID: PMC25128.

47. Chen C, Soto G, Dumrongprechachan V, Bannon N, Kang S, Kozorovitskiy Y, Parisiadou L. Pathway-specific dysregulation of striatal excitatory synapses by LRRK2 mutations. Elife. 2020;9. Epub 20201002. doi: 10.7554/eLife.58997. PubMed PMID: 33006315; PMCID: PMC7609054.

48. Giesert F, Hofmann A, Burger A, Zerle J, Kloos K, Hafen U, Ernst L, Zhang J, Vogt-Weisenhorn DM, Wurst W. Expression analysis of Lrrk1, Lrrk2 and Lrrk2 splice variants in mice. PLoS One. 2013;8(5):e63778. Epub 20130510. doi: 10.1371/journal.pone.0063778. PubMed PMID: 23675505; PMCID: PMC3651128.

49. Matikainen-Ankney BA, Kezunovic N, Mesias RE, Tian Y, Williams FM, Huntley GW, Benson DL. Altered Development of Synapse Structure and Function in Striatum Caused by Parkinson’s Disease-Linked LRRK2-G2019S Mutation. J Neurosci. 2016;36(27):7128–41. doi: 10.1523/JNEUROSCI.3314-15.2016. PubMed PMID: 27383589; PMCID: PMC4938860.

50. Jitsuki-Takahashi A, Jitsuki S, Yamashita N, Kawamura M, Abe M, Sakimura K, Sano A, Nakamura F, Goshima Y, Takahashi T. Activity-induced secretion of semaphorin 3A mediates learning. Eur J Neurosci. 2021;53(10):3279–93. Epub 20210405. doi: 10.1111/ejn.15210. PubMed PMID: 33772906; PMCID: PMC8252788.

51. Tepper JM, Sharpe NA, Koos TZ, Trent F. Postnatal development of the rat neostriatum: electrophysiological, light- and electron-microscopic studies. Dev Neurosci. 1998;20(2-3):125–45. doi: 10.1159/000017308. PubMed PMID: 9691188.

52. Pratt KG, Aizenman CD. Homeostatic regulation of intrinsic excitability and synaptic transmission in a developing visual circuit. J Neurosci. 2007;27(31):8268–77. doi: 10.1523/JNEUROSCI.1738-07.2007. PubMed PMID: 17670973; PMCID: PMC6673059.

53. Sim S, Antolin S, Lin CW, Lin Y, Lois C. Increased cell-intrinsic excitability induces synaptic changes in new neurons in the adult dentate gyrus that require Npas4. J Neurosci. 2013;33(18):7928–40. doi: 10.1523/JNEUROSCI.1571-12.2013. PubMed PMID: 23637184; PMCID: PMC3853377.

54. Corbett CM, Miller END, Wannen EE, Rood BD, Chandler DJ, Loweth JA. Cocaine Exposure Increases Excitatory Synaptic Transmission and Intrinsic Excitability in the Basolateral Amygdala in Male and Female Rats and Across the Estrous Cycle. Neuroendocrinology. 2023. Epub 20230602. doi: 10.1159/000531351. PubMed PMID: 37271140.

55. Fernandez-Perez EJ, Gallegos S, Armijo-Weingart L, Araya A, Riffo-Lepe NO, Cayuman F, Aguayo LG. Changes in neuronal excitability and synaptic transmission in nucleus accumbens in a transgenic Alzheimer’s disease mouse model. Sci Rep. 2020;10(1):19606. Epub 20201111. doi: 10.1038/s41598-020-76456-w. PubMed PMID: 33177601; PMCID: PMC7659319.

56. Turrigiano GG, Nelson SB. Homeostatic plasticity in the developing nervous system. Nat Rev Neurosci. 2004;5(2):97–107. doi: 10.1038/nrn1327. PubMed PMID: 14735113.

57. Braccioli L, Vervoort SJ, Adolfs Y, Heijnen CJ, Basak O, Pasterkamp RJ, Nijboer CH, Coffer PJ. FOXP1 Promotes Embryonic Neural Stem Cell Differentiation by Repressing Jagged1 Expression. Stem Cell Reports. 2017;9(5):1530–45. doi: 10.1016/j.stemcr.2017.10.012. PubMed PMID: 29141232; PMCID: PMC5688236.

58. Pearson CA, Moore DM, Tucker HO, Dekker JD, Hu H, Miquelajauregui A, Novitch BG. Foxp1 Regulates Neural Stem Cell Self-Renewal and Bias Toward Deep Layer Cortical Fates. Cell Rep. 2020;30(6):1964–81 e3. doi: 10.1016/j.celrep.2020.01.034. PubMed PMID: 32049024; PMCID: PMC8397815.

59. Konstantoulas CJ, Parmar M, Li M. FoxP1 promotes midbrain identity in embryonic stem cell-derived dopamine neurons by regulating Pitx3. J Neurochem. 2010;113(4):836–47. Epub 20100219. doi: 10.1111/j.1471-4159.2010.06650.x. PubMed PMID: 20175877.

60. Li X, Han X, Tu X, Zhu D, Feng Y, Jiang T, Yang Y, Qu J, Chen JG. An Autism-Related, Nonsense Foxp1 Mutant Induces Autophagy and Delays Radial Migration of the Cortical Neurons. Cereb Cortex. 2019;29(7):3193–208. doi: 10.1093/cercor/bhy185. PubMed PMID: 30124790.

61. Li X, Xiao J, Frohlich H, Tu X, Li L, Xu Y, Cao H, Qu J, Rappold GA, Chen JG. Foxp1 regulates cortical radial migration and neuronal morphogenesis in developing cerebral cortex. PLoS One. 2015;10(5):e0127671. Epub 20150526. doi: 10.1371/journal.pone.0127671. PubMed PMID: 26010426; PMCID: PMC4444005.

62. Palmesino E, Rousso DL, Kao TJ, Klar A, Laufer E, Uemura O, Okamoto H, Novitch BG, Kania A. Foxp1 and lhx1 coordinate motor neuron migration with axon trajectory choice by gating Reelin signalling. PLoS Biol. 2010;8(8):e1000446. Epub 20100810. doi: 10.1371/journal.pbio.1000446. PubMed PMID: 20711475; PMCID: PMC2919418.

63. Wang J, Frohlich H, Torres FB, Silva RL, Poschet G, Agarwal A, Rappold GA. Mitochondrial dysfunction and oxidative stress contribute to cognitive and motor impairment in FOXP1 syndrome. Proc Natl Acad Sci U S A. 2022;119(8). doi: 10.1073/pnas.2112852119. PubMed PMID: 35165191; PMCID: PMC8872729.

64. Wang J, Rappold GA, Frohlich H. Disrupted Mitochondrial Network Drives Deficits of Learning and Memory in a Mouse Model of FOXP1 Haploinsufficiency. Genes (Basel). 2022;13(1). Epub 20220111. doi: 10.3390/genes13010127. PubMed PMID: 35052467; PMCID: PMC8775322.

65. Pisani A, Calabresi P, Centonze D, Bernardi G. Enhancement of NMDA responses by group I metabotropic glutamate receptor activation in striatal neurones. Br J Pharmacol. 1997;120(6):1007–14. doi: 10.1038/sj.bjp.0700999. PubMed PMID: 9134210; PMCID: PMC1564563.

66. Pisani A, Gubellini P, Bonsi P, Conquet F, Picconi B, Centonze D, Bernardi G, Calabresi P. Metabotropic glutamate receptor 5 mediates the potentiation of N-methyl-D-aspartate responses in medium spiny striatal neurons. Neuroscience. 2001;106(3):579–87. doi: 10.1016/s0306-4522(01)00297-4. PubMed PMID: 11591458.

67. Jin DZ, Xue B, Mao LM, Wang JQ. Metabotropic glutamate receptor 5 upregulates surface NMDA receptor expression in striatal neurons via CaMKII. Brain Res. 2015;1624:414–23. Epub 20150806. doi: 10.1016/j.brainres.2015.07.053. PubMed PMID: 26256252; PMCID: PMC4630094.

68. Chen HH, Liao PF, Chan MH. mGluR5 positive modulators both potentiate activation and restore inhibition in NMDA receptors by PKC dependent pathway. J Biomed Sci. 2011;18(1):19. Epub 20110222. doi: 10.1186/1423-0127-18-19. PubMed PMID: 21342491; PMCID: PMC3050796.

69. Chatterton JE, Awobuluyi M, Premkumar LS, Takahashi H, Talantova M, Shin Y, Cui J, Tu S, Sevarino KA, Nakanishi N, Tong G, Lipton SA, Zhang D. Excitatory glycine receptors containing the NR3 family of NMDA receptor subunits. Nature. 2002;415(6873):793–8. Epub 20020130. doi: 10.1038/nature715. PubMed PMID: 11823786.

70. West AB, Cowell RM, Daher JP, Moehle MS, Hinkle KM, Melrose HL, Standaert DG, Volpicelli-Daley LA. Differential LRRK2 expression in the cortex, striatum, and substantia nigra in transgenic and nontransgenic rodents. J Comp Neurol. 2014;522(11):2465–80. Epub 20140412. doi: 10.1002/cne.23583. PubMed PMID: 24633735; PMCID: PMC4076169.

71. Mandemakers W, Snellinx A, O’Neill MJ, de Strooper B. LRRK2 expression is enriched in the striosomal compartment of mouse striatum. Neurobiol Dis. 2012;48(3):582–93. Epub 20120729. doi: 10.1016/j.nbd.2012.07.017. PubMed PMID: 22850484.

72. Skelton PD, Tokars V, Parisiadou L. LRRK2 at Striatal Synapses: Cell-Type Specificity and Mechanistic Insights. Cells. 2022;11(1). Epub 20220105. doi: 10.3390/cells11010169. PubMed PMID: 35011731; PMCID: PMC8750662.

73. Parisiadou L, Yu J, Sgobio C, Xie C, Liu G, Sun L, Gu XL, Lin X, Crowley NA, Lovinger DM, Cai H. LRRK2 regulates synaptogenesis and dopamine receptor activation through modulation of PKA activity. Nat Neurosci. 2014;17(3):367–76. Epub 20140126. doi: 10.1038/nn.3636. PubMed PMID: 24464040; PMCID: PMC3989289.

74. Volta M, Beccano-Kelly DA, Paschall SA, Cataldi S, MacIsaac SE, Kuhlmann N, Kadgien CA, Tatarnikov I, Fox J, Khinda J, Mitchell E, Bergeron S, Melrose H, Farrer MJ, Milnerwood AJ. Initial elevations in glutamate and dopamine neurotransmission decline with age, as does exploratory behavior, in LRRK2 G2019S knock-in mice. Elife. 2017;6. Epub 20170920. doi: 10.7554/eLife.28377. PubMed PMID: 28930069; PMCID: PMC5633343.

75. Durakoglugil MS, Wasser CR, Wong CH, Pohlkamp T, Xian X, Lane-Donovan C, Fritschle K, Naestle L, Herz J. Reelin Regulates Neuronal Excitability through Striatal-Enriched Protein Tyrosine Phosphatase (STEP(61)) and Calcium Permeable AMPARs in an NMDAR-Dependent Manner. J Neurosci. 2021;41(35):7340–9. Epub 20210721. doi: 10.1523/JNEUROSCI.0388-21.2021. PubMed PMID: 34290083; PMCID: PMC8412985.

76. Feng X, Ippolito GC, Tian L, Wiehagen K, Oh S, Sambandam A, Willen J, Bunte RM, Maika SD, Harriss JV, Caton AJ, Bhandoola A, Tucker PW, Hu H. Foxp1 is an essential transcriptional regulator for the generation of quiescent naive T cells during thymocyte development. Blood. 2010;115(3):510–8. doi: 10.1182/blood-2009-07-232694. PubMed PMID: 19965654; PMCID: PMC2810984.

77. Ade KK, Wan Y, Chen M, Gloss B, Calakos N. An Improved BAC Transgenic Fluorescent Reporter Line for Sensitive and Specific Identification of Striatonigral Medium Spiny Neurons. Front Syst Neurosci. 2011;5:32. doi: 10.3389/fnsys.2011.00032. PubMed PMID: 21713123; PMCID: PMC3113108.

78. Gong S, Doughty M, Harbaugh CR, Cummins A, Hatten ME, Heintz N, Gerfen CR. Targeting Cre recombinase to specific neuron populations with bacterial artificial chromosome constructs. J Neurosci. 2007;27(37):9817–23. doi: 10.1523/JNEUROSCI.2707-07.2007. PubMed PMID: 17855595.

79. Agmon A, Connors BW. Thalamocortical responses of mouse somatosensory (barrel) cortex in vitro. Neuroscience. 1991;41(2-3):365–79. PubMed PMID: 1870696.

80. Hays SA, Huber KM, Gibson JR. Altered Neocortical Rhythmic Activity States in Fmr1 KO Mice Are Due to Enhanced mGluR5 Signaling and Involve Changes in Excitatory Circuitry. The Journal of Neuroscience. 2011;31(40):14223–34. doi: 10.1523/jneurosci.3157-11.2011.

81. Stroud H, Yang MG, Tsitohay YN, Davis CP, Sherman MA, Hrvatin S, Ling E, Greenberg ME. An Activity-Mediated Transition in Transcription in Early Postnatal Neurons. Neuron. 2020;107(5):874–90 e8. Epub 20200625. doi: 10.1016/j.neuron.2020.06.008. PubMed PMID: 32589877; PMCID: PMC7486250.

82. Andrews S. FastQC: A quality control tool for high throughput sequence data.

83. Fleming SJ, Chaffin MD, Arduini A, Akkad AD, Banks E, Marioni JC, Philippakis AA, Ellinor PT, Babadi M. Unsupervised removal of systematic background noise from droplet-based single-cell experiments using CellBender. Nat Methods. 2023. Epub 20230807. doi: 10.1038/s41592-023-01943-7. PubMed PMID: 37550580.

84. McGinnis CS, Murrow LM, Gartner ZJ. DoubletFinder: Doublet Detection in Single-Cell RNA Sequencing Data Using Artificial Nearest Neighbors. Cell Syst. 2019;8(4):329–37 e4. Epub 20190403. doi: 10.1016/j.cels.2019.03.003. PubMed PMID: 30954475; PMCID: PMC6853612.

85. Korsunsky I, Millard N, Fan J, Slowikowski K, Zhang F, Wei K, Baglaenko Y, Brenner M, Loh PR, Raychaudhuri S. Fast, sensitive and accurate integration of single-cell data with Harmony. Nat Methods. 2019;16(12):1289–96. Epub 20191118. doi: 10.1038/s41592-019-0619-0. PubMed PMID: 31740819; PMCID: PMC6884693.

86. Saunders A, Macosko EZ, Wysoker A, Goldman M, Krienen FM, de Rivera H, Bien E, Baum M, Bortolin L, Wang S, Goeva A, Nemesh J, Kamitaki N, Brumbaugh S, Kulp D, McCarroll SA. Molecular Diversity and Specializations among the Cells of the Adult Mouse Brain. Cell. 2018;174(4):1015–30 e16. doi: 10.1016/j.cell.2018.07.028. PubMed PMID: 30096299; PMCID: PMC6447408.

87. Robinson MD, McCarthy DJ, Smyth GK. edgeR: a Bioconductor package for differential expression analysis of digital gene expression data. Bioinformatics. 2010;26(1):139–40. Epub 20091111. doi: 10.1093/bioinformatics/btp616. PubMed PMID: 19910308; PMCID: PMC2796818.

88. Squair JW, Gautier M, Kathe C, Anderson MA, James ND, Hutson TH, Hudelle R, Qaiser T, Matson KJE, Barraud Q, Levine AJ, La Manno G, Skinnider MA, Courtine G. Confronting false discoveries in single-cell differential expression. Nat Commun. 2021;12(1):5692. Epub 20210928. doi: 10.1038/s41467-021-25960-2. PubMed PMID: 34584091; PMCID: PMC8479118.

89. Yu G, Wang LG, Han Y, He QY. clusterProfiler: an R package for comparing biological themes among gene clusters. OMICS. 2012;16(5):284–7. Epub 20120328. doi: 10.1089/omi.2011.0118. PubMed PMID: 22455463; PMCID: PMC3339379.

90. Supek F, Bosnjak M, Skunca N, Smuc T. REVIGO summarizes and visualizes long lists of gene ontology terms. PLoS One. 2011;6(7):e21800. Epub 20110718. doi: 10.1371/journal.pone.0021800. PubMed PMID: 21789182; PMCID: PMC3138752.

91. Koopmans F, van Nierop P, Andres-Alonso M, Byrnes A, Cijsouw T, Coba MP, Cornelisse LN, Farrell RJ, Goldschmidt HL, Howrigan DP, Hussain NK, Imig C, de Jong APH, Jung H, Kohansalnodehi M, Kramarz B, Lipstein N, Lovering RC, MacGillavry H, Mariano V, Mi H, Ninov M, Osumi-Sutherland D, Pielot R, Smalla KH, Tang H, Tashman K, Toonen RFG, Verpelli C, Reig-Viader R, Watanabe K, van Weering J, Achsel T, Ashrafi G, Asi N, Brown TC, De Camilli P, Feuermann M, Foulger RE, Gaudet P, Joglekar A, Kanellopoulos A, Malenka R, Nicoll RA, Pulido C, de Juan-Sanz J, Sheng M, Sudhof TC, Tilgner HU, Bagni C, Bayes A, Biederer T, Brose N, Chua JJE, Dieterich DC, Gundelfinger ED, Hoogenraad C, Huganir RL, Jahn R, Kaeser PS, Kim E, Kreutz MR, McPherson PS, Neale BM, O’Connor V, Posthuma D, Ryan TA, Sala C, Feng G, Hyman SE, Thomas PD, Smit AB, Verhage M. SynGO: An Evidence-Based, Expert-Curated Knowledge Base for the Synapse. Neuron. 2019;103(2):217–34 e4. Epub 20190603. doi: 10.1016/j.neuron.2019.05.002. PubMed PMID: 31171447; PMCID: PMC6764089.

92. Deacon RM. Assessing nest building in mice. Nat Protoc. 2006;1(3):1117–9. doi: 10.1038/nprot.2006.170. PubMed PMID: 17406392.

